# Persistent LPS insertion, spatial segregation and vesicle biogenesis drive growth-independent adaptation of the *Escherichia coli* outer membrane

**DOI:** 10.1101/2025.10.09.681453

**Authors:** Joe Nabarro, Natasha E. Hatton, Dmitri O. Pushkin, Martin A. Fascione, Christoph G. Baumann

## Abstract

The barrier and load-bearing functions of the Gram-negative bacterial outer membrane (OM) depend on ordered, dense packing of lipopolysaccharide (LPS), the major constituent of its outer leaflet. Despite its importance, LPS spatiotemporal dynamics and turnover mechanisms remain poorly understood. Existing models posit that LPS turnover only occurs via passive dilution during growth-dependent OM expansion, restricting adaptive LPS turnover, especially in nutrient-limited conditions. Here, using innovative pulse–chase LPS metabolic labelling techniques in combination with super-resolution microscopy, we demonstrate that *Escherichia coli* maintains OM homeostasis through continuous removal of pre-existing LPS and insertion of new LPS, even during stationary phase. We show that newly inserted LPS localises at discrete sites across the OM, remaining spatially segregated from pre-existing, background LPS. These observations challenge established OM organisational principles, suggesting an insertion-trapping mechanism can maintain LPS-rich clusters independent of a thermodynamically-driven phase separation. Time-lapse super-resolution imaging, biochemical assays, and nanoparticle tracking collectively reveal that OM vesicle (OMV) release mediates background LPS clearance. Our findings reveal bacteria have the capacity to remodel their surface architecture through OMV-mediated LPS turnover in a growth-independent manner. These insights redefine OM homeostasis and establish OMV biogenesis as fundamental to OM adaptation in Gram-negative bacteria.

## Introduction

The asymmetric outer membrane (OM) of Gram-negative bacteria provides an innate permeability barrier that protects the cell from toxic compounds, large antibiotics (typically >500 Da), and host immune defence molecules such as antimicrobial peptides and antibodies ^1–4^. In addition to this protective role, the OM contributes to the structural integrity of the bacterial cell envelope, providing rigidity and mechanical stability ^5,6^. These functions are critically dependent on OM molecular organisation, whose outer leaflet comprises a densely packed layer of lipopolysaccharide (LPS) molecules interspersed with a range of transmembrane β-barrel outer membrane proteins (OMPs). While OMPs enable selective permeability and impart key functionalities such as intercellular signalling and toxin secretion ^7,8^, it is LPS, as the major constituent of the OM outer leaflet, that intrinsically underpins the barrier and load-bearing functional roles of the OM ^4,9^. Notably, extreme lateral confinement of LPS in the OM, governed by a network of cooperative intermolecular interactions^10,11^, reinforces its rigidity and creates a highly-ordered glycolipid matrix restricting the lateral mobility of all OM components, which is essential for maintaining both permeability resistance and mechanical robustness ^11,12^.

Despite its accepted functional significance, remarkably little is known about how LPS organisation in the OM is maintained and the subsequent fate of LPS after it is inserted in the OM. Existing models largely assume that LPS remains in the OM indefinitely, diluted only by envelope area expansion during growth and partitioned during division ^10,13^. Thus, whether LPS can be turned over in a growth-independent manner or preferentially removed from the OM facilitating adaptive remodelling of the primary environmental interface in Gram-negative bacteria, has yet to be established. This represents a significant blind spot in our understanding of bacterial envelope homeostasis and adaptation, particularly in non-dividing or slow-growing cells in stationary phase occupying pathologically relevant nutrient-limited niches. These include for example, the uroepithelium, the viscous, hypoxic mucus of cystic fibrosis airways, and membrane-bound intracellular compartments within host cells ^14–16^. How OM integrity is preserved or actively maintained in non-growing cells found within these environments remains poorly understood.

Herein, through combined pulse-chase metabolic labelling experiments and super-resolution microscopy (SRM) we demonstrate that *Escherichia coli* cells are capable of actively turning over their OM LPS content in growth-arrested stationary phase, with outer membrane vesicle (OMV) biogenesis identified as a primary route for preferential removal of background LPS. We also re-examine the spatial distribution of LPS within the OM and present a revised hypothesis for OM organisation wherein outer leaflet architecture and organisation is governed by localised, stochastic insertion events and extremely restricted lateral diffusion, rather than thermodynamically-driven phase separation. Most notably our results reveal that (1) newly inserted LPS is delivered to discrete sites distributed across the OM; (2) once inserted, new LPS remains spatially segregated from background LPS; (3) LPS cluster size and frequency evolve over time with no evidence of patch coalescence and (4) LPS insertion and turnover continue during stationary phase, with turnover mediated by OMV release that preferentially removes older background LPS.

## Results

### Dual metabolic and fluorescent labelling reveals discrete colocalisation of LptD and newly inserted LPS in the *E. coli* outer membrane

To investigate the spatial dynamics of LPS insertion within the Gram-negative outer membrane (OM), we implemented a novel dual-labelling strategy in combination with two-colour SRM analysis enabling simultaneous, orthogonal detection of newly inserted LPS and its cognate OM transporter, LptD ^11,17^ ***(Fig. 1A, Extended Data Fig. S1)***. Newly synthesised LPS was metabolically tagged via incorporation of Kdo-N_3_ within its inner core oligosaccharide domain, while LptD was site-specifically labelled at residue position D600 (LptD*, ***Fig. 1A, Extended Data Fig. S1B and S1C*)** using genetic code expansion ^11,18,19^. This combination of bio-orthogonal handles permitted highly specific, simultaneous fluorescent labelling of newly inserted LPS and LptD* using copper-catalysed azide-alkyne cycloaddition (CuAAC) ‘*click’* reactions and complementarily functionalised fluorescent dyes (AF488 and AF647). By coupling this dual-labelling approach with two-colour direct stochastic optical reconstruction microscopy (dSTORM), we were able to visualise newly inserted LPS and LptD distributions across the OM surface of individual cells with nanometre-scale resolution ***(Fig. 1A and 1B, Extended Data Fig. S1A)*.**

**Figure 1:**
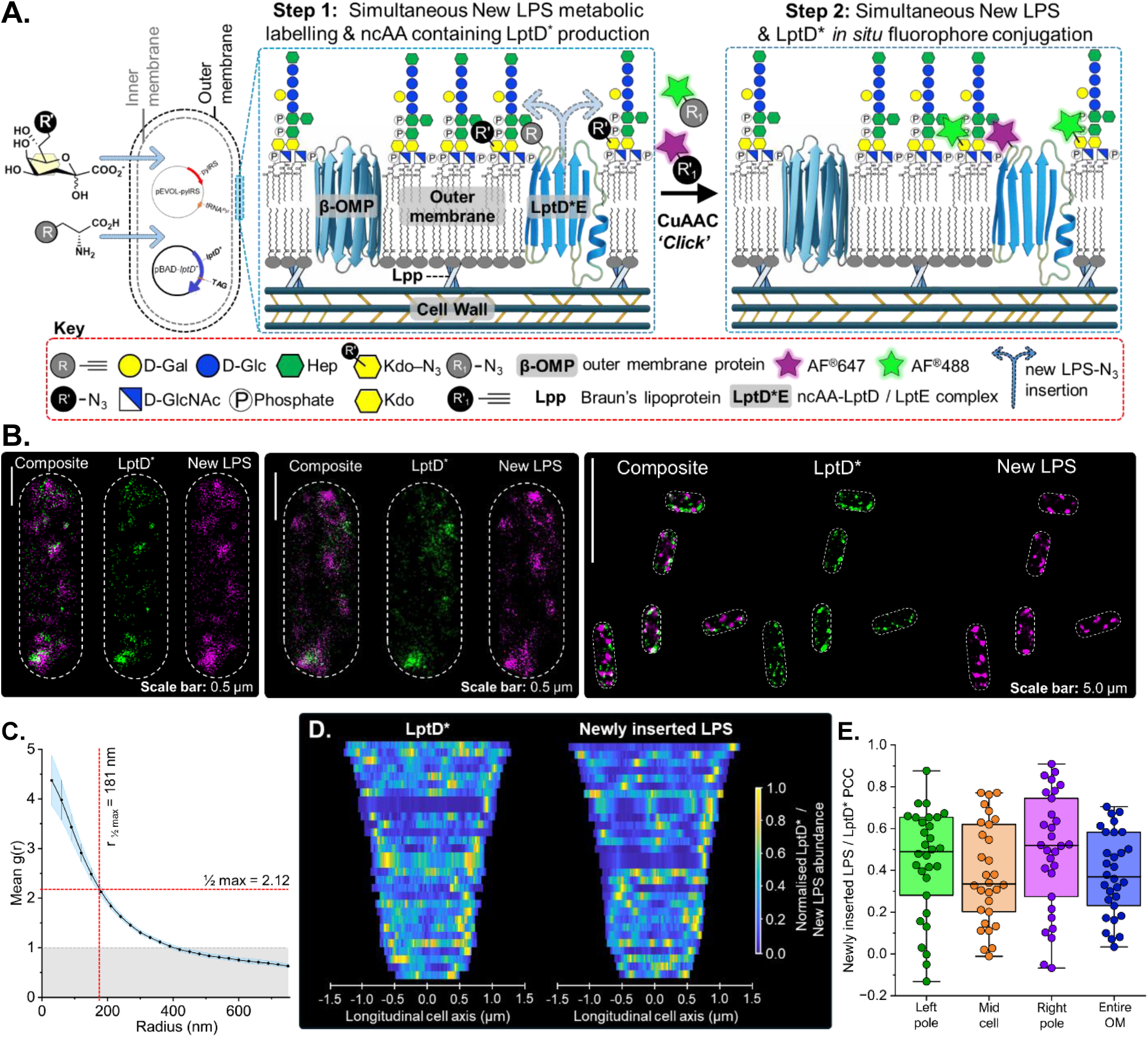
*In situ* dual bio-orthogonal labelling in combination with two-colour direct stochastic optical reconstruction microscopy (dSTORM) revealed LptD and newly inserted LPS colocalise at discrete sites across the *E. coli* outer membrane. **A. Schematic illustrating the dual LptD* / newly inserted LPS labelling strategy.** Newly synthesised LPS was metabolically labelled via Kdo-N_3_ incorporation into its inner core oligosaccharide domain, while LptD* was site-specifically labelled at position D600 using genetic code expansion to incorporate propargyl-L-Lysine. Simultaneous CuAAC ‘*click*’ chemistry enabled orthogonal fluorescent tagging of LptD* and newly inserted LPS using azide- and alkyne-functionalised dye pairs, respectively. Glycan symbols follow the SNFG convention. **B. Representative two-colour dSTORM single cell (left) and widefield (right) images showing the colocalised distributions of fluorescently labelled, newly inserted LPS and LptD* at discrete sites across the OM of *E. coli* cells.** Consistent, colocalisation of LptD* and newly inserted LPS was observed with new LPS being inserted at discrete sites. This resulted in the emergence of high density newly inserted LPS patches across the OM. **Single-cell scale bars:** 0.5 µm. **Widefield scale bars:** 5 µm. **C. Mean cross-correlation function (g*(r)*) calculated across 30 cells for newly inserted LPS and LptD\*** (N = 3, includes both dye combinations). Strong positive correlation at small radii indicated consistent colocalisation of newly inserted LPS and LptD* on sub-diffraction length scales. The r ½ max value (radius at which g(r) falls to half of its peak value) quantifies the typical colocalisation domain size at 181 nm, while the ½ max line (= 2.12) indicates the half-maximal g*(r)* value used to determine this radius. **D. Demographs of LptD and newly inserted LPS distribution along the longitudinal cell axis.** Heatmaps show normalised signal intensity of LptD* *(left)* and newly inserted LPS *(right)* along the longitudinal axis of individual *E. coli* cells (n = 30). Each horizontal row corresponds to a single cell, aligned by PCA and sorted by cell length. Localisation intensity is normalised within each cell to visualise relative enrichment patterns. While both LptD* and new LPS exhibit spatial confinement, their longitudional distribution varies across individual cells, with no consistent, stereotyped localisation pattern at the population level. This suggests LPS insertion occurs at discrete, random sites, coincident with LptD, and does not appreciable mix with older LPS in the OM. **E. Box plot showing distributions of Pearson correlation coefficient in different regions of individual *E. coli* cells and for the entire OM surface.** Consistent positive PCC values support LptD* / new LPS colocalisation (no significant statistical difference between distributions observed via Mann Whitney T-test, *p* values range from 0.07 – 0.97, median PCC values range from +0.34 – 0.52, see Extended Data Table S11 for all statistical parameters). Positive PCC distribution patterns were obtained irrespective of OM region analysed in agreement with PCC values obtained when colocalisation was calculated across entire OM surface. The colocalisation of newly inserted LPS and LptD* is consistent across the entire OM surface for all cells. Each data point represents the calculated PCC value for a single cell (n = 30, N = 3), with boxes representating interquatile ranges and the horizontal line representing the median PCC value.

Two-colour dSTORM revealed that both LptD and newly inserted LPS localised to discrete puncta scattered across the OM surface, rather than forming diffuse or uniform distributions ***(Fig. 1B, Extended Data Fig. S1A)***. These patches were consistently observed on cells irrespective of the dye combinations employed ***(Extended Data Fig. S1A and S1D)***, supporting the robustness of the dual-labelling and imaging strategy. Notably, these LPS-rich patches of varying intensity colocalised with LptD* signals, suggesting that LPS insertion may be spatially restricted, and involve LptD-incorporating sites with different LPS insertion rates ***(Fig. 1B, 1C and 1D)***, consistent with previous studies visualising new LPS insertion in the OM ^10,12^. The distribution of LptD–LPS colocalisation sites across the OM indicate that LPS insertion occurs at discrete locations over the entire OM ***(Fig. 1D and 1E)***, rather than being confined to specific regions of the OM such as the division septum as is the case for BAM- mediated OMP insertion ^20^.

To quantify colocalisation between newly inserted LPS and LptD, we performed pairwise cross-correlation analysis across 30 cells from 3 biological replicates. The resulting mean cross-correlation function, g*(r),* showed a clear peak at short radii, confirming sub- diffraction co-enrichment of the two signals ***(Fig. 1C, Extended Data Fig. S1D and S1E)***. The r₁/₂ max value, representing the radius at which g*(r)* drops to half of its peak height, was 181 nm, indicating the typical size of colocalised domains, which is consistent with previous LPS patch and confinement diameter characterisations ^10–12,21^. The ½ max line value (g*(r)* = 2.12) further supports a strong local enrichment of new LPS near LptD, significantly above the random expectation of 1.

To investigate whether LPS insertion follows any stereotyped spatial pattern equivalent to BAM-mediated OMP insertion in the OM ^20^, we also generated demographs of LptD and newly inserted LPS along the longitudinal axis of individual cells (n = 30), aligning and sorting cells by length following principle component analysis (PCA)-based axis normalisation ***(Fig. 1D)***. While both signals were spatially confined, they exhibited heterogeneous distributions between cells with no consistent macrodomain localisation or polar enrichment. Instead, puncta appeared distributed yet spatially constrained, suggesting that LPS insertion occurs at discrete but stochastically positioned sites. These demographs also show that not all LptD*- dense regions in the OM were accompanied by equivalent dense regions of newly inserted LPS suggesting not all LptDE complexes are engaged in LPS insertion at once, consistent with the transient nature of some Lpt LPS translocon bridges that form between the sites of LPS production at the inner membrane and its insertion into the OM ^10,20,22–24^.

We further quantified LptD*–LPS spatial relationships using their associated Pearson correlation coefficient (PCC) across the full OM, and within discrete polar and mid-cell regions of individual cells ***(Fig. 1E)***. Box plots revealed consistently positive PCC values across all regions, with no statistically significant differences between the regional PCC values (Mann– Whitney U test, p > 0.05, ***Extended Data Table S11*)**, indicating that the mechanism of LPS insertion and its spatial relationship with LptD is maintained across the OM surface ***(Fig. 1E)***, consistent with the results of previous studies ^10,12^. Results showed that LPS insertion was not concentrated within specific regions of the OM, but instead takes place in a stochastic, dispersed manner across the OM.

### Spatial segregation of background and newly inserted LPS persists over time

To further investigate the spatiotemporal dynamics of LPS insertion and specific LPS subpopulations within the OM of *E. coli*, we developed a novel pulse–chase metabolic labelling strategy utilising differentially functionalised Kdo sugar analogues that we combined with two-colour dSTORM. A schematic overview of this approach is presented in ***Figure 2A***. Preliminary control experiments were performed to verify that both Kdo-analogues are incorporated in the inner core oligosachride domain of *E. coli* K12 LPS and enable equivalent levels of *in situ* fluorescent labelling ***(Extended Data Fig. S2A and S2C)***. To probe spatiotemporal LPS dynamics, cells were pulse-labelled with a Kdo-alkyne probe to generate what we designated as the pre-existing or “background” LPS population ^25^ ***(Extended Data Fig. 2A)***. After thorough washing to remove residual Kdo-alkyne, the cells were chased with an azide-functionalised Kdo analogue (Kdo-N_3_) to specifically label newly synthesised LPS molecules. Differential fluorescent labelling using CuAAC and distinct azide and alkyne functionalised fluorescent dyes enabled simultaneous visualisation of both LPS subpopulations in separate fluorescence channels. Two-colour 2D dSTORM imaging was performed on aliquots of the growing cells isolated at 15 min intervals over a two-hour period which had been dual-labelled with the functionalised dyes to facilitate precise spatiotemporal tracking of the LPS subpopulations at the single-cell level. Representative dSTORM images **(*Fig. 2B and 2C)*** illustrate the spatiotemporal evolution of the two LPS subpopulations. In agreement with results presented in ***Figure 1***, newly inserted LPS appeared predominantly as discrete, punctate clusters across the OM and had minimal overlap with pre-existing background LPS patches **(*Fig. 2B and 2C)***. Identical patterning was observed irrespective of dye selection ***(Fig. 2B and 2C, Extended Data Fig. S1D)***. The discrete spatial arrangement of background and newly inserted LPS subpopulations in the OM of *E. coli* cells also persisted throughout the entire experimental time course, strongly indicating that newly inserted and background LPS containing patches remain largely segregated, with limited colocalisation or mixing even as total membrane coverage by newly inserted LPS increases.

**Figure 2.**
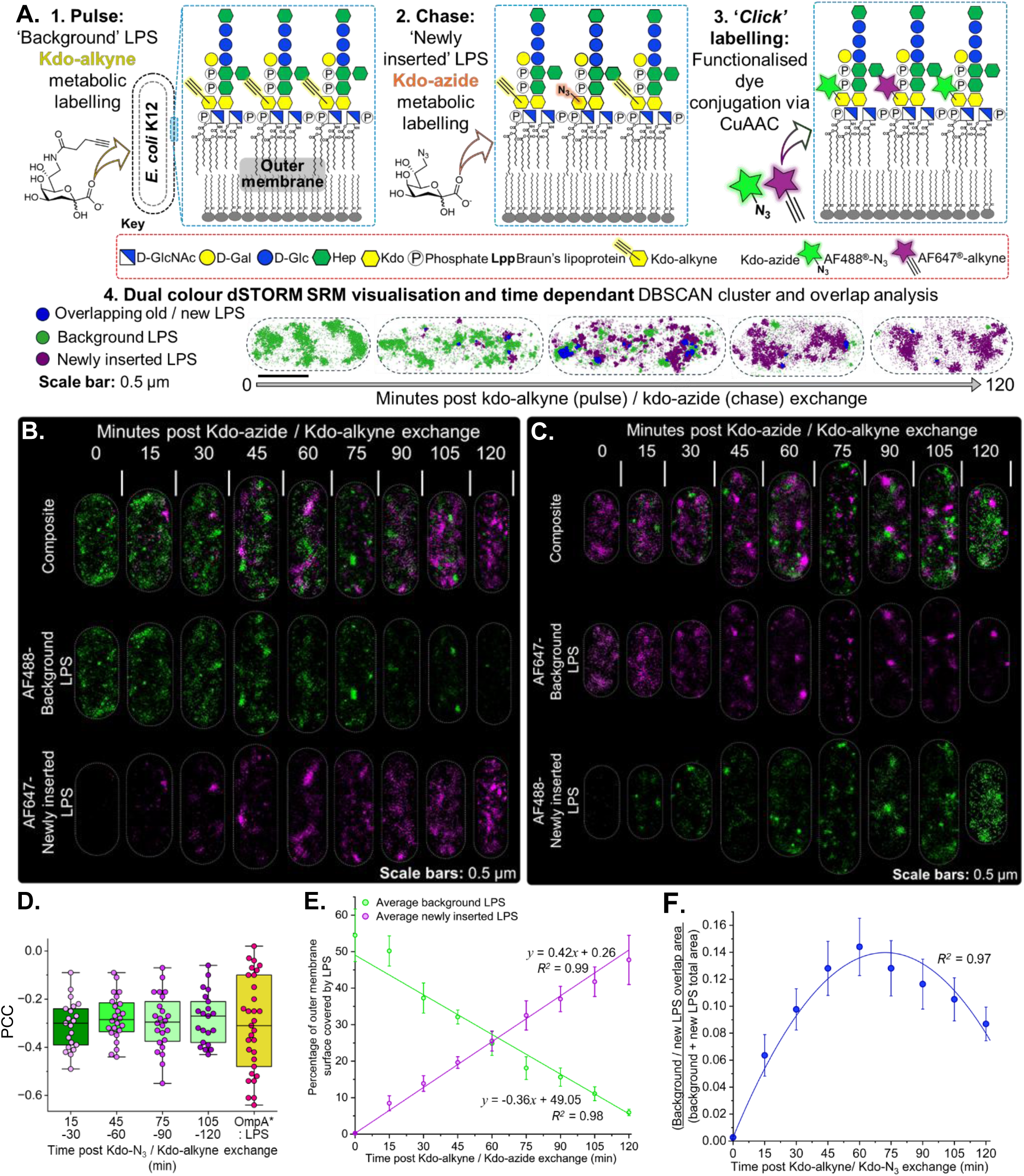
Kdo-alkyne / Kdo-N_3_ pulse–chase experiments done in combination with two-colour 2D dSTORM SRM imaging demonstrated newly inserted LPS and background LPS domains remain discrete in the OM of *E. coli.* This clustered organisation in the OM is likely generated by stochastic LPS insertion rates and maintained by the very restricted lateral diffusion of all OM components. **A. Schematic overview of the experimental strategy to visualise LPS dynamics in live *E. coli.*** A Kdo-alkyne probe **(1)** is first incorporated into the inner core oligosaccharide domain of background LPS. After removal of residual Kdo-alkyne, cells presenting background Kdo-alkyne-containing LPS at their surfaces are pulsed with a Kdo-N_3_ probe **(2)**, which is similarly incorporated into newly synthesised LPS. Cells were removed at 15 min intervals post Kdo-alkyne removal / Kdo-N_3_ addition after which background and newly inserted LPS were differentially fluorescently labelled via CuAAC with azide- (background) or alkyne- (newly inserted) functionalised fluorescent dyes **(3)**. Two-colour dSTORM SRM imaging was used to visualise the evolution of OM regions containing background and newly inserted LPS across the sampling time points on a cell-by-cell basis. Subsequent DBSCAN co-cluster analysis was used to quantify the spatiotemporal patterning of background *(green)* and newly inserted *(purple)* LPS in the OM of *E. coli* cells **(4)**. Mixed LPS patches including both background (old) and newly inserted LPS are highlighted in blue. Glycan symbols follow the SNFG convention. **Scale bars:** 0.5 µm **B. Representative two-colour 2D dSTORM images from Kdo-alkyne / Kdo-N_3_ pulse–chase experiments visualising the evolution of background LPS** *(Alexa Fluor 488 (AF488); green; middle row)* and newly inserted LPS *(Alexa Fluor 647 (AF647); magenta; bottom)* regions in the OM of *E. coli* cells at 15 min intervals post Kdo-analogue exchange over a period of 2 hours. Newly inserted LPS appeared in discrete foci that usually did not overlap with the existing background LPS patches. **Scale bars:** 0.5 µm. **C. Representative two-colour 2D dSTORM images from Kdo-alkyne/Kdo-N_3_ pulse–chase experiments visualising background LPS** *(AF647; magenta; middle row)* and newly inserted LPS *(AF488; green; bottom)* in the OM of *E. coli* cells at 15 min intervals post Kdo-analogue exchange over a period of 2 hours. These replicate two-colour SRM experiments used different dyes to label newly inserted LPS and LptD*, and the results obtained were identical to those presented in panel B. **Scale bars:** 0.5 µm **D. Box plots showing the Pearson correlation coefficient values calculated for background and newly inserted LPS as a function of time.** The consistently negative PCC values indicated that pre-existing background LPS and newly inserted LPS remain spatially segregated throughout the experiment. These anticorrelated values are comparable to those observed between OmpA* and LPS-rich patches *(yellow box)*, suggesting that the observed compartmentalisation of the two LPS populations does not represent phase separation. It instead reflects the distinct organisation of the *E. coli* OM which is defined by restricted lateral diffusion and discrete non-overlapping sites where OM components (*i.e*. OMPs and LPS) are inserted. **E. Quantification of outer membrane coverage and spatial overlap between background and newly inserted LPS clusters.** Percentage of OM area occupied by DBSCAN-defined background *(green)* and newly inserted *(magenta)* LPS clusters over a 120 min time course. Values represent mean ± standard error from individual cells (n ≥ 10 per timepoint). Linear regression of these data sets showed the loss of background LPS and accumulation of newly inserted LPS occurred at very similar reciprocal rates. **F. Normalised spatial overlap between background and newly inserted LPS clusters, calculated as background and newly inserted overlap area / (background + newly inserted LPS combined total area).** Overlap remained low throughout, peaking at 60 min before decreasing again, and was consistent with continued spatial segregation despite the close proximity of the two LPS populations in the OM. The strong adherence of the overlap data to a second-order polynomial fit (*R²* > 0.95) suggested that the formation of significant overlap is prevented by the aforementioned distinct organisation of the *E. coli* OM. This supports a model in which newly inserted LPS accumulates in domains that remain spatially segregated from old LPS clusters despite increasing membrane coverage by the former. Error bars represent standard error.

PCC values calculated from dSTORM data outputs revealed consistently negative values ***(Fig. 2D)***, confirming sustained spatial segregation between background and newly inserted LPS. These anti-correlation values were comparable to those we calculated for labelled OmpA*-rich domains and LPS patches in a previous study^11^, confirming that the observed LPS compartmentalisation was not driven by phase separation. Instead, our data support a distinct organisational model based on a stochastic distribution of LPS insertion sites with variable insertion rates, combined with highly restricted lateral diffusion within the OM.

To rigorously quantify the spatial relationships between background and newly inserted LPS domains, we applied a density-based clustering non-parametric algorithm (DBSCAN) to our dSTORM-derived datasets ^26^ ***(Extended Data Fig. S2E)***. Quantitative analysis of OM coverage by DBSCAN defined LPS clusters further reinforced our findings **(*Fig. 2E and 2F)***. Background LPS cluster coverage gradually decreased over the 120 min time course, concomitant with a reciprocal increase in coverage by newly inserted LPS clusters ***(Fig. 2E, Extended Data Fig. S2D)***. Spatial overlap between these two LPS populations remained consistently low, reaching only a modest transient peak at approximately 60 min before declining ***(Fig. 2F, Extended Data Fig. S2E*)**. Normalised spatial overlap, defined as the ratio of overlapping area to the total combined area of both LPS populations, closely fitted a second-order polynomial trajectory (*R²* > 0.95). This pattern is biologically meaningful; rather than increasing linearly or asymptotically with time, the overlap peaked transiently and then declined. Such behaviour suggests that LPS cluster merging is not a diffusive process driven by crowding, but instead reflects a mechanism in which lateral segregation of newly inserted and background LPS-rich OM domains arises and is maintained due to the highly restricted lateral diffusion in the OM. Collectively, these results robustly demonstrate that the spatial segregation of newly inserted and background LPS domains can persist over time. Therefore, once inserted into the OM, LPS molecules are effectively trapped in place with minimal lateral diffusion, consistent with prior findings that LPS lateral mobility in the OM is tightly confined ^10–12^, which we define as an ‘insertion-trapping’ mechanism.

### Kdo-N_3_ pulse–chase experiments reveal continuous insertion of new LPS and removal of background LPS in a growth-independent manner

Having shown that newly inserted LPS exists in discrete, spatially segregated patches located across the entire OM (***Figs. 1 and 2)***, we adapted our pulse–chase methodology to enable us to study the kinetics of LPS incorporation and turnover during exponential phase growth of the *E. coli* population and its transition to stationary phase. We performed complementary single-colour pulse–chase experiments using Kdo-N_3_ and native Kdo combined with single-colour dSTORM imaging. One experiment tracked rates of new LPS insertion into the OM over time on a cell-by-cell basis, whilst the other monitored loss of pre-existing, background LPS from the OM of individual *E. coli* cells. To track the accumulation of newly inserted LPS over time, cells were initially cultured under standard conditions and then transferred to defined, liquid CDM containing excess Kdo-N_3_ once they had reached the mid-exponential phase of growth (OD_600_ = 0.5), thereby initiating metabolic labelling of newly synthesised LPS with the azido-functionalised Kdo analogue. Samples were removed from the culture at 30 min intervals over a three hour time course. Newly inserted LPS was fluorescently labeled via CuAAC in these samples and analysed on a cell-by-cell basis via dSTORM. By maintaining standardised imaging parameters over an attenuated number of imaging frames (to minimise photobleaching) we were able to use the density of discrete fluorescent LPS localisations per OM area as a proxy for the relative abundance of newly inserted LPS present in the OM at each time point.

Single-cell level dSTORM imaging over a 210 min chase period ***(Fig. 3A)*** again revealed ongoing incorporation of newly synthesized LPS at discrete punctate sites distributed across the OM. Notably, despite the culture having transitioned into stationary phase ***(Fig. 3B)*** these insertion events continued and insertion foci remained spatially discrete rather than becoming more diffuse over time. Subsequent quantification of the number of discrete fluorescent LPS localisations (n ≥ 30, N = 3 for each time point – ***Extended Data Table S17***) revealed a continuous increase in newly inserted LPS throughout the experiment ***(Fig. 3B, Extended Data Fig. S3B)***. Notably, the rate of new LPS accumulation per cell did not slow, even between 150–210 min, when cell growth had markedly slowed as result of the culture entering stationary phase **(*Fig. 3B, Extended Data Fig. S3B-S3D)***. This indicates that LPS insertion rates in the OM are maintained beyond the point of cell elongation, suggesting that LPS insertion in stationary phase *E. coli* cells becomes decoupled from rates of cell growth and division.

**Figure 3:**
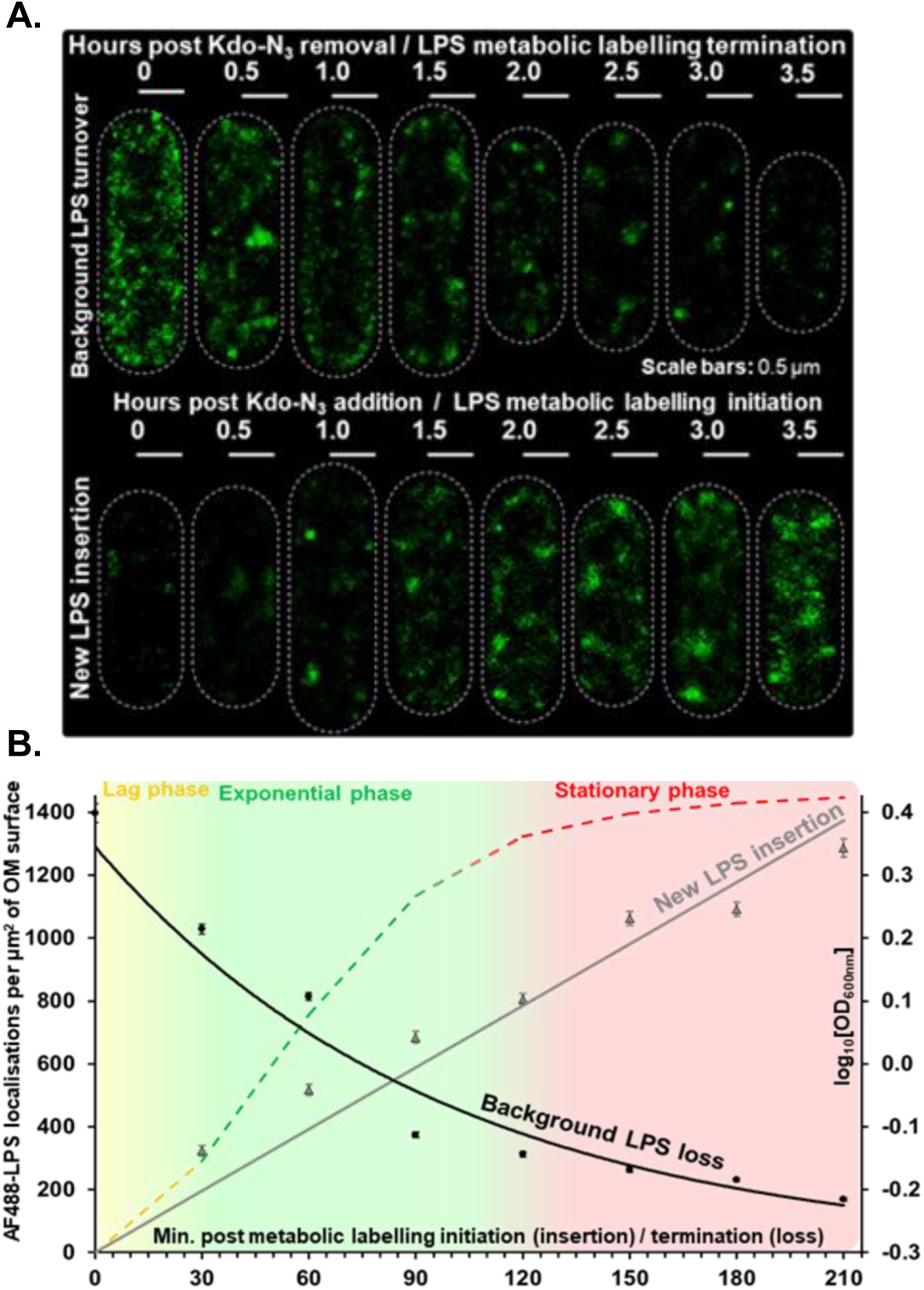
Kdo-N_3_ pulse–chase dSTORM experiments reveal ongoing LPS insertion and turnover in stationary phase. **A. Representative dSTORM images showing sites of background LPS *(top)* and new LPS insertion *(bottom)* in the OM of single-cells over time.** New LPS insertion events (bottom) appear as bright foci at discrete locations across the OM, while the background LPS (top) remains in discrete patches that appear to diminish in size, frequency and density over time. These images confirm that new LPS is inserted at discrete sites across the OM, and that old LPS is not sequestered into specific regions of the OM during cell elongation. **Scale bars:** 0.5 µm **B. Quantification of the relative rates of background LPS turnover and new LPS insertion on a per cell basis (n ≥ 30 cells per time point, N = 3 biological replicates for each time point).** The average number of discrete AF488–LPS localisations in the OM per cell was used as a proxy for the relative amount of newly inserted or background LPS in the OM. Both the insertion of new LPS and the loss of background LPS continued throughout the 210 min chase period, even as the rate of cell growth (monitored as OD_600_ of the culture) slowed significantly as the culture entered stationary phase.

In a second set of experiments, we monitored the loss of old LPS from the OM. Cells were pre-labeled extensively with Kdo-N_3_ (labelling pre-existing ‘background’ LPS), then transferred to fresh medium containing native Kdo. The decline of the fluorescent signal from background LPS was similarly quantified by single-cell SRM imaging ***(Fig. 3B, Extended Data Fig. S3A, Extended Data Table S16)***, but also by bulk biochemical analysis of extracted LPS ***(Fig. 6D, Extended Data Fig. S6A and S6C)***. At the single-cell level, dSTORM imaging revealed a progressive decline in fluorescently labelled LPS density across the OM ***(Fig. 3B, Extended Data Fig. S3A)***, while at the bulk-population level fluorescence analysis of Tricine SDS-PAGE gels showed steady loss of (fluorescent) background LPS over time ***(Fig. 6, Extended Data Fig. S6A and S6C)***. In both single-cell dSTORM and bulk biochemical assays the LPS signal reduction persisted during stationary phase, despite cell elongation and division having slowed significantly ***(Fig. 3B, Extended Data Fig. S3A and S6A)***. These results show that background LPS is not just diluted in the OM of multiplying cells, as previously postulated ^10^, but also preferentially removed from the OM in growth-arrested, non-dividing cells in stationary phase.

### LPS-rich clusters remain spatially discrete and heterogeneously distributed, inconsistent with phase separation in the OM

To investigate the observed nanoscopic clustered organisation in the OM of *E. coli* cells, we analysed the spatiotemporal behaviour of background and newly inserted LPS by projecting single-molecule localisations from the dSTORM pulse–chase onto a half-rod cylindrical model of each cell ***(Fig. 4A)***, which enabled the identification of LPS clusters using a DBSCAN algorithm ^27,28^. Organisation in the OM is currently proposed to be governed by phase separation ^21,29^, in which classical models dictate smaller domains should coalesce into larger ones over time ^27,28,30–32^. However, in our analysis of newly inserted LPS ***(Fig. 4B)***, the number of clusters per cell increased steadily over the 210 min experiment, while the total area covered by LPS also expands. Crucially, the mean cluster size plateaued after 90 min. This result, together with earlier observations that new LPS does not merge with pre-existing domains ***(Fig. 2)***, indicates that LPS incorporation results in stable, spatially discrete patches that persist over time, contrary to the equilibrium-driven coarsening expected during phase separation ^30^. Analysis of background LPS clusters reinforced this conclusion, with cluster number, size, and density decreasing over the same period ***(Fig. 4C)***. Rather than fusing into larger domains, these pre-existing LPS-rich patches gradually dissipated, consistent with their dilution during OM expansion in growing and multiplying exponential phase cells. No evidence was observed of smaller domains vanishing as larger ones grew in contrast to results predicted by Ostwald ripening or domain fusion ^32^. These observations are incompatible with a phase-separated system, where ageing domains generally expand and grow with time as a result of smaller domain coalescence ^33^.

**Figure 4.**
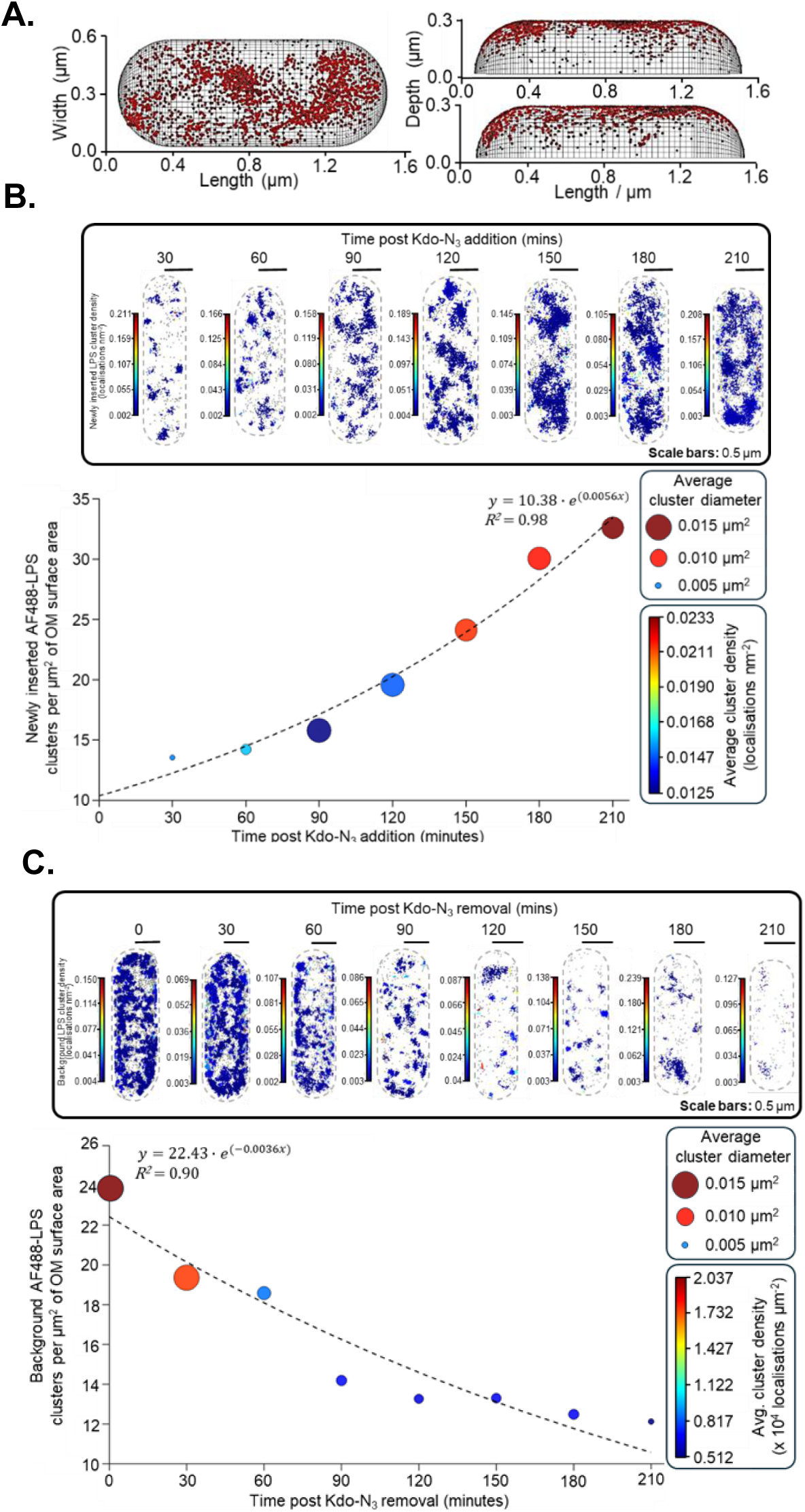
Cluster analysis of single-colour dSTORM data revealed that LPS-rich domains remain spatially discrete and do not coarsen over time in the OM of *E. coli* cells. **A. Schematic illustrating 3D curvature correction.** Single-molecule localisations from 2D dSTORM images were projected onto a half-rod cylindrical model of each *E. coli* cell to correct for surface curvature and enable more accurate membrane-wide analysis of clustering. **B. Newly inserted LPS cluster analysis.** Representative cluster maps (top, scale bars: 0.5 µm) and quantification (bottom) showed that newly inserted Kdo-N_3_-containing, AF488-labelled LPS accumulates in discrete patches whose number, density and total area increase over time. However, the mean cluster size plateaus after ∼90 min, and clusters do not merge. Each point on the graph represents the mean cluster value per cell; colour and size correspond to average cluster density and diameter, respectively. The exponential growth of cluster number (*R²* = 0.98) was consistent with continued LPS insertion, even in stationary phase. These results indicate that neither background nor newly inserted LPS domains undergo coarsening or coalescence over time, which indicates that LPS cluster formation does not involve membrane phase separation **C. Background LPS cluster analysis.** Representative cluster maps (top, scale bars: 0.5 µm) and quantification (bottom) show that background Kdo-N_3_-containing, AF488-labelled LPS clusters steadily decreased in abundance, size and density over the 210 min time course. The exponential decay of the cluster number (*R²* = 0.90) suggested dilution of background LPS in the OM during exponential phase growth followed by preferential removal during stationary phase with no evidence for domain coalescence.

To examine how LPS-rich clusters were spatially distributed within the OM, we also categorised clusters from individual cells into left pole, mid-cell, and right pole regions based on their position along the longitudinal cell axis ***(Extended Data Fig. S4)***. Using global area scaling for visualisation, per-cell plots were generated to examine clustering patterns over time for both background and newly inserted LPS ***(Extended Data Fig. S4G and S4H)***. Quantitative analysis showed no significant differences in cluster number, density, or size between polar and mid-cell regions at any timepoint. These spatial patterns remained stable throughout both LPS turnover and insertion experimental time courses, without evidence of domain coalescence or regional convergence ***(Extended Data Fig. S4A-S4F)***. These findings further support the conclusion that LPS does not laterally diffuse within the OM at a sufficient rate to homogenise membrane organisation ^20^.

Together, these results demonstrate that the OM of *E. coli* does not behave as a phase- separated system. Instead, it is organised as a kinetically-trapped mosaic of spatially segregated domains. Old LPS is progressively lost without domain fusion, and new LPS accumulates in stable patches, supporting a model of hierarchical insertion and spatial confinement ^20,21,34^. In this model, LPS-rich domains emerge and persist because newly inserted LPS is effectively frozen in place, rather than laterally diffusing and coalescing into larger equilibrated domains.

### Live-cell 3D-SIM² imaging reveals that background LPS turnover does not undergo binary partitioning or internal cytoplasmic recycling

To identify the potential mechanisms responsible for LPS turnover in the OM of *E. coli*, we tested whether background LPS was recycled internally, or partitioned directionally during division, analogous to the behaviour of OMPs during cell elongation ^20^. If LPS was being partitioned in the same way as OMPs, one would expect its removal from the OM simply as a consequence of division (with old LPS being segregated to the poles and diluted out in daughter cells), without requiring any additional turnover mechanism such as OM shedding. Alternatively, old LPS could conceivably be internalised into the cell and recycled internally (for example, via envelope-spanning complexes that transport LPS back across the periplasm)^35^. However, previous published studies have suggested that buildup of LPS in the inner membrane is toxic to cells, making an active recycling pathway unlikely ^36–41^. To investigate these possibilities, we performed time-lapse 3D-structured illumination super-resolution microscopy (SIM²) to track background LPS dynamics in the OM of growing *E. coli* cells after fluorescently labelling background Kdo-N_3_-containing LPS using copper-free ‘*click*’ (SPAAC)^42^. After mounting on M9 CDM / agarose pads, viable cells with AF-488 labelled LPS were imaged over multiple generations at 37 °C with 3D-SIM^2^ enabling optical sectioning in the z-direction, and isolation of background, fluorescent LPS signal in the upper cell envelope, cytoplasm and lower cell envelope ***(Fig. 5, Extended Data Fig. S5)***. Rates of cell elongaton and division were assessed ***(Extended Data Fig. S5D)*** to ensure that *E. coli* growth was unaffected by labelling or electromagentic radiation exposure (resulting from excitation laser). We found no evidence for intracellular accumulation or binary partitioning of background LPS during cell growth. Instead, fluorescently labelled LPS remained in discrete patches across the OM and was inherited uniformly by daughter cells without polar enrichment ***(Fig. 5, Extended Data Fig. S5).*** These live-cell observations are consistent with our fixed-cell dSTORM data ***(Figs. 1 – 3)***. Together, they confirm that LPS turnover does not occur via binary partitioning (as seen for OMPs), nor via internal recycling ^20^.

**Figure 5:**
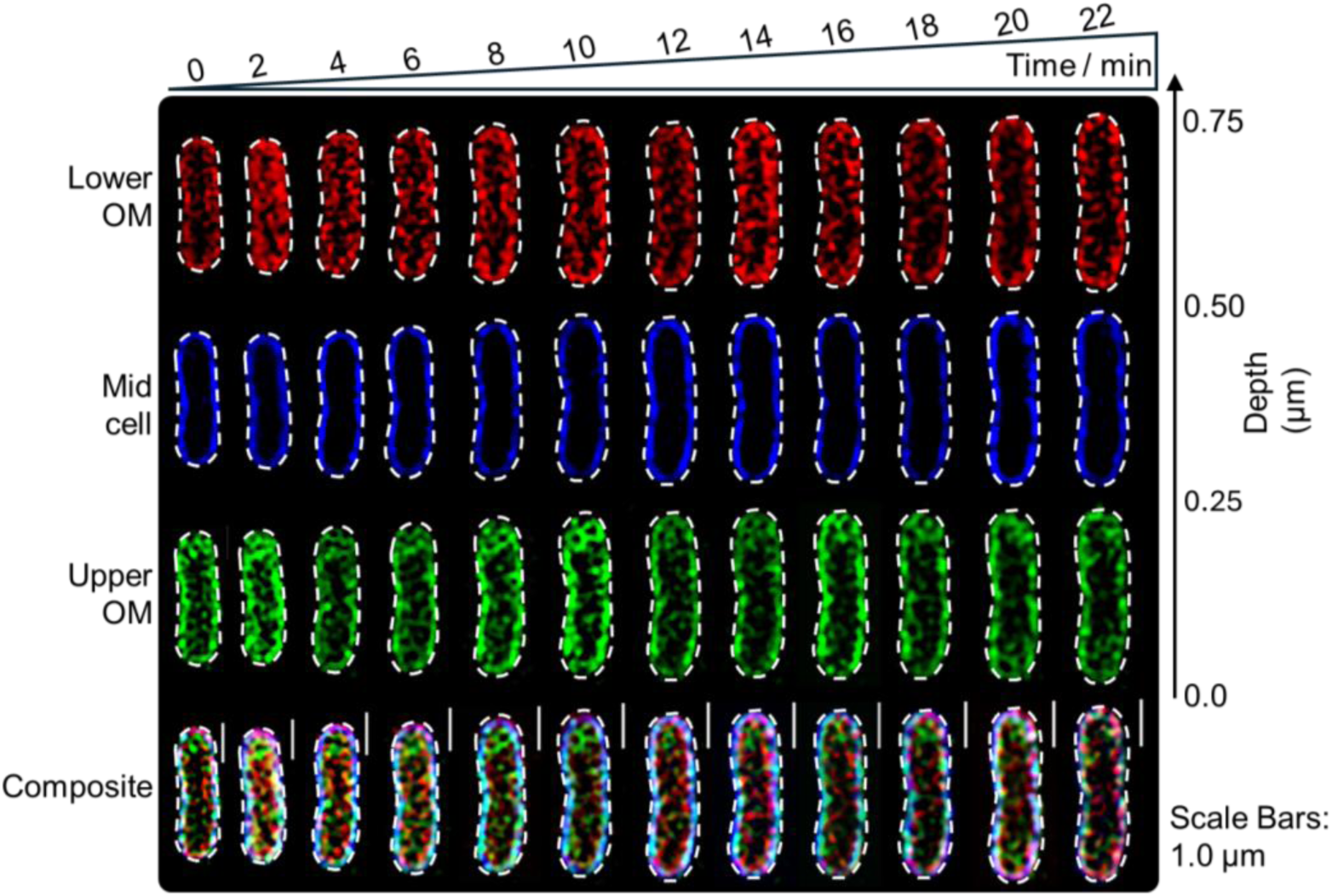
Live-cell 3D-SIM² imaging showed that background LPS was not partitioned to cell poles or internalised during cell growth. No accumulation of fluorescently labelled LPS was observed within the polar regions of the OM in single cells (see optical sections for lower OM and upper OM). The optical section of the mid-cell region (*i.e*. inside the cytoplasm) did not show any accumulation of fluorescence signal. Background LPS distributions are maintained in the OM during cell elongation indicating that LPS was not undergoing binary partitioning or being internalised during cell growth. **Scale bars:** 1.0 µm.

### Outer membrane vesicle biogenesis contributes to LPS turnover in stationary phase *E. coli* cells

Our data shows that in stationary phase, new LPS insertion and background LPS turnover rates in the OM are maintained despite significant slowing in the rates of cell elongation and division ***(Figs. 2–4)***. Therefore, non-dividing cells must actively remove LPS to maintain OM homeostasis and permit continual LPS insertion in the absence of cell elongation. Having excluded LPS internalisation, we investigated whether outer membrane vesicle (OMV) biogenesis might mediate LPS turnover in stationary-phase cells ^43–46^.

Using live-cell 3D-SIM², we captured OMV formation events in real-time ***(Fig. 6A)***. Time-lapse imaging revealed dynamic OM blebs that progressively expanded and subsequently detached as discrete vesicles, carrying labelled, background LPS away from the cell surface and thus creating ‘space’ in the OM for new LPS insertion. ***Figure 6A*** shows a representative 3D-SIM² depth-coded projection of OMV blebbing and release in a live, stationary phase *E. coli* cell over a 10 min time period. This direct visualisation confirms that stationary phase wild-type *E. coli* cells can harness OMV release to mediate LPS removal from the OM.

**Figure 6:**
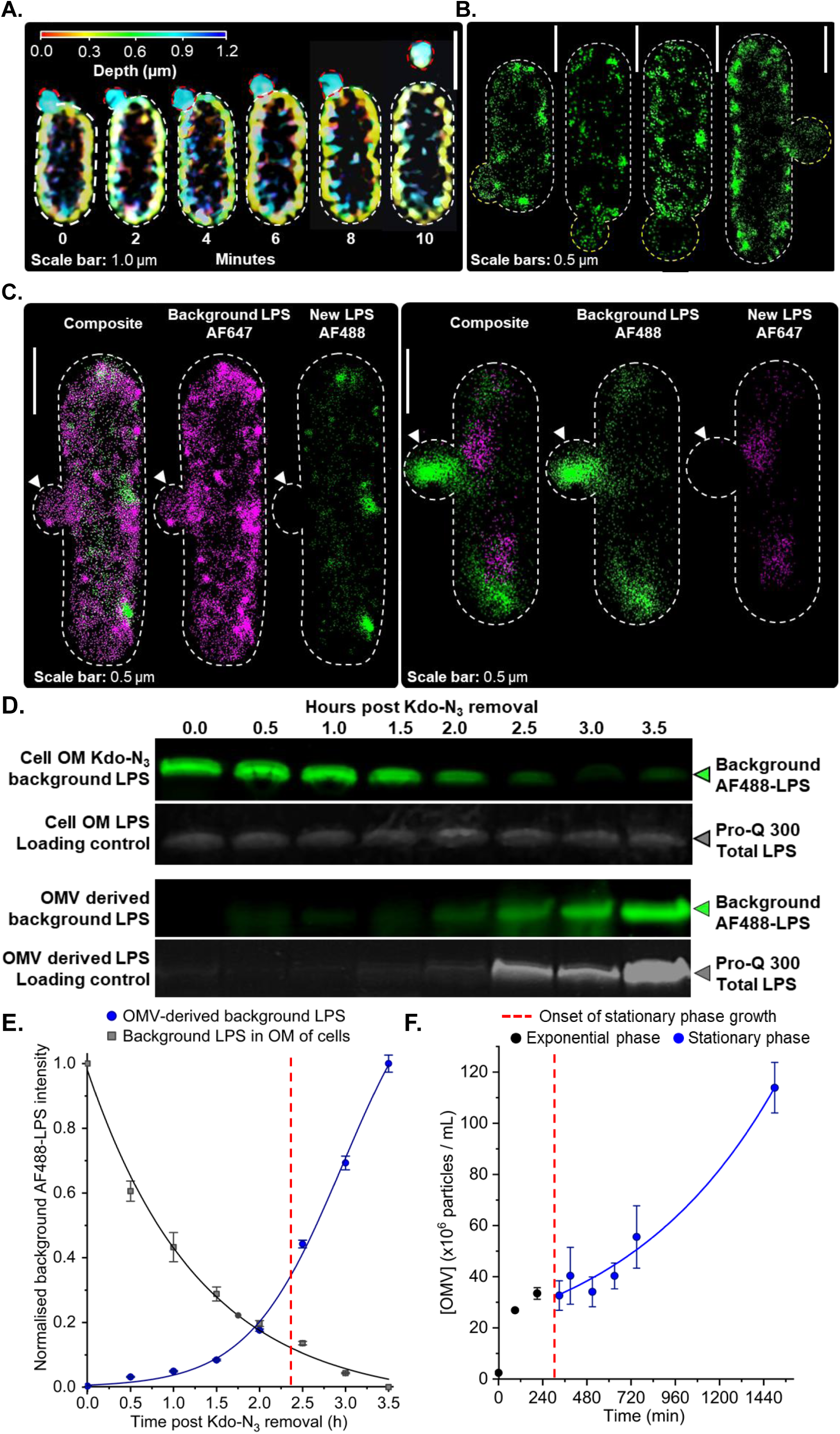
Preferential OMV-mediated clearance of background LPS from the OM of non-elongating, stationary phase *E. coli* cells. **A. Live-cell 3D-SIM² visualisation of OMV formation in a stationary phase *E. coli* cell presenting fluorescently labelled background LPS.** A depth-coded 3D projection from time-lapse imaging shows a AF488-LPS-rich OMV budding from the cell surface and being released over a 10 min period. **Scale bars:** 1.0 µm **B. dSTORM images of stationary phase *E. coli* cells in the process of background LPS shedding via OMV biogenesis. Scale bars:** 0.5 µm **C. Background LPS was enriched in OMVs produced by stationary phase *E. coli* cells.** Representative dual-colour dSTORM images showing the spatial distribution of background (pre-existing) and newly inserted LPS in the OM of *E. coli* cells undergoing OMV formation. Background and newly inserted LPS were metabolically labelled with Kdo-alkyne and Kdo-azide, respectively, using an *in vivo* pulse–chase method which enabled selective fluorescent labelling by CuAAC with azide- or alkyne-functionalised fluorophores. Background LPS (magenta; left set of images or green; right set of images) was enriched in the vesicle-like membrane protrusions (white arrows), while newly inserted LPS (green; left set of images or magenta; right set of images) appears to be excluded from these protrusions. Images indicate spatial segregation of background and new LPS in the OM, with preferential incorporation of background LPS into membrane blebs in a dye-independent manner. This observation suggests stationary phase cells can employ OMV blebbing to facilitate removal of background LPS in the absence of cell elongation and division. **Scale bars:** 0.5 μm. **D. Biochemical characterisation of background LPS levels in the OM of intact cells and released OMVs as a function of time by Tricine SDS-PAGE.** The material from an equivalent number of cells was loaded in each lane for the LPS extracted from the OM of intact cells. For the OMV-derived LPS fractions, an equal volume of the culture was centrifuged to remove intact cells and LPS was extracted from the resulting supernatant. The material from an equivalent volume of the supernatant was loaded in each lane. The Pro-Q 300 stained LPS loading controls confirm the total amount of LPS loaded in each lane. **E. Quantification of background LPS levels in the OM of intact cells and released OMVs as a function of time.** The decrease in background LPS content during exponential phase growth (time = 0 - 2.5 hours post Kdo-N_3_ removal) was consistent with dilution of this LPS subpopulation in the OM of growing and dividing cells. The onset of stationary phase growth is indicated by the red vertical dashed line. During stationary phase (time = 2.5 - 3.5 hours post Kdo-N_3_ removal), the background LPS content in the OM of cells continues to decrease (albeit more slowly) even though the cells are not growing or dividing (filled circles), while the amount of background LPS appearing in the OMV-derived fraction increases drastically (filled squares). These data support a model in which OMV biogenesis acts as a preferential clearance mechanism for background LPS in non-growing, stationary phase cells. **F. Concentration of OMVs in culture during exponential and stationary phases of growth by *E. coli* BW25113 strain in chemically defined medium.** The OMV concentration in a representative bacterial culture was measured at defined time points using nanoparticle tracking. Each data point represents the mean OMV concentration (± SD) measured for technical replicates (N = 3) on a representative sample. The onset of stationary phase growth is indicated by the red vertical dashed line at ∼270 min (equivalent to 2.5 hours post Kdo-N_3_ removal in panel E). Non-linear regression of the OMV concentration increase during stationary phase was well described by an exponential function (*R*^2^ = 0.97).

dSTORM imaging further supported active OMV biogenesis, frequently identifying LPS-rich puncta consistent with nascent or recently released vesicles on the surface of stationary-phase cells ***(Fig. 6B and 6C)***. Such structures were notably more prevalent under growth-arrested conditions aligning with enhanced OMV production in stationary phase. Two- colour dSTORM experiments, using differentially labelled pre-existing background LPS and newly inserted LPS, showed preferential enrichment of background LPS within OMVs ***(Fig. 6C)***. This preferential packaging of older LPS into OMVs strongly supports their role in adaptive OM remodelling during normal (unstressed) growth by a wild-type strain. Similar preferential LPS incorporation into OMVs has been previously observed in *E. coli* ^47^ and in *Salmonella spp.* responding to antimicrobial peptide stress ^46^, suggestive of a conserved OMV- based strategy among Gram-negative bacteria in which OMV-blebbing is used for remodelling of the LPS OM content. To quantitatively assess OMV-mediated LPS turnover, we measured average changes in the distribution of fluorescently labelled, background LPS for OMs isolated from intact cells and an OMV-derived fraction isolated from culture supernatants ***(Fig. 6D, Extended Data Fig. S6C)***. Strikingly, as cultures entered stationary phase and cell growth ceased, OM-associated background LPS fluorescence continued to decrease and was mirrored by a corresponding increase in OMV-associated fluorescence **(*Fig. 6E*)**. This reciprocal relationship provides compelling biochemical evidence that stationary phase cells preferentially shed old LPS via OMVs, in the absence of cell growth and division.

Native OMV production was also assessed by growing the *E. coli* BW25113 strain in the same M9 CDM used for LPS metabolic labelling, removing aliquots of the culture as a function of time, and measuring the vesicle concentration in these aliquots by nanoparticle tracking analysis (NTA) ^48–50^. The OMV concentration was observed to increase by ∼14-fold during mid-exponential growth **(*Fig. 6F*)** when changes in OMV concentration are associated with the growth of rapidly multiplying bacteria. This increase in OMV concentration also coincides with the initial appearance of AF488-labelled LPS in the OMV-derived background LPS fraction analysed by Tricine SDS-PAGE **(*Fig. 6D and 6E*)**. OMV release during normal exponential phase growth has been observed routinely in wild-type Gram-negative bacterial strains ^51^. However, during stationary phase growth the OMV concentration continued to rise, increasing 1.7-fold between 390 min and 750 min post-inoculation of the culture and another 2-fold between 750 min and 1500 min **(*Fig. 6F*)**, even though the bacterial cell density in culture did not change (OD_600_, ***Extended Data Table S23***). Critically, the number of viable cells also remained constant during the stationary phase culture (*i.e*. 390 min to 1500 min post-inoculation) based on a CFU determination done at each time point ***(Extended Data Table S23)***. These results confirm that OMV production begins during mid-to-late exponential growth but also reveal that OMV production occurs continuously during stationary phase even when the cells are not multiplying. Furthermore, these results are consistent with our observation that bacterial cells continue to incorporate LPS metabolically labelled with Kdo-N_3_ in the OM during stationary phase even though the viable cells present are not elongating ***(Extended Data Fig. S7)***.

We propose that OMV formation during stationary phase occurs preferentially in regions of the OM containing older background LPS based on our single-cell **(*Fig. 6C*)** and population level ***(Fig. 6D)*** observations. This OMV-mediated LPS turnover would allow stationary phase *E. coli* to continually remodel and rejuvenate their OM composition, independently of cell growth. The ability of bacteria to preferentially eliminate older LPS through OMV shedding underscores a sophisticated mechanism of OM homeostasis, likely crucial for adaptation and survival under diverse environmental stresses that affect OM integrity, *e.g*. changes in temperature, local divalent cation concentration or exposure to cationic antimicrobial peptides ^11,52,53^.

### Outer membrane buckling model predicts critical OMV size

The mean diameter of the OMVs produced during the different phases of bacterial growth was also determined by NTA (***Extended Data Fig. S8A***) and the size did not change appreciably (= 80-95 nm, ***Extended Data Fig. S8B***). This lack of dependence on growth phase for OMV size has been previously observed with *Neisseria meningitidis* ^50^. The OMV size distribution obtained for stationary phase *E. coli* BW25113 cultures (***Extended Data Fig. S8B***, time = 1500 min) was also consistent with previous work on the same strain ^54^ . We have interpreted this observation as evidence of OMV biogenesis involving the formation of a *critically sized* particle.

Asymmetric growth of the OM relative to the peptidoglycan (PG) cell wall has previously been postulated to favour local outward bulging of the OM, perhaps biased towards regions where a paucity of bonds exists between the PG and Braun’s lipoprotein ^51^ . We developed an OM buckling model (***Fig. 7A***) to test whether accumulation of compressive stress in these membrane regions due to asymmetric growth of the OM relative to the cell wall was sufficient to cause buckling given the intrinsic curvature and physical stiffness of the OM (see Methods section for mathematical derivation). This accumulation of compressive stress will become more widespread in the OM as a cell enters the stationary phase of growth where we have observed that cell elongation slows or stops but LPS incorporation continues ***(Figs. 2, 3 and 6)***. Continued LPS insertion elevates in-plane compressive stress within LPS-dense OM patches pinned at their rims by OMP-rich domains and PG anchoring by Braun’s lipoprotein **(*Fig. 7A*)**, aligning with the load-bearing role of the OM and mechanics of OM-PG tethers ^5,55^. Increased vesiculation is known to occur with weakened OM-PG tethers ^56,57^.

**Figure 7.**
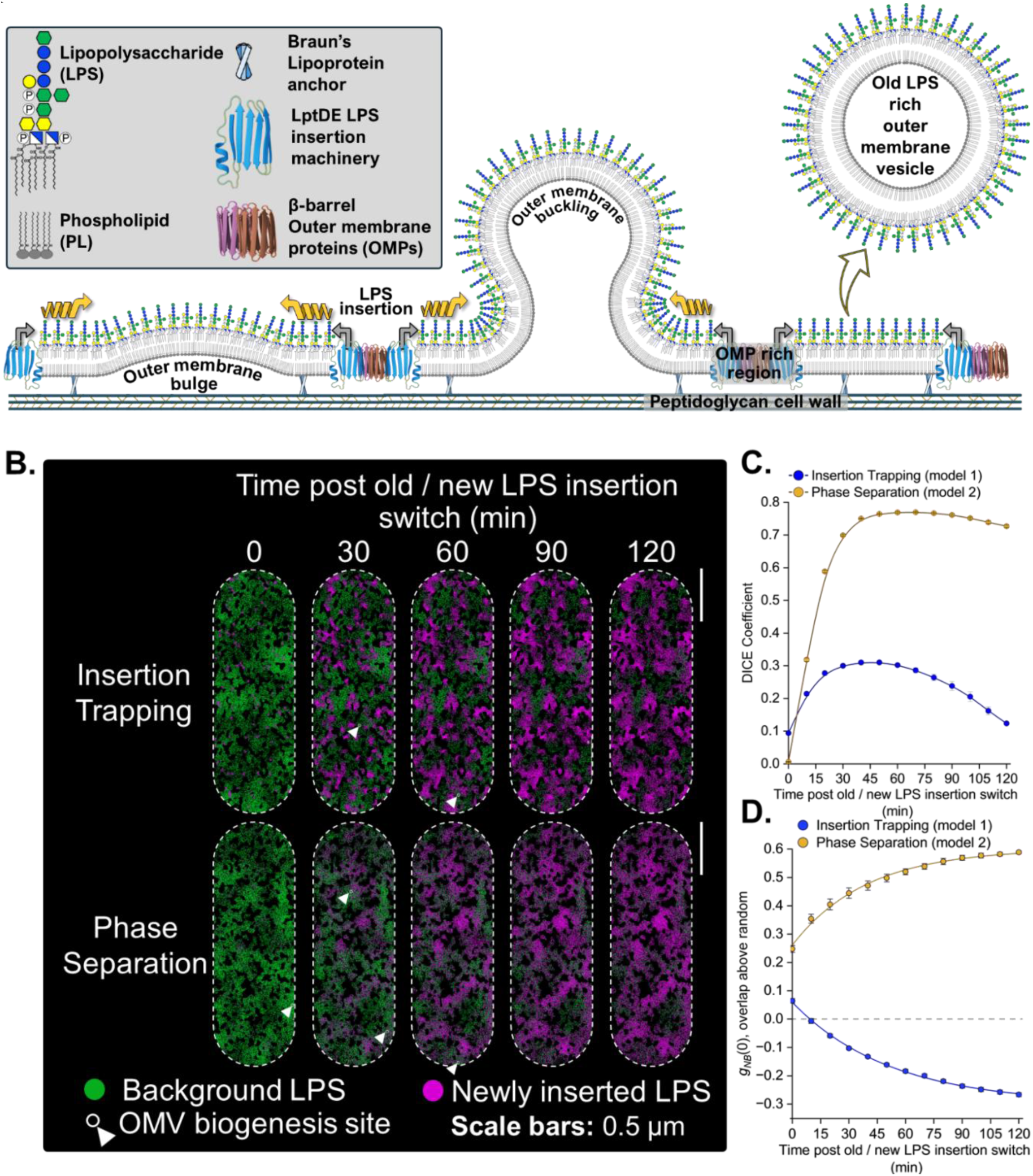
Particle-based simulations indicate an insertion-trapping model rather than a phase separation model predicts the spatiotemporal organisation observed in the *E. coli* outer membrane by two-colour 2D dSTORM SRM imaging. Insertion-trapping and phase separation models were constrained to match the experimentally derived observations and only the organising rule differs (*i.e*. restricted diffusion with hotspots for LPS insertion and short-range Ising/Cahn–Hilliard-style coupling constant for LPS particles, respectively). The total LPS is approximately constant in the accessible membrane area, background LPS half-life ≈ 90 min with new LPS fraction rising to ∼65% by 120 min, and vesiculation events occur randomly in LPS-dense regions with a frequency / size that is consistent with experimental measurements. **A. Model for OMV formation in the OM of a stationary phase *E. coli* cell.** Continuous LPS insertion induces compressive stress in LPS-rich regions which drives OM bulging and blebbing in these regions. **1) OM deformation.** Continuous insertion of new LPS (*grey arrows*) into an LPS-rich region increases local area density while PG expansion lags due to the slow rate of cell elongation in stationary phase. This asymmetric expansion of OM relative to the PG induces in-plane compressive stress in OM (*orange arrows*) resulting in a bulge in the old LPS-dense regions. LPS patch rims are pinned by OMP-rich and PG-anchored zones (for example via Braun’s lipoprotein). **2) Blebbing and neck formation.** The bulge grows into a bleb forming a neck between the surrounding OM and the protruding membrane. **3) OMV formation and compressive stress-release.** The bleb pinches off to produce an OMV, thereby removing older background LPS locally and releasing compressive stress in the OM. This stress-release results in the flattening of the OM around the vesiculation site. **LPS:** Lipopolysaccharide. **OM:** Outer membrane. **OMP:** Outer membrane protein. **PG:** Peptidoglycan. **OMV:** Outer membrane vesicle. **Lpp:** Braun’s lipoprotein. **PL:** phospholipid. **B. Particle-based simulations of the OM under two alternative OM organising models. Insertion-trapping model (top):** New LPS is delivered in bursts at discrete hotspots (LptDE-like sites) and lateral diffusion of both newly inserted and background LPS is strongly confined, resulting in new patches remaining localised while old LPS is depleted locally by OMV formation. **Phase separation model (bottom):** A short-range ‘like–like’ coupling constant favours LPS-LPS contacts irrespective of old or new LPS type and drives domain growth under otherwise identical LPS turnover rates. This short-range favourable interaction tests whether old-new LPS co-clustering could reproduce the experimental observations. **Magenta:** newly inserted LPS. **Green:** background (old) LPS. **Black / vacant spaces**: LPS-inaccessible OMP-occupied voids (∼25% area). **White arrowhead / circle:** Location of OMV biogenesis event arising in LPS-dense region according to the OM buckling model. **Scale bars:** 0.5 µm. **C. Time courses for newly inserted and background LPS from both models after DICE analysis of DBSCAN co-clustering.** Cluster masks were built from core points only (noise and non-core border points were excluded before rasterisation, ε = 30 nm; minPts = 7) and rasterised on the simulation grid (3.33 nm pixel⁻¹) within the allowed membrane. Points show the mean ± SEM (n = 5 independent simulation runs per model) per time point. The phase-separation model (orange) rises monotonically toward a high plateau, indicating persistent co-clustering as domains coarsen. The insertion-trapping model (blue) increases to a maximum around 60 min and then declines toward the starting level by 120 min, consistent with sustained segregation as newly inserted LPS replaces background LPS locally. DICE is unitless (0–1). **D. Zero-lag coverage-normalised excess overlap of cluster masks (*g_NB_(0)*) quantifying colocalisation of newly inserted and background LPS relative to random placement (horizontal dashed line at 0)**. Coverage-normalised excess overlap of the same DBSCAN masks, computed as *gNB*(0) = *pAB − pA pB* (dimensionless), where *pA* and *pB* are the fractional areas of old and new cluster masks and *pAB* their joint occupancy. Dashed line marks 0 (random overlap). Points show mean ± SEM (n = 5 independent simulation runs per model). The phase-separation control (orange) yields positive values that increase and then saturate, reflecting growing co-location. The insertion-trapping model (blue) remains at or below zero and becomes more negative over time, indicating anti-association.

The OM buckling model **(*Fig. 7A*)** was used to predict the *critical size* of the OMV released in background LPS-rich regions (see Methods section for model calculations), which can be compared to our experimentally determined diameters (***Extended Data Fig. S8B***). When a bulging LPS-rich OM patch exceeds a characteristic radius (*R_b_*), a bleb forms and is released as an OMV with a critical diameter (*d*). We estimated *d* = 73-122 nm for OMVs released by this membrane buckling mechanism which agrees with our experimental results (***Extended Data Fig. S8B***, grey shaded region). This compressive stress-release by OM blebbing is consistent with bilayer-coupled, leaflet-asymmetry routes to OMV formation ^58,59^ and with broad OMV biogenesis frameworks ^43,44^.

### Particle-based simulations of LPS spatiotemporal dynamics distinguish insertion– trapping mechanism from phase separation in the OM

To test whether the spatiotemporal LPS behaviour observed by pulse–chase dSTORM (***Fig. 2, Extended Data Fig. S2)*** requires equilibrium phase separation, we compared two particle-based simulations of the *E. coli* outer membrane that share identical, experimentally-derived LPS turnover rates ***(Figs. 2, 3 and 6)***, but differ in their organising rules ***(Fig. 7)***. The simulated field covers half of the OM surface of the cell (2.6 µm × 1.1 µm) and contains a static “allowed” mask (75% of the area) representing LPS-accessible membrane ^60–62^. The remaining area (25%) is considered as OMP-rich, LPS-inaccessible voids (black regions in schematics), thereby creating the heterogenous OMP and LPS clustered OM organisation observed in previous experimental studies ^10,11,21^. The total LPS in the accessible area is held approximately constant (target N ≈ 8×10^4^ LPS simulation particles, ∼9-fold coarse-graining of the expected biological copy number ^61–63^). LPS turnover and insertion rates were informed by experimental data presented in **Figures 2, 3 and 6**. Background (‘old’) LPS decays with a half-life of ∼90 min, newly inserted (‘new’) LPS rises to ∼65% by 120 min; and OMV events are placed stochastically at LPS-density maxima with realistic footprints (80-90 nm radii) and frequencies matched to experimental data ***(Fig. 6, Fig. 7A and 7B)***. Thus, these constraints reproduce the measured turnover (reciprocal changes in old and new LPS coverage, and weak old-new co-clustering around ∼60 min, ***Fig. 2F***) and ensure that any divergence between models reflects organisation rather than bulk insertion rates. To facilitate robust quantitative comparison with the imaging analyses, cluster masks were defined by DBSCAN using identical cluster-defining parameters (ε = 30 nm; minPts = 7; core points only). We tracked the normalised old–new overlap (DICE, Sørensen–Dice coefficient) and the zero-lag coverage-normalised overlap of cluster masks, *g_NB_*(0), where 0 denotes the level expected for random colocalisation of newly inserted and background LPS particles, while negative values indicate segregation of the two species and positive values indicate colocalisation ***(Fig. 7C and 7D)*.**

The insertion-trapping model was meant to reproduce our experimental observations and incorporated the restricted lateral diffusion of LPS in the OM ***(Fig. 7B, Extended Data Fig. S9A).*** New LPS is inserted in short bursts at a changing subset of pre-sampled hotspots located across the entire OM surface (consistent with the mapped LptD-linked LPS delivery sites characterised in ***Fig. 1D***) ^10,24^, while both background and newly inserted LPS undergo strongly confined lateral diffusion in-line with previous observations ^10,11^. Background LPS loss is biased away from newly inserted LPS-rich locations at the edges of OMP-rich, LPS- inaccessible voids, and OMV budding sites are drawn from the top LPS density quantile, consistent with vesiculation from LPS-dense patches. Notably this model reproduces the following three key experimental observations.

Despite substantial turnover, the model preserves spatial segregation ***(Fig. 7B-7D)***, the normalised old–new overlap or co-clustering stays low with a weak maximum at ∼60 min ***(Figs. 2 and 7C, Extended Data Fig. S9)***, and *g_NB_*(0) remains ≤ 0 from 10-120 min ***(Fig. 7D)***. Cluster sizes of new LPS patches broaden modestly as coverage grows as was observed experimentally ***(Fig. 4)***, but there is no late-stage, OM-wide coarsening.

The phase separation model was used to emulate old-new LPS co-clustering in the OM due to phase separation while keeping all bulk rates and the restricted lateral diffusion of LPS particles identical to the insertion-trapping model **(*Fig. 7B, Extended Data Fig. S9A*).** We introduced a short-range like–like coupling constant (Ising/Cahn–Hilliard-style drift up the density gradient of each LPS type) ^64^ with strength tuned so that global densities and OMV statistics match the output from the insertion-trapping model. Strikingly, and in sharp contrast to the insertion-trapping model and our experimental data, this model predicts the opposite time dependencies. The simulation shows a monotonic increase in the overlap of background and newly inserted LPS ***(Fig. 7C)***, a rise in *g_NB_*(0) above zero ***(Fig. 7D)*** and a gradual right-shift in domain-size distribution or coarsening, which are hallmarks of phase ordering. Critically, these behaviours were not observed in our experimental data ***(Figs. 2 and 4)*** ^64^.

We further connected spatiotemporal LPS dynamics to vesicle biogenesis by incorporating membrane buckling-induced OMV formation in our simulations. In the simulation, this is captured phenomenologically by Poisson-distributed OMV formation events at LPS-density maxima that result in the removal of local, background LPS without leaving holes, reproducing the observed depletion of background LPS at vesiculation sites while maintaining overall membrane coverage ***(Fig. 6A and 6C, Fig. 7A and 7B, Extended Data Fig. S9)***. OM buckling-driven vesiculation links new LPS insertion to the release of background LPS-rich regions in OMVs. With experimental LPS turnover rates, the insertion–trapping model yielded persistent old-new LPS segregation, low and transient old-new LPS co-clustering, and an absence of coarsening, all of which aligns strongly with our *in vivo* experimental data ***(Figs. 2, 4 and 6)*.** This sharply contrasts with the phase separation model, incorporating identical experimentally-derived turnover rates, which predicts increased co-clustering of old and new LPS and domain growth (coarsening). The output of these particle-based simulations supports a kinetically-trapped OM organisation which is maintained in a nonequilibrium state by site-specific insertion of OM components and their restricted lateral diffusion. OMV biogenesis driven by continuous LPS insertion contributes to the preferential loss of older background LPS, enabling stationary phase *E. coli* cells to modify and adapt the LPS content of their OM in a growth-independent manner.

## Discussion

Our experiments reveal a dynamic, yet tightly constrained organisation of the LPS content in the *E. coli* OM. We found that LPS turnover in the OM is not static, even in stationary phase cells, where new LPS continues to be inserted and old LPS is preferentially removed (***Figs. 2-4 and 6)***. However, this turnover occurs without any large-scale mixing of LPS within the membrane ***(Figs. 2 and 4)***. Instead, the OM maintains a “frozen” mosaic organisation where domains are preserved over time. These findings strongly suggest a refined model for OM organisation and homeostasis in Gram-negative bacteria. In this model, the spatial arrangement of LPS is governed not by thermodynamically-driven phase separation into larger domains but by localised insertion events coupled with extremely limited lateral diffusion that effectively trap LPS molecules near their point of insertion (insertion-trapping mechanism). Meanwhile, the removal of LPS from the OM in stationary phase cells, where passive dilution is hindered due to the absence of growth-dependent OM expansion, is achieved through vesiculation.

This work will profoundly influence the way we consider the OM organisation, while it has also confirmed the extremely limited lateral mobility of LPS in the OM ^10,11^. Using two-colour SRM, we directly visualised that newly inserted LPS molecules remain spatially segregated from older LPS domains even after hours of incubation ***(Fig. 2)***. The persistence of distinct, newly inserted and background LPS patches in our pulse–chase experiments indicate minimal mixing between the two populations (***Figs. 2-4)***. This provides an intuitive, visual confirmation of what earlier single-particle tracking data had implied: LPS lateral diffusion in the OM is so slow (∼10^-2^ μm²/s or lower) ^10,11^ and restricted that an LPS molecule essentially remains near its point of insertion. Notably, our data show that even across multiple generations, LPS does not redistribute homogeneously. Instead, each cohort of LPS inserted at a given time retains its identity as a cluster, “frozen” in place. Consequently, LPS does not undergo binary partitioning in the manner of OMPs ^20^ which are inserted at mid-cell and become apportioned to new cell poles during cell elongation and division ^20^. This means that in the absence of growth and division old LPS would accumulate indefinitely in the OM, limiting the ability of the bacterium to remodel its lipid composition in response to environmental change. *E. coli* resolves this by shedding old, background LPS via OMVs, enabling OM renewal and adaptation even in non-growing cells. This process appears to be a fundamental aspect of OM maintenance in non-growing *E. coli* cells. By generating OMVs, the bacterium can preferentially eject portions of its OM which are rich in background or ‘old’ LPS ^46,47,58,65^.

Our time-lapse imaging ***(Figs. 6A-C)*** captured nascent vesicles with old LPS budding off cells, and our bulk fractionation experiments showed a clear transfer of background LPS from the OM of cells to vesicles, with both events occurring as stationary phase ensued ***(Fig. 6D)***. Notably, we observed a substantial increase in OMV production precisely when cell growth ceased **(*Fig. 6F*)**, which suggests that vesicle shedding is increased in response to entry into stationary phase, consistent with prior reports of elevated OMV formation under stress or nutrient limitation ^66–68^. This vesicle-mediated LPS turnover allows the cell to “rejuvenate” its OM, with background LPS molecules continuously removed and replaced by newly inserted molecules, even when the cell is not dividing. Such a mechanism is likely important for preserving OM integrity and functionality during a prolonged period in stationary phase, and for facilitating adaptation in situations where growth is arrested due to nutrient-limited conditions when the environment may still be challenging (for example, in host tissues, acidic lysosomes post-phagocytosis or in biofilms). By discarding a portion of its endotoxin via OMVs, a bacterial cell might also modulate its interactions with the host immune system or neighbouring microbes, though the specific consequences of shedding old LPS remain to be explored.

To test whether the observed spatiotemporal dynamics arise from phase separation or kinetic-trapping, we built simulations of OM expansion constrained by experimental rates of LPS insertion and turnover ***(Fig. 7, Extended Data Fig. S9)***. The insertion-trapping model ***(Fig. 7B, Extended Data Fig. S9)*** yielded stable, discrete patches of newly inserted and background LPS ***(Figs. 1, 2 and 7, Extended Data Fig. S9)***, loss of background LPS without coarsening, and sustained spatial segregation, quantified by a weak maximum in the old-new LPS overlap, and *g_NB_(0) ≤ 0* across 0 – 120 min ***(Figs. 2, 4 and 7C-7D).*** Therefore, this model closely reproduced the hallmark features seen in the dSTORM images ***(Figs. 2-4).*** In sharp contrast, a phase separation model with identical bulk LPS insertion and turnover rates ***(Fig. 7B, Extended Data Fig. S9A)*** showed contrasting LPS spatiotemporal dynamics, yielding instead characteristics expected for phase ordering and liquid–liquid demixing in model membranes ^28,69^. This included a monotonic rise in colocalization (*g_NB_(0)* > 0) ^64,70^, and growth of the LPS domain size **(*Fig. 7C and 7D*)**. Incorporation of membrane buckling-driven OM vesiculation into the simulations ***(Fig. 7A)*** provided a biophysical mechanism for how an increase in local compressive membrane stress due to continued LPS insertion could drive OMV release. Poisson distributed vesiculation events nucleated at LPS particle density maxima, where LPS-rich patch centres are weakly anchored to PG, removed old LPS content from the OM, and relieved compressive stress without leaving persistent voids. These simulation outputs mirrored our dSTORM imaging and bulk fractionation data ***(Figs. 2, 3 and 6)***, and aligned with established OM mechanics and OMV biogenesis paradigms ^43,44,56–59^.

Together, these particle-based simulations support a kinetically-trapped, non-equilibrium OM spatiotemporal organisation, the properties of which are dictated by site-specific insertion of LPS, restricted lateral diffusion of OM components ^10,11,20^ and OMV-mediated preferential removal of older LPS. The concept of a kinetically-trapped OM organisation, recapitulated by the simulations, also explains the stable coexistence of LPS-rich and OMP-rich domains without the need for equilibrium (thermodynamically-driven) phase separation ^10,11^. In classical phase separation, one would expect coarsening over time with small domains merging into fewer, larger domains to minimise interfacial energy. We see no evidence of such coarsening ***(Figs. 2 and 4)***. Instead, the patterns we observed (many small LPS patches that persist and do not fuse) align with a scenario where membrane components are inserted in certain areas and then constrained from moving far ***(Figs. 2 and 4)***. Our clustering analysis showed that new LPS clusters proliferate in number but not in size, and that old clusters dissipate without merging into others ***(Fig. 4)***. This is inconsistent with Ostwald ripening and other phase separation phenomena, and is better described by a diffusion-limited, far-from-equilibrium process. In essence, the crowded, rugged energy landscape of the OM (shaped by strong LPS–LPS interactions, peptidoglycan tethers, and limited lateral mobility) means that whatever arrangement of molecules is established through insertion is essentially trapped in place, giving rise to a patchwork of persistent nanodomains. Our findings thus bridge mechanistic insights at the molecular level (LPS transport and diffusion) with the mesoscopic organisation of the membrane.

In summary, this study highlights a previously unappreciated dynamism in the *E. coli* OM. We show that, far from being a static barrier, the OM actively undergoes renewal even in stationary phase. New LPS continues to be incorporated, and background LPS is continuously shed via OMVs. This mode of LPS turnover ensures that the OM can be remodelled and adapt to changing conditions without requiring cell growth or division. The advanced imaging and labelling strategies we employed were crucial for uncovering these phenomena, enabling us to directly observe LPS being inserted and removed *in vivo*. In future studies, the high spatial precision methodologies employed here for studying LPS turnover, might also be applied in the exploration of turnover (or reorganisation) of other OM components (such as lipoproteins or certain OMPs) in non-dividing cells, and to determine whether similar vesicle-mediated processes occur in other Gram-negative species.

More broadly, our work emphasises the importance of OM vesiculation as not only a means of secreting molecules but also a mechanism for turnover and renewal of its fundamental glycolipid component. Understanding how bacteria manage the composition and organisation of their envelopes in different growth states could inform new strategies to target their vulnerabilities, especially in contexts like chronic infections where many invading pathogenic Gram-negative cells are in a growth-arrested state.

## Supporting information

Codes and Read.me files - referenced clearly where relevant in methods

Extended Data Figures and Tables - referenced clearly where relevant in manuscript

Extended Methods - referenced where relevant in manuscript

## Acknowledgements

We thank the University of York Bioscience Technology Facility (BTF) for access to microscopy facilities, and G. Calder (BTF) and N. Sergent (Zeiss) for expert assistance with dSTORM and 3D-SIM^2^ data collection and analysis. We thank E. Lemke for the pEVOL-pylRS plasmid, C. Sharrock for assistance with site-directed mutagenesis in LptD, F. Hands for technical assistance with OMV purification and characterisation, R. Farley for assistance with NTA data acquisition and analysis, and Mesenbio Ltd. for access to their ZetaView® Quatt NTA instrument. This work was supported by The University of York, the BBSRC (19ALERT Mid-Range Equipment Initiative Award to the Department of Biology to purchase ZEISS Elyra 7 system, BB/T017589/1), the BBSRC White Rose Doctoral Training Partnership (PhD studentship award to J.N., 2272649), and a Horizon Europe Guarantee Award to M.A.F. (selected by the ERC and funded by UKRI, EP/X023680/1).

## Author contributions

J.N., M.A.F. and C.G.B. designed the experiments. J.N. characterised the native and fluorescently labelled LPS glycoforms and OMPs. N.E.H. and M.A.F. performed chemical synthesis. J.N. collected and analysed ensemble-averaged LPS turnover data for intact cell- and OMV-derived fractions. C.G.B. collected and analysed NTA data with assistance from J.N. J.N. collected and analysed FRAP, dSTORM and live-cell 3D-SIM^2^ data with assistance from C.G.B. J.N. with assistance from D.O.P. designed Python and MATLAB scripts for dSTORM image analysis and interrogation. J.N., D.O.P., C.G.B., and M.A.F. devised model for OM organisation. D.O.P., C.G.B. and J.N. devised the OM buckling model. J.N. prepared all figures with input from C.G.B., D.O.P. and M.A.F. J.N., M.A.F. and C.G.B. wrote the manuscript. All co-authors had the opportunity to comment on the final submitted version of the manuscript.

