## Supplementary material for "Persistent LPS insertion, spatial segregation and vesicle biogenesis drive growth-independent adaptation of the *Escherichia coli* outer membrane": Codes and Read.me files - referenced clearly where relevant in methods: new_lps_or_old_LPS_cluster_summary_graphs_code.txt

%% Summary Graphs for LPS Cluster Properties
% Version: 1.0 (2025-10-01)
% Authors: Joe Nabarro, Christoph G. Baumann, Martin A. Fascione
% Contact:
% Institution: University of York
%
% Description:
%   Generate aggregate plots (means ± s.e.m.) for cluster counts, areas, and densities per region and condition.
%
% Inputs:
%   DBSCAN summary spreadsheets per replicate/time
%
% Outputs:
%   PNG/PDF plots for Extended Data
%
% Dependencies:
%   MATLAB R2024b or later; Statistics and Machine Learning Toolbox (for pca/DBSCAN if used).
%
% Usage:
%   See README.md in the root of this package for end-to-end instructions and example invocations.
%
% Notes:
%   This script implements the analysis described in the main text and Methods.
%   Parameter values and bin sizes were chosen to match the reported analyses.
%

%% new_lps_or_old_LPS_cluster_summary_graph.m
% This script generates a publication-quality scatter plot to visualise how LPS cluster number,
% surface area, and localisation density change over time during pulse–chase experiments.
% It fits a single-phase exponential model to cluster number data and encodes surface area and
% density using marker size and colour, respectively.

%% 1. Load clustering summary data
% Replace the filename below with your own Excel (.xlsx or .csv) file.
% Expected columns: time (min), cluster_number (per µm²), cluster_area (nm²), cluster_density (localisations/nm²)

filename = 'your_clustering_summary_file.xlsx';  % Insert your summary stats file here
data = readtable(filename);
data = data(:, 1:4);  % Ensure correct format if file includes extra columns
data.Properties.VariableNames = {'time', 'cluster_number', 'cluster_area', 'cluster_density'};

%% 2. Preprocess variables
time = data.time;
cluster_number = data.cluster_number;

% Convert area from nm² to µm² for plotting
cluster_area = data.cluster_area / 1e6;

% Convert density from localisations/nm² to localisations/µm²
cluster_density = data.cluster_density * 1e6;

% Handle potential outliers or negative values in area
cluster_area(cluster_area < 0) = NaN;

% Rescale cluster areas for visual encoding as marker size (range: 50–1000 pts²)
sizes = rescale(cluster_area, 50, 1000);  % Adjust range here if desired

%% 3. Generate scatter plot
fig = figure('Color', 'w');
scatter(time, cluster_number, sizes, cluster_density, ...
    'filled', 'MarkerEdgeColor', 'k', 'MarkerFaceAlpha', 0.85);
colormap(jet);  % Set colormap for density
hold on;

%% 4. Fit exponential model to cluster number vs time
% Fit model: y = a * exp(b * x)
ft = fittype('a * exp(b * x)');
opts = fitoptions('StartPoint', [max(cluster_number), 0.01]);  % Adjust start point if needed
[expFit, gof] = fit(time, cluster_number, ft, opts);

% Evaluate and plot fitted curve
xfit = linspace(min(time), max(time), 300);
yfit = expFit.a * exp(expFit.b * xfit);
plot(xfit, yfit, '--k', 'LineWidth', 1.5);

%% 5. Label axes and add legend
xlabel('Time (min)', 'FontSize', 12);
ylabel('Clusters per µm²', 'FontSize', 12);

% Add colourbar for cluster density
cb = colorbar;
cb.Label.String = 'Cluster Density (localisations / µm²)';
cb.Label.FontSize = 12;

% Add dynamic legend entry with equation
legend({'Cluster trend', ...
    sprintf('y = %.2f × exp(%.4f x), R² = %.2f', ...
    expFit.a, expFit.b, gof.rsquare)}, ...
    'Location', 'northwest');

%% 6. Export outputs
% Replace filenames below with desired output names
exportgraphics(fig, 'LPS_cluster_trend_plot.png', 'Resolution', 700);
writetable(data, 'LPS_cluster_trend_summary_output.xlsx');
