## Supplementary material for "Persistent LPS insertion, spatial segregation and vesicle biogenesis drive growth-independent adaptation of the *Escherichia coli* outer membrane": Codes and Read.me files - referenced clearly where relevant in methods: Python_code_New_Old_LPS_DBSCAN.txt

#!/usr/bin/env python3
# -*- coding: utf-8 -*-
"""
Background vs New LPS Coverage & Normalised Overlap (DBSCAN-based)
Version: 1.0 (2025-10-01)
Authors: Joe Nabarro, Christoph G. Baumann, Martin A. Fascione
Contact:
Institution: University of York

Description:
  Compute outer-membrane (OM) coverage by background ("old") and newly inserted ("new") LPS clusters
  and the normalised spatial overlap as a function of time/condition. The script accepts tables of DBSCAN
  clusters for each population and exports per-cell and aggregated results along with figures, matching
  the manuscript (Main Text Fig. 2E–F; Methods: DBSCAN and overlap analysis).

Inputs:
  - CSV files for "old" and "new" LPS localisations or DBSCAN cluster tables. Expected columns:
    * For localisations: ['Position X [nm]', 'Position Y [nm]'] (+ optional photon/precision fields)
    * For clusters: ['CellID','CentroidX_nm','CentroidY_nm','Area_nm2','Time_min' or 'Condition']
  - Optional parameters via CLI to set pixel size, convex-hull buffer, region-of-interest (ROI) dilation, etc.

Outputs:
  - Per-cell CSV: coverage_old, coverage_new, overlap_area, normalised_overlap
  - Summary CSV: time/condition means ± s.e.m.
  - Figures: PNG/PDF plots (coverage vs time; normalised overlap vs time with 2nd-order polynomial fit)

Dependencies:
  Python 3.9+, numpy, pandas, scipy, shapely (>=2.0), matplotlib

Usage:
  python DBSCAN_background_vs_newLPS_coverage_overlap.py --old path/to/old.csv --new path/to/new.csv --out results/

Notes:
  - Normalised overlap = overlap_area / (area_old + area_new)
  - ROI is estimated from the combined convex hull (optionally buffered) of all localisations in a cell.
  - For cluster tables, areas are unioned as shapely polygons to compute coverage and overlaps.
"""


# Quantitative analysis of spatial overlap between background
# (“old”) and newly inserted (“new”) LPS clusters in individual
# E. coli cells imaged by dual‑colour 2D dSTORM.

# Author: Joe Nabarro
# Date: Spring 2025
# Institution: University of York
# Contact:

# Print Python environment info (useful for reproducibility or debugging)
import sys
# === Standard and scientific libraries ===
import os
import numpy as np
import pandas as pd
import matplotlib.pyplot as plt

# Geometry handling for clustering and shape analysis
from shapely.geometry import Point, MultiPoint, Polygon
from shapely.ops import unary_union
from shapely.affinity import rotate

# Convex hull (for defining OM boundary) outlines cell by finding the outer boundary of all (combined old and new) defined dSTORM emissions
from scipy.spatial import ConvexHull

# Clustering and dimensionality reduction
from sklearn.cluster import DBSCAN
from sklearn.decomposition import PCA

#  Load old and new LPS CSV files 
old_path = "input background LPS dSTORM table path in .csv format here"
new_path = "input newly inserted LPS dSTORM table path in .csv format here"

# Load into pandas DataFrames
old_lps = pd.read_csv(old_path)
new_lps = pd.read_csv(new_path)

#  Set up output folder to save figure 
base_name = os.path.splitext(os.path.basename(old_path))[0] + "__" + os.path.splitext(os.path.basename(new_path))[0]
out_folder = os.path.join(os.path.dirname(old_path), base_name)
os.makedirs(out_folder, exist_ok=True)  # Create folder if it doesn't exist

#  Combine LPS coordinates to define the cell outline 
combined = pd.concat([
    old_lps[['Position X [nm]', 'Position Y [nm]']],
    new_lps[['Position X [nm]', 'Position Y [nm]']]
])
all_coords = combined.to_numpy()

#  Compute convex hull + buffer to approximate the OM shape 
hull = ConvexHull(all_coords)
smoothed = all_coords[hull.vertices]
# It expands the convex hull outward by 20 nanometers in all directions.
#Think of it like adding padding or a cushion around the shape. Simplify smoothss polygon removing kinks and jagged edges with 2 nm tolerance
om_shape = MultiPoint(smoothed).convex_hull.buffer(20).simplify(2)  # Expand & smooth

# Align shape along its long axis using PCA 
mean_coords = all_coords.mean(axis=0)
pca = PCA(n_components=2).fit(all_coords - mean_coords)
angle_rad = np.arctan2(pca.components_[0,1], pca.components_[0,0])  # Angle of first principal component
angle_deg = np.degrees(-angle_rad)  # Clockwise rotation
centroid = tuple(om_shape.centroid.coords[0])  # Centroid of OM shape
om_shape = rotate(om_shape, angle_deg, origin=centroid)  # Rotate shape to align long axis horizontally

#  Function: identify cluster polygons using DBSCAN and clip to OM 
def get_cluster_shapes(coords, eps=30, min_samples=7):
    clustering = DBSCAN(eps=eps, min_samples=min_samples).fit(coords)
    shapes = []
    for lbl in set(clustering.labels_):
        if lbl == -1: continue  # skip noise
        pts = coords[clustering.labels_ == lbl]
        hull = MultiPoint(pts).convex_hull
        clipped = hull.intersection(om_shape)  # restrict to OM
        if not clipped.is_empty and clipped.area > 0:
            shapes.append(clipped)
    return shapes

# Detect and rotate cluster shapes for both old and new LPS 
old_shapes = get_cluster_shapes(old_lps[['Position X [nm]', 'Position Y [nm]']].values)
new_shapes = get_cluster_shapes(new_lps[['Position X [nm]', 'Position Y [nm]']].values)
old_shapes = [rotate(s, angle_deg, origin=centroid) for s in old_shapes]
new_shapes = [rotate(s, angle_deg, origin=centroid) for s in new_shapes]

#  Compute union of all old and new cluster shapes, then find intersection (overlap) 
old_union, new_union = unary_union(old_shapes), unary_union(new_shapes)
overlap_geom = old_union.intersection(new_union)  # Shared area = overlap

#  Rotate all individual emission points 
old_points = [rotate(Point(x, y), angle_deg, origin=centroid)
              for x, y in old_lps[['Position X [nm]', 'Position Y [nm]']].values]
new_points = [rotate(Point(x, y), angle_deg, origin=centroid)
              for x, y in new_lps[['Position X [nm]', 'Position Y [nm]']].values]
old_coords = np.array([(p.x, p.y) for p in old_points])
new_coords = np.array([(p.x, p.y) for p in new_points])

# Re-run DBSCAN on rotated coordinates to classify clustered vs. noise points 
db_old = DBSCAN(eps=30, min_samples=6).fit(old_coords)
old_labels = db_old.labels_
old_cluster, old_noise = old_labels != -1, old_labels == -1  # Boolean masks

db_new = DBSCAN(eps=30, min_samples=6).fit(new_coords)
new_labels = db_new.labels_
new_cluster, new_noise = new_labels != -1, new_labels == -1

# Identify individual points falling inside the overlapping polygon area 
overlap_old = [(p.x, p.y) for p in old_points if overlap_geom.contains(p)]
overlap_new = [(p.x, p.y) for p in new_points if overlap_geom.contains(p)]

# Start plotting 
fig, ax = plt.subplots(figsize=(10,6))

# turn off the grid
ax.grid(False)

#  Plot old/new noise points as semi-transparent circles
ax.scatter(old_coords[old_noise,0], old_coords[old_noise,1], c='green', s=30, alpha=0.3,
           edgecolor='black', linewidth=0.3, label='Old LPS')
ax.scatter(new_coords[new_noise,0], new_coords[new_noise,1], c='darkviolet', s=30, alpha=0.3,
           edgecolor='black', linewidth=0.3, label='New LPS')

# Plot clustered points as more prominent circles -
ax.scatter(old_coords[old_cluster,0], old_coords[old_cluster,1], c='green', s=30, alpha=0.7,
           edgecolor='black', linewidth=0.3)
ax.scatter(new_coords[new_cluster,0], new_coords[new_cluster,1], c='darkviolet', s=30, alpha=0.7,
           edgecolor='black', linewidth=0.3)

#  Plot overlapping points in aqua blue 
if overlap_old:
    ox, oy = zip(*overlap_old)
    ax.scatter(ox, oy, c='blue', s=30, alpha=0.9, edgecolor='black', linewidth=0.4, label='Old Overlap')
if overlap_new:
    nx, ny = zip(*overlap_new)
    ax.scatter(nx, ny, c='blue', s=30, alpha=0.9, edgecolor='black', linewidth=0.4, label='New Overlap')

# Add scale bar 
bar_nm = 500             # Length in nanometers
start_frac = 0.05        # X-start (in axes fraction units)
dx = ax.get_xlim()[1] - ax.get_xlim()[0]
bar_frac = bar_nm / dx   # Convert nanometers → relative axis fraction
bar_y = 0.01             # Y-position (in axes fraction units)

# Draw scale bar line
ax.plot([start_frac, start_frac+bar_frac], [bar_y, bar_y],
        transform=ax.transAxes, color='black', lw=2, solid_capstyle='butt')

# Add scale bar label
ax.text(start_frac+bar_frac/2, bar_y - 0.025, '500 nm',
        transform=ax.transAxes, ha='center', va='top', fontsize=10)

#  Remove visual clutter from axes 
for spine in ax.spines.values():
    spine.set_visible(False)
ax.tick_params(left=False, bottom=False, labelleft=False, labelbottom=False)
ax.set_xlabel('')
ax.set_ylabel('')

#  Set figure aesthetics 
ax.set_aspect('equal')
ax.set_title('Aligned Combined Overlap Map', fontsize=14, pad=15)

# Add clean legend beside the plot
ax.legend(frameon=False, loc='center left', bbox_to_anchor=(1.02,0.5), fontsize=10)

# Final layout & export
plt.tight_layout()
plt.savefig(os.path.join(out_folder, 'combined_overlap_map_clustered_highres.png'),
            dpi=1200, bbox_inches='tight')  # Save high-resolution figure
plt.show()

if __name__ == '__main__':
    pass
