## Supplementary material for "Persistent LPS insertion, spatial segregation and vesicle biogenesis drive growth-independent adaptation of the *Escherichia coli* outer membrane": Codes and Read.me files - referenced clearly where relevant in methods: READ.ME_LptD_new_LPS_crosscorrelation.txt

Purpose: Compute radial cross-correlation g(r) between LptD* and newly inserted LPS from dual-colour dSTORM.
Inputs: Two CSVs (LptD, new LPS) with X_nm, Y_nm 
Run: Open the MATLAB version or adapt to your environment; set input paths at the top of the script.
Outputs: g(r) per cell (CSV) and summary plot(s).
