## Extended Data Figures and Tables - referenced clearly where relevant in manuscript for "Persistent LPS insertion, spatial segregation and vesicle biogenesis drive growth-independent adaptation of the *Escherichia coli* outer membrane"

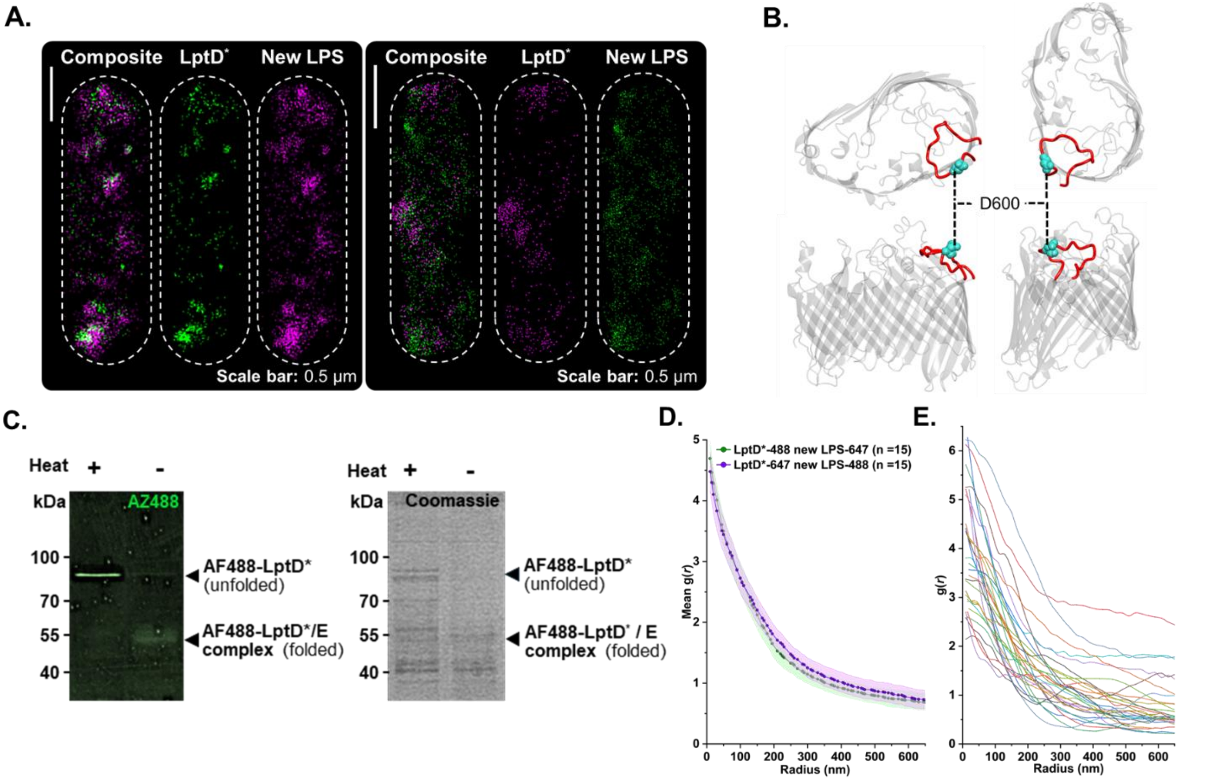


**Extended Data Figure S1. Dual-colour labelling confirms site-specific incorporation and co-localisation of functional LptD* and newly inserted LPS.**

A. **Additional dual-colour dSTORM images of *E. coli* outer membranes labelled for recombinant LptD* and newly inserted LPS 30 min after co-addition of N-propargyl-L-lysine and Kdo-$N_3$.** LptD* was mutated with an amber stop codon replacing D600 in extracellular loop 9 and expressed via genetic code expansion, enabling site-specific conjugation to an azide functionalised fluorophore by CuAAC. Newly inserted LPS was metabolically labelled with Kdo-$N_3$ enabling site-specific conjugation to an alkyne functionalised fluorophore via CuAAC. Discrete, co-localised puncta are observed across the OM, independent of dye combination. **Scale bars:** 0.5 µm.

B. **LptD crystal structure (PDB:** 4HRB**) highlighting loop 9 (red) and D600 mutation site used for ncAA incorporation.**

C. **In-gel AF488 fluorescence (*left*) and Coomassie-stained (*right*) SDS-PAGE analysis of cell lysates containing AF488-labelled LptD*.** Fluorescent LptD* and native LptD*/LptE complex were visualised in unheated samples (–), while heat-denatured samples (+) show dissociated LptD* with reduced mobility, confirming that ncAA labelling does not disrupt LptD/LptE* complex formation.

D. **Mean cross-correlation profiles for LptD*–AF488 / new LPS-AF647 (n = 15) and LptD*-AF647 / new LPS–AF488 (n = 15) reciprocal dye configurations show near identical, strong sub-diffraction co-localisation, indicating spatial correlation was independent of dye-selection.**

E. **Single-cell cross-correlation profiles (n = 30) showing consistent strong sub-diffraction co-localisation between LptD* and newly inserted LPS across the population of screened *E. coli* OMs.**

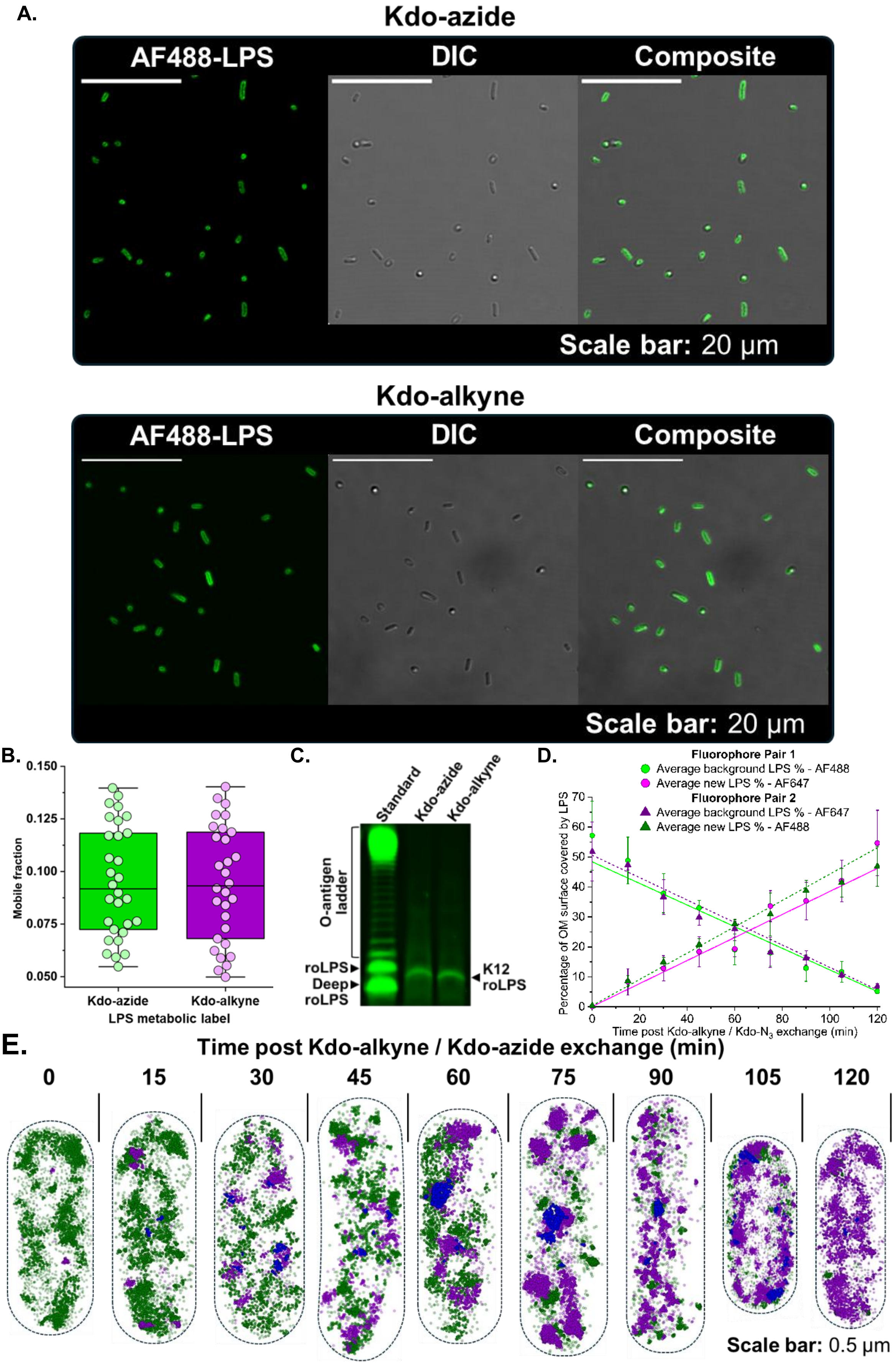

A.
Kdo-azide
AF488-LPS
DIC
Composite
Scale bar: 20 μm
Kdo-alkyne
AF488-LPS
DIC
Composite
Scale bar: 20 μm
B.
Mobile fraction
Kdo-azide
Kdo-alkyne
LPS metabolic label
C.
Standard
Kdo-azide
Kdo-alkyne
O-antigen ladder
roLPS
Deep roLPS
K12 roLPS
D.
Fluorophore Pair 1
Average background LPS % - AF488
Average new LPS % - AF647
Fluorophore Pair 2
Average background LPS % - AF647
Average new LPS % - AF488
Percentage of OM surface covered by LPS
Time post Kdo-alkyne / Kdo-N3 exchange (min)
E.
Time post Kdo-alkyne / Kdo-azide exchange (min)
0
15
30
45
60
75
90
105
120
Scale bar: 0.5 μm

**Extended Data Figure S2: Kdo-alkyne and Kdo-azide analogues were incorporated into the LPS of *E. coli* BW25113 cells at equivalent levels.** Representative DBSCAN outputs derived from Kdo-alkyne / Kdo-$N_3$ pulse–chase experiments in combination with two-colour 2D dSTORM super-resolution imaging demonstrate persistent spatial separation of newly inserted and background LPS-rich regions in the OM.

A. **Confocal microscopy of *E. coli* BW25113 cells metabolically labelled *in vivo* with either Kdo-azide (top row) or Kdo-alkyne (bottom row) reveals robust, equivalent levels of LPS fluorescent labelling achieved via $Cu^+$-catalysed azide-alkyne cycloaddition.** Representative widefield (AF488 fluorescence – left), brightfield (middle), and composite (right) images are shown. **Scale bars:** 20 µm.

B. **Quantitative FRAP analysis shows no significant differences in lateral mobility of LPS labelled with Kdo-azide (green, n = 30, median mobile fraction = 0.092) or Kdo-alkyne (purple, n = 30, median mobile fraction = 0.877).** Mobile fraction distributions are statistically indistinguishable (Mann–Whitney U = 461, $p$ = 0.877), indicating that incorporation of the functionalised Kdo-analogues does not affect LPS lateral mobility *in vivo*. Each data point represents the mobile fraction derived from a FRAP experiment carried out on an individual cell. Boxes show interquartile ranges and midline denotes median mobile fraction value.

C. **Fluorescent TSDS-PAGE analysis of AF488-labelled LPS confirms comparable click labelling levels of Kdo-azide (lane 2) and Kdo-alkyne (lane 3).** A commercially purchased O55:B5 AF488-LPS conjugate standard (lane 1) served as a molecular weight reference. **roLPS:** rough LPS; **deep roLPS:** deep rough LPS; **K12 roLPS:** rough LPS extracted from Kdo-analogue labelled *E. coli* K-12.

D. **Dye selection does not influence coverage by or overlap between background and newly inserted LPS in the OM of *E. coli* BW25113 cells.** Quantification of OM coverage and spatial overlap between background and new LPS clusters for both AF488-background LPS / AF647-newly inserted LPS and AF647-background LPS / AF488-newly inserted LPS (n ≥ 5 for each data point). Percentage of OM area occupied by DBSCAN-defined old and newly inserted LPS clusters over a 120-minute time course. Values represent mean ± standard error from individual cells (n ≥ 10 per timepoint). Well fitted ($R^2$ > 0.94 for all trendlines), near identical linear fits highlight the reciprocal loss and accumulation of old and newly inserted LPS, with rates of change being unaffected by dye selection.

E. **Representative DBSCAN clustering outputs from Kdo-alkyne, Kdo-azide pulse-chase experiments visualised via two-colour dSTORM showing spatial organisation of background (green) and newly inserted (purple) LPS in the OM of *E. coli* cells over a 120-minute monitoring period post Kdo-analogue exchange.** Background and newly inserted LPS rich regions remain discrete with consistently low levels of overlap (blue) across all time points, supporting a model of minimal mixing between old and newly synthesised LPS domains.

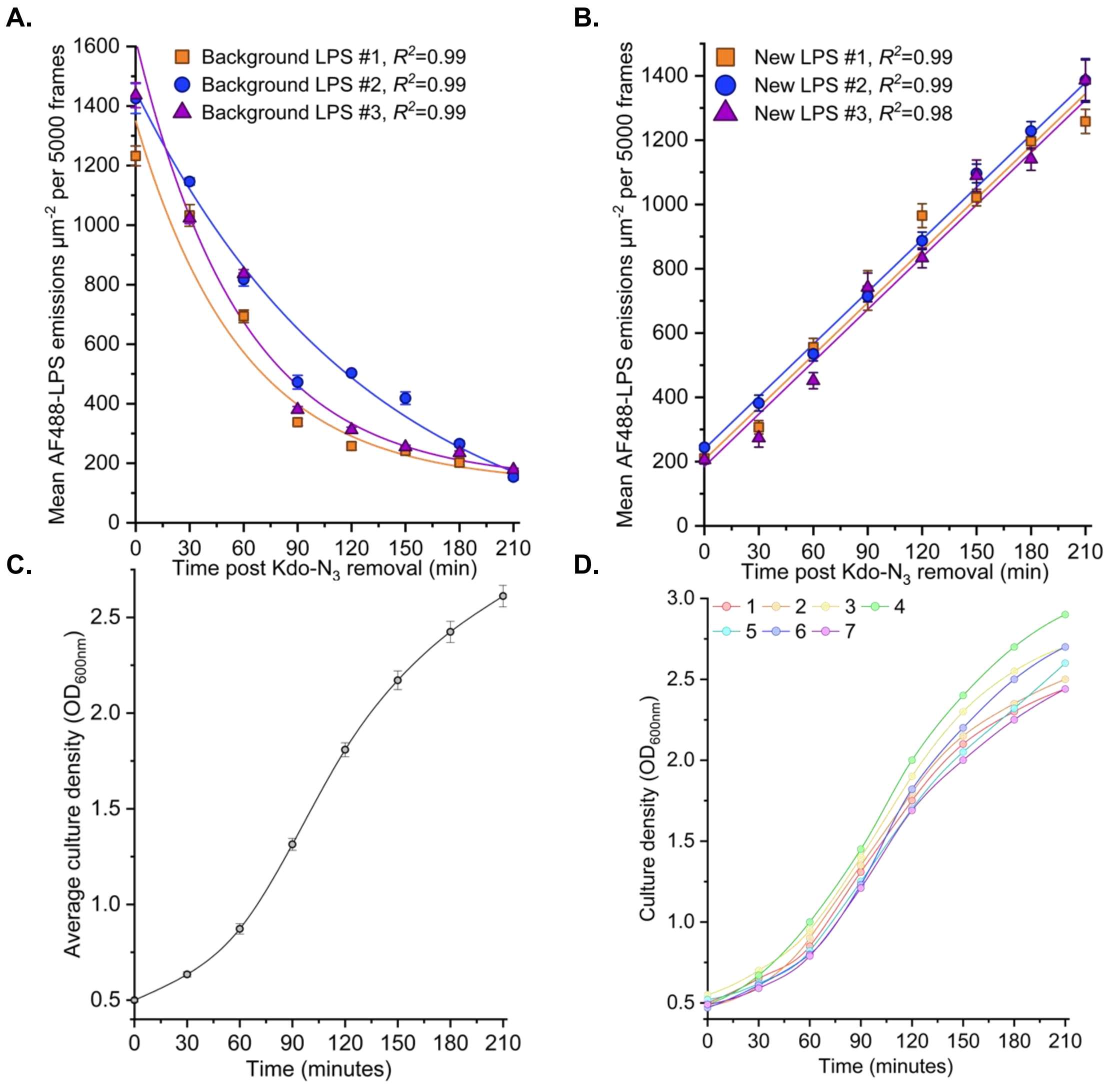


**Extended Data Figure S3: Quantitative analysis of background LPS loss, new LPS insertion, and growth rate during individual 210-minute Kdo-$N_3$ / native Kdo pulse–chase experimental replicates.**

**A. Per-cell background LPS abundance over time for each of three independent *E. coli* BW25113 replicates ($N = 3$; $n \geq 10$ cells per time point).** The number of discrete AF488–LPS localisations per outer membrane (OM), derived from dSTORM, is used as a proxy for background LPS content. Loss of background LPS continues throughout the 210-minute chase, even as cell division slows, and follows an exponential decay ($R^2 \geq 0.99$).

**B. Per-cell abundance of newly inserted LPS over time for each experimental replicate ($N = 3$; $n \geq 10$).** Newly inserted LPS was labelled using native Kdo / Kdo–$N_3$ metabolic pulse–chase and detected via AF488 fluorescence. Insertion proceeds linearly ($R^2 \geq 0.98$) over the full chase period, despite the onset of stationary phase between 120-minute and 150-minute time points.

**C. Mean growth curve ($n = 7$ replicates) showing culture $OD_{600}$ over the 210-minute chase periods.** A decrease in growth rate is evident beyond 120-minute time point, consistent with transition from late exponential to stationary phase.

**D. Individual $OD_{600}$ growth curves for all biological replicates used for the mean growth curve in panel C.** The source data for these growth curves are provided in Extended Data Table S22.

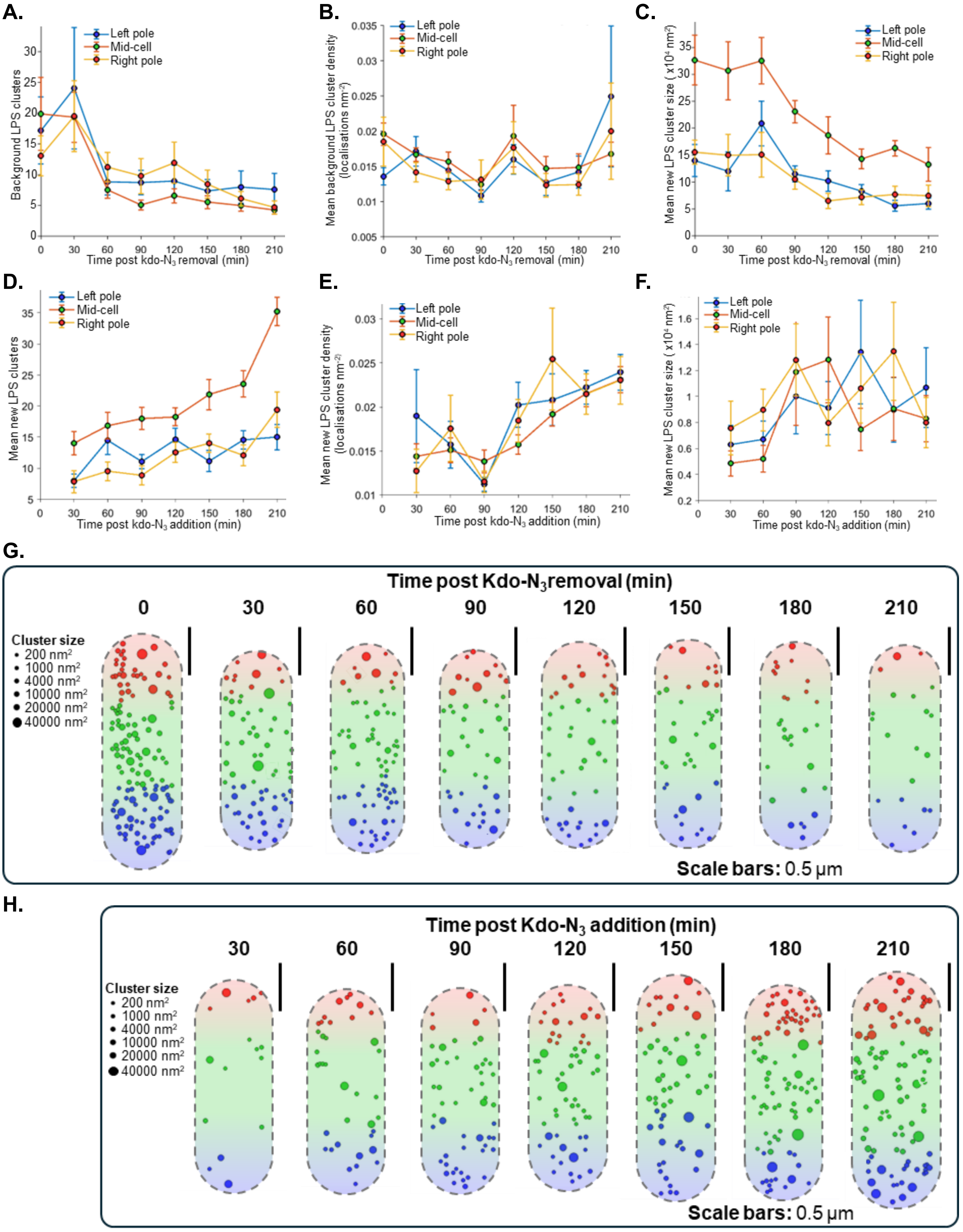

A.
Left pole
Mid-cell
Right pole
Background LPS clusters
Time post kdo-N3 removal (min)
B.
Mean background LPS cluster density (localisations nm-2)
Time post kdo-N3 removal (min)
C.
Mean new LPS cluster size ( x10^4 nm^2)
Time post kdo-N3 removal (min)
D.
Mean new LPS clusters
Time post kdo-N3 addition (min)
E.
Mean new LPS cluster density (localisations nm-2)
Time post kdo-N3 addition (min)
F.
Mean new LPS cluster size ( x10^4 nm^2)
Time post kdo-N3 addition (min)
G.
Time post Kdo-N3 removal (min)
0 30 60 90 120 150 180 210
Cluster size
200 nm2
1000 nm2
4000 nm2
10000 nm2
20000 nm2
40000 nm2
Scale bars: 0.5 μm
H.
Time post Kdo-N3 addition (min)
30 60 90 120 150 180 210
Cluster size
200 nm2
1000 nm2
4000 nm2
10000 nm2
20000 nm2
40000 nm2
Scale bars: 0.5 μm

**Extended Data Figure S4. Spatio-temporal analysis of LPS-rich clusters across outer membrane regions of *E. coli* revealed stable organisation of LPS-rich domains.**

**A–C.** Mean number, density, and size of background (AF488-labelled) LPS-rich clusters detected in polar (red and blue) vs. mid-cell (green) regions over time following Kdo-azide labelling.

**D–F.** Mean number, density, and size of newly inserted (AF488-labelled) LPS-rich clusters across the same regions over time following Kdo-azide pulse-labelling.

For all plots, n ≥ 10 cells per time point.

**G–H. Representative single-cell DBSCAN cluster maps visualised from dSTORM data for individual *E. coli* BW25113 cells.** Clusters are colour-coded by OM region (left pole: blue, mid-cell: green, right pole: red) and scaled globally by area. Images represent selected time points from background LPS turnover **(panel G)** and new LPS insertion **(panel H)** experiments. **Scale bars:** 0.5 μm

**A.**

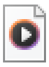

lower-OM-video.avi

**B.**

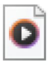

mid-cell-video.avi

**C.**

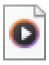

upper OM-video.avi

**D.**

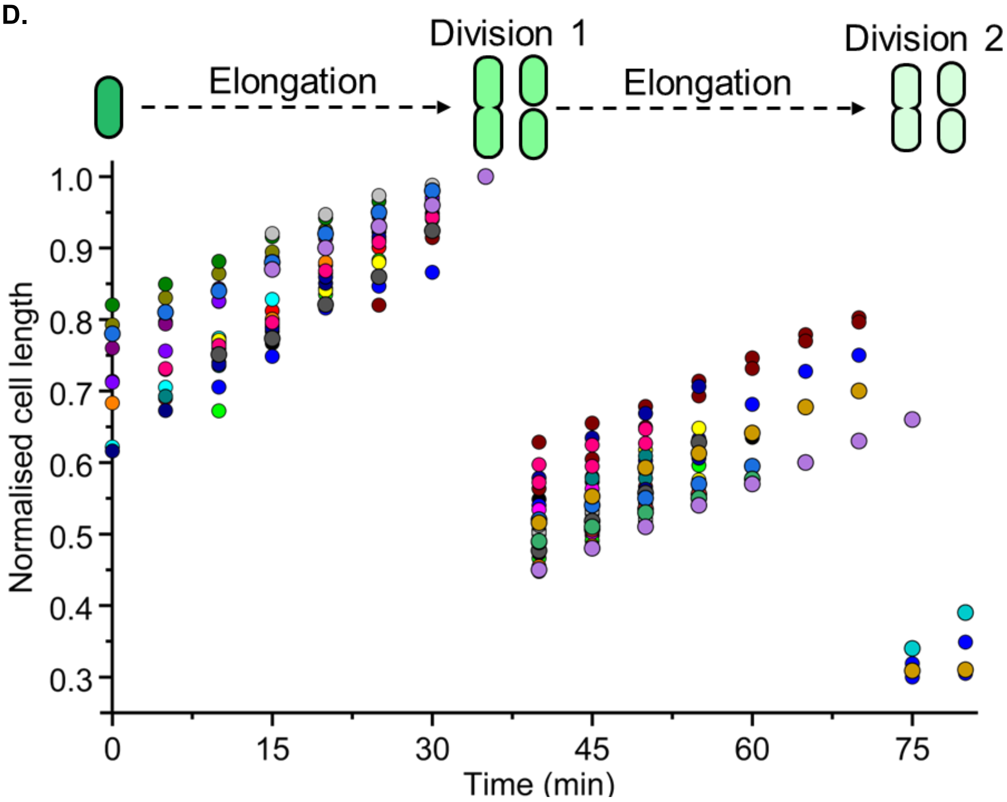


**Extended Data Figure S5: Real-time mapping of background LPS in live *E. coli* BW25113 cells via 3D SIM$^2$ at 37 °C, mounted on M9 CDM / 2% (w/v) agarose pads confirms LPS does not undergo binary partitioning, internalisation or cytoplasmic recycling.**

**A-C. Videos compiled from time-lapse 3D SIM$^2$ imaging experiment lasting 25 minutes showing changes in the distribution of background LPS in lower OM (*A.*), mid-cell (*B.*) and upper OM (*C.*) optical sections of live, growing and dividing *E. coli* BW25113 cells**. The clustered distributions of background LPS are maintained in the upper and lower OMs with no evidence of background LPS becoming segregated in the polar regions of the OM over time. No increase in background LPS fluorescent signal was observed within the cell boundaries at mid-cell depth indicating background LPS is not undergoing internalisation followed by cytoplasmic recycling. Kdo-$N_3$-containing LPS was fluorescently labelled via $Cu^+$-free strain promoted azide, alkyne cycloaddition ensuring cell viability was preserved. **Scale bars:** 5.0 µm

**D. Graph showing normalised changes in cell lengths (n = 30) observed over the course of the 3D SIM$^2$ experiment.** Cell lengths were normalised for each individual cell with the maximum length set to 1. An average doubling time of approximately 35 minutes was determined over the course of the time-lapse imaging experiment which matches the value measured in bulk bacterial cultures. This confirms cell elongation and division were not affected by Kdo-$N_3$ incorporation, SPAAC conjugation of DBCO-AF488, or exposure to the 488 nm excitation laser.

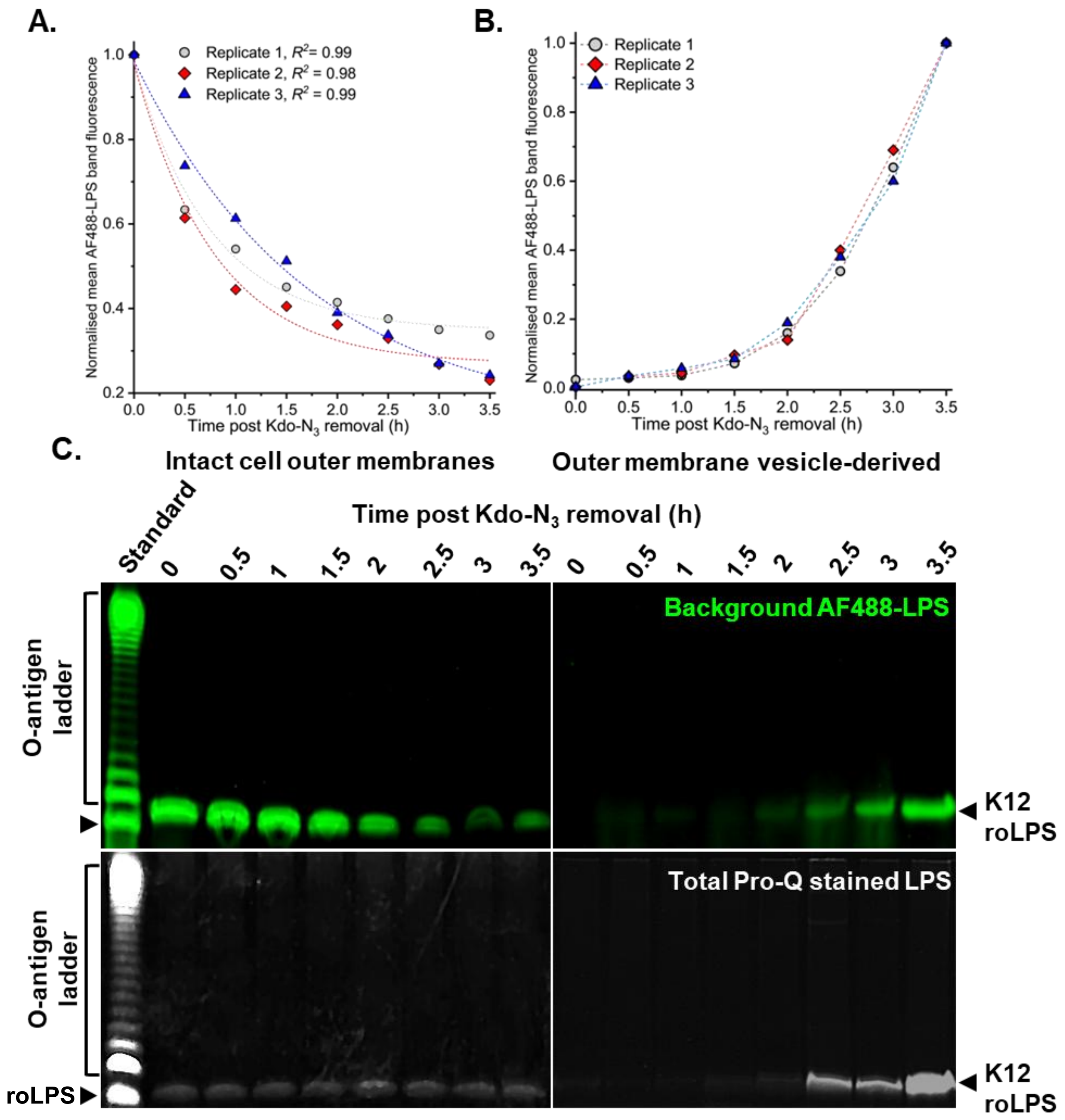


**Extended Data Figure S6: Outer membrane vesicles (OMVs) mediate clearance of background (pre-existing) LPS from the OM in non-growing, stationary phase *E. coli* BW25113 cells.**

**A–B.** Quantification of background AF488-labelled LPS over time in **(A)** intact-cell OM fractions and **(B)** OMV-derived fractions for individual experimental replicates. For intact-cell OM fractions, lanes contain a standardised mass of total LPS so that each lane corresponds to the LPS recovered from an equivalent number of cells at that time point. For OMV fractions, lanes contain the complete LPS extracted from the cell-free supernatant collected at each time point (standard culture volumes). Replicate datasets are well described by single-exponential functions ($R^2 \geq 0.98$), consistent with analogue partitioning of background LPS as expected during exponential growth but that continues into stationary growth phase.

**C.** TSDS-PAGE analysis showing decrease in background AF488-LPS (*top)* concentration in intact-cell OM fractions (*left*) and reciprocal increase in background AF488-LPS in OMV-derived fractions (*right*) with associated total Pro-Q stained total LPS loading controls (*bottom).* A commercially purchased O55:B5 AF488-LPS conjugate standard (lane 1, labelled 'standard') served as a molecular weight reference. **roLPS:** rough LPS. **K12 roLPS:** rough LPS extracted from *E. coli* BW25113 cells cultured in the presence of Kdo-azide or OMVs released by these cells and purified from cell-free culture supernatants.

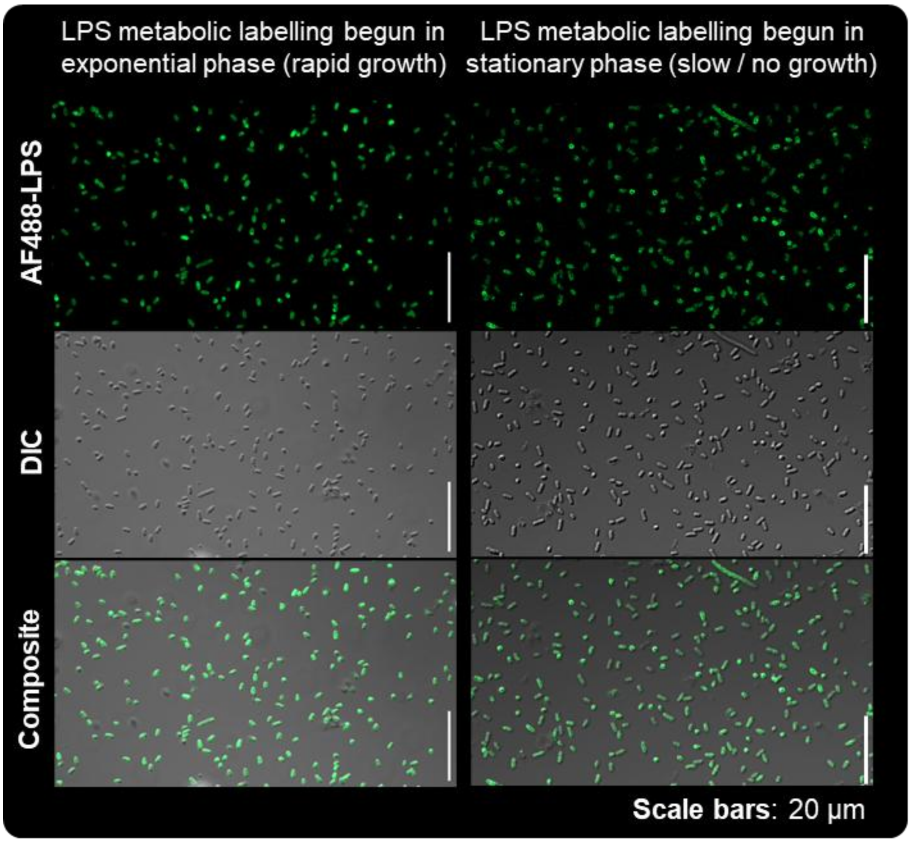


**Extended Data Figure S7: Kdo-$N_3$ incorporation in LPS of *E. coli* BW25113 cells is independent of cell growth phase and does not require cell elongation.** Equivalent levels of cell fluorescence were observed via fluorescence confocal microscopy in cells where Kdo-$N_3$ was added to the culture during the early stages of exponential growth ($OD_{600}$ = 0.05, *left*) or during early stationary phase ($OD_{600}$ = 2.5, *right*). After incubation for 12 h in M9 CDM charged with 4 mM Kdo-$N_3$, the LPS was labelled via CuAAC with AF488-alkyne. **Scale bars:** 20 µm.

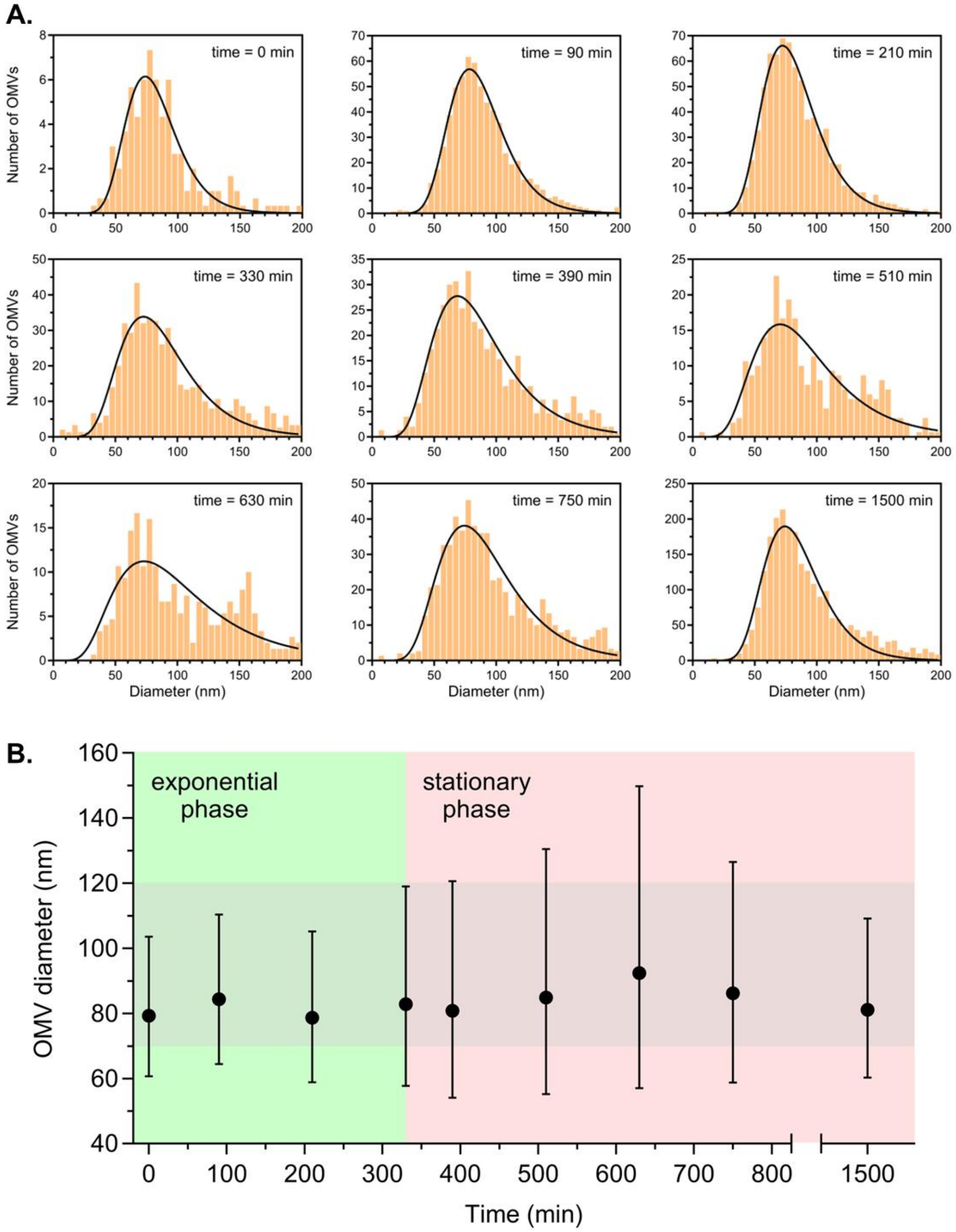
A.
time = 0 min
time = 90 min
time = 210 min
time = 330 min
time = 390 min
time = 510 min
time = 630 min
time = 750 min
time = 1500 min
Number of OMVs
Diameter (nm)
B.
exponential phase
stationary phase
OMV diameter (nm)
Time (min)

**Extended Data Figure S8. Characterisation of OMVs produced by *E. coli* during exponential and stationary phases of growth in chemically defined medium (CDM).**

**A.** Histograms of the diameters measured by nanoparticle tracking (NTA) for OMVs released by *E. coli* BW25113 cells cultured in M9 CDM at 37°C for different growth times (see top right corner of each histogram for growth time). Each histogram was compiled using data from technical replicates (N = 3) done on a representative sample. NTA data from the same technical replicates were analysed to obtain the OMV concentration in the bacterial culture (see Figure 6F). The skewed size histograms at each timepoint were best described by a log-normal distribution function with mostly smaller vesicle sizes observed at each time point. Non-linear regression of the histograms with this distribution function yielded the following $R^2$ values (listed in chronological order): 0.69, 0.96, 0.96, 0.78, 0.60, 0.58, 0.50, 0.75 and 0.90.

**B. The mean OMV diameters (± SD) obtained from the analysis of the size histogram for each time point**. The grey shaded region indicates the predicted range of OMV diameters (*d* = 73-122 nm) from the OM buckling model for a range of experimentally determined OM thicknesses (= 7.5-21 nm, see Methods section for mathematical description of the OM buckling model).

**A.**

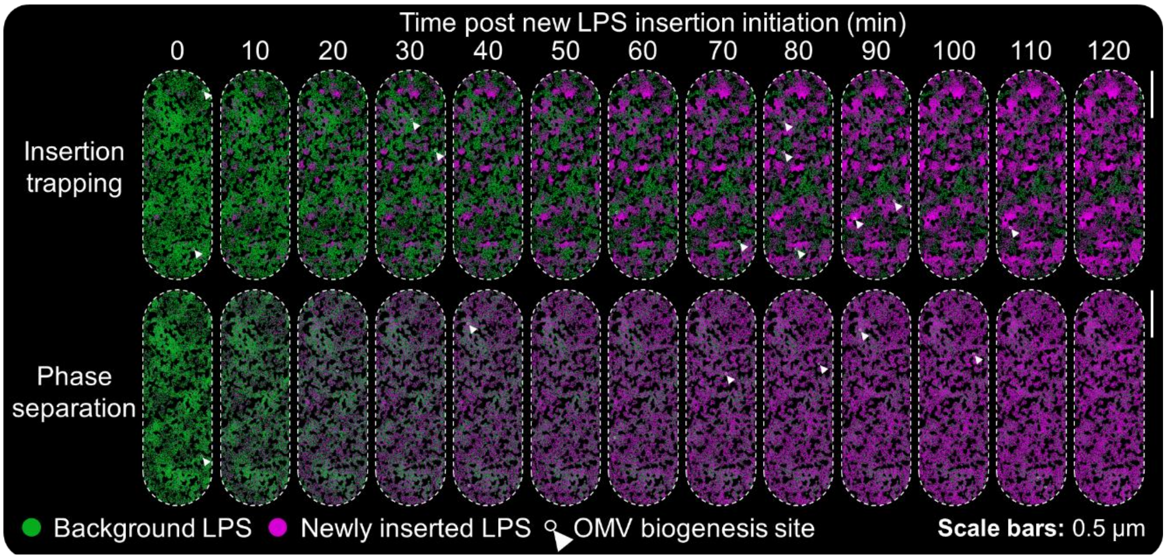


**B.**

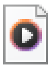

Video_Insertion trapping_Restricted

**Extended Data Figure S9. Particle-based simulations of the outer leaflet in the *E. coli* OM for two different organisation models.**

A. **Extended snapshot sequences (0-120 min, 10 min intervals) from the two organisation models.** **Insertion-trapping model (top):** New LPS is delivered in bursts at discrete hotspots (LptD-like sites) and lateral motion of both new and background LPS is strongly confined, resulting in new patches remaining localised whilst background LPS is depleted locally by OMV formation.
**Phase separation model (bottom):** A short-range 'like–like' coupling constant favours LPS-LPS contacts irrespective of old or new LPS type and drives domain growth under otherwise identical LPS turnover rates. This short-range favourable interaction tests whether old-new LPS co-clustering could reproduce the experimental observations. **Magenta:** Newly inserted LPS. **Green:** Background LPS. **Black / vacant spaces:** LPS-inaccessible OMP-occupied regions (~25% area). **White arrowhead / circles:** OMV biogenesis events arising in LPS-dense regions according to the OM buckling model. **Scale bars:** 0.5 µm**.**

B. **Video sequence comparing changes in newly inserted and background LPS distributions and abundances in the OM of *E. coli* for the insertion-trapping simulation (model 1, left) and phase separation simulation (model 2, right).** Video is composed of simulation snapshots at 5 min intervals from 0 – 120 min post-initiation of new LPS insertion**.** **Magenta:** Newly inserted LPS. **Green:** Background LPS. **Black / vacant spaces:** LPS-inaccessible OMP-occupied regions (as described for panel A). **White circles:** OMV biogenesis events arising in LPS-dense regions according to the OM buckling model. **Scale bars:** 0.5 µm.

### Extended Data Tables

**Extended Data Table S1.** Details of Gram-negative bacterial strains used in this study

| Strain | Organism | Selection antibiotic | Notes | Ref. |
|---|---|---|---|---|
| **K-12 substr. BW25113** | *E. coli* | None | Parent strain of Keio collection. *lacI$^q$ rrnB$_{T14}$ ΔlacZ$_{WJ16}$ hsdR514 ΔaraBAD$_{AH33}$ ΔrhaBAD$_{LD78}$* | 1 |
| **BW25113 pBAD-lptD$^*$** | *E. coli* | 100 µg mL$^{-1}$ AMP 35 µg mL$^{-1}$ CHL | BW25113 strain co-transformed with pBAD-lptD$^*$ and pEVOL-pylRS plasmids. Produces recombinant LptD with ncAA propargyl-L-Lysine (Lysine analogue) at position D600 when provided with ncAA resulting from leaky *lptD$^*$* expression (L-arabinose was omitted to prevent over-production of LptD$^*$). | 1 |
| **Δ*ompA* pBAD-ompA*** | *E. coli* | 100 µg mL$^{-1}$ AMP 35 µg mL$^{-1}$ CHL 30 µg mL$^{-1}$ KAN | Keio collection strain JW0940-KC with *ompA* gene replaced by kanamycin resistance cassette and transformed with pBAD-ompA* and pEVOL-pylRS plasmids. Produces recombinant OmpA with ncAA propargyl-L-Lysine (Lysine analogue) at position E89 when provided with the ncAA resulting from leaky *ompA$^*$* gene expression (L-arabinose was omitted to prevent over-production of OmpA$^*$). | 2 |

**Antibiotic abbreviations: KAN =** Kanamycin (Sigma-Aldrich), **AMP =** Ampicillin (Melford Laboratories Ltd.), **CHL =** Chloramphenicol (Sigma-Aldrich)

**Extended Data Table S2.** DNA primer sequences

| Primer ID | Sequence (5' → 3') | Notes |
|---|---|---|
| lptD$^{WT}$-for | TTTGGGCTAACAGGAGGAATTACATATGAAAAAACGTATCCCCACTCTCC | *lptD* gene PCR amplification |
| lptD$^{WT}$-rev | GAGATGAGTTTTTGTTCTAGAAAGCTTACAAAGTGTTTTGATACGGCAGA | *lptD* gene PCR amplification |
| pBADcLIC-for | TCTGCCGTATCAAAACACTTTGTAAGCTTTCTAGAACAAAAACTCATCTC | pBADcLIC vector PCR amplification |
| pBADcLIC-rev | GGAGAGTGGGGATACGTTTTTTCATATGTAATTCCTCCTGTTAGCCCAAA | pBADcLIC vector PCR amplification |
| pBADcLIC-forS | ATGCCATAGCATTTTTATCC | pBAD-lptD sequencing |
| pBADcLIC-revS | GATTTAATCTGTATCAGG | pBAD-lptD sequencing |
| lptDmid-forS | TACTTTGAGTTCTACCTGCC | pBAD-lptD sequencing |
| lptDmid-revS | ATGCTGGAGTTACTGGTCGC | pBAD-lptD sequencing |
| lptD$^{[D600]}$-forM | CGATGACAACATAACATGGGAGAATTAGGACAAAACGGGTTCACTGGT | pBAD-lptD PCR mutagenesis |
| lptD$^{[D600]}$-revM | ACCAGTGAACCCGTTTTGTCCTAATTCTCCCATGTTATGTTGTCATCG | pBAD-lptD PCR mutagenesis |
| ompA$^{WT}$-for | GCTAACAGGAGGAATTAACCATGGATGAAAAAGACAGCTATCGCGATTG | *ompA* gene PCR amplification |
| ompA$^{WT}$-rev | GATGAGTTTTTGTTCTAGAAAGCTTCGTTAAGCCTGCGGCTGAGTTAC | *ompA* gene PCR amplification |
| pBAD-for-2 | CGTTGTAACTCAGCCGCAGGCTTAACGAAGCTTTCTAGAACAAAAACTCATCTCAG | pBADcLIC vector PCR amplification |
| pBAD-rev-2 | CAATCGCGATAGCTGTCTTTTTCATCCATGGTTAATTCCTCCTGTTAGCC | pBADcLIC vector PCR amplification |
| ompA$^{[E89]}$-for | CGTATGCCGTACAAAGGCAGCGTTTAGAACGGTGCATACAAAGCTCAGGGC | pBAD-ompA PCR mutagenesis |
| ompA$^{[E89]}$-rev | GCCCTGAGCTTTGTATGCACCGTTCTAAACGCTGCCTTTGTACGGCATACG | pBAD-ompA PCR mutagenesis |

**Extended Data Table S3.** Composition of supplemented M9 chemically defined medium (pH 7.2)

| **Component** | **Concentration** |
|---|---|
| $Na_2HPO_4$ | 48.0 mM |
| $KH_2PO_4$ | 22.0 mM |
| NaCl | 8.6 mM |
| D-glucose | 0.4% (w/v) |
| $NH_4Cl$ | 1.0 g $L^{-1}$ |
| Casamino acids | 0.05% (w/v) |
| $FeSO_4$ | 0.1 mM |
| $MgSO_4$ | 2.0 mM |
| $CaCl_2$ | 0.1 mM |

**Extended Data Table S4.** Final concentrations of alkyne functionalised dyes (Click Chemistry Tools, Vector Laboratories, Inc.) used in CuAAC ‘Click-iT’ reaction mixtures

| **AFDye** | **Final concentration** (μM) |
|---|---|
| AF488 | 10 |
| AF647 | 50 |

**Extended Data Table S5.** Final concentration of AF488-DBCO (Click Chemistry Tools, Vector Laboratories, Inc.) used in SPAAC reaction mixtures

| **AFDye DBCO type** | **Final concentration** (μM) |
|---|---|
| AF488 | 25 |

**Extended Data Table S6.** LptD* and newly inserted, Kdo-$N_3$-LPS dual labelling combinations used to confirm fluorescent dye type and labelling order did not influence two-colour dSTORM results

| **First labelled species / fluorescent dye** | **Second labelled species / fluorescent dye** |
|---|---|
| LptD* / AF488-azide | Newly inserted LPS / AF647-alkyne |
| Newly inserted LPS / AF647-alkyne | LptD* / AF488-azide |
| LptD* / AF647-azide | Newly inserted LPS / AF488-alkyne |
| Newly inserted LPS / AF488-alkyne | LptD* / AF647-azide |

**Extended Data Table S7.** Components in glucose oxidase (GluOx) / catalase (Cat) oxygen scavenging buffer system

| **Component** | **Volume** (μL) |
|---|---|
| Degassed deionised $H_2O$ | 895 |
| PBS pH 7.4 [40 x] (BioStatus Ltd.) | 25 |
| GluOx [10 mg $mL^{-1}$] | 20 |
| Cat [2 mg $mL^{-1}$] | 20 |
| Glucose [300 mg $mL^{-1}$] | 40 |

**Extended Data Table S8.** Additional settings for dSTORM image acquisition with Zeiss Elyra 7 super-resolution imaging system

| **Dye** | **Laser beam splitter** | **Laser** | **Emission filter** (nm) | **xy scaling** (µm pixel$^{-1}$) | **TIRF mirror angle** | **TIRF collimator** |
|---|---|---|---|---|---|---|
| AF488 | 405/488/561/642 | 488 nm DPSS | BP420-480 + BP490-550 | 0.097 | 60° | 230 |
| AF647 | 405/488/561/642 | 561 nm DPSS | BP570-620 + LP655 | 0.097 | 60° | 400 |

**Extended Data Table S9.** Typical dSTORM image filtering settings

| **Filter** | **AF488 dye** | **AF647 dye** |
|---|---|---|
| Precision (nm) | 1 – 40 | 1 – 40 |
| Number of photons | 1 – 1750 | 1 – 1250 |
| Point squared function half width (nm) | 60 – 200 | 100 – 300 |
| Background variance | 1 – 300 | 1 – 300 |
| Chi-squared | 0.4 – 1.2 | 0.4 – 1.2 |

**Extended Data Table S10:** Mean LptD* vs newly inserted LPS cross-correlation function data derived from analysis of dual-colour dSTORM outputs for individual cells (n = 30, N = 3)

| **Radius (nm)** | **Mean g*(r)* (n = 30, N = 3)** | **Standard Error [Mean g(r)]** |
|---|---|---|
| 30 | 4.37 | 0.51 |
| 60 | 3.98 | 0.40 |
| 90 | 3.43 | 0.27 |
| 120 | 2.91 | 0.18 |
| 150 | 2.49 | 0.12 |
| 180 | 2.13 | 0.09 |
| 210 | 1.84 | 0.07 |
| 240 | 1.63 | 0.06 |
| 270 | 1.45 | 0.05 |
| 300 | 1.30 | 0.04 |
| 330 | 1.21 | 0.04 |
| 360 | 1.11 | 0.04 |
| 390 | 1.03 | 0.04 |
| 420 | 0.96 | 0.04 |
| 450 | 0.92 | 0.04 |
| 480 | 0.87 | 0.04 |
| 510 | 0.83 | 0.04 |
| 540 | 0.81 | 0.04 |
| 570 | 0.78 | 0.04 |
| 600 | 0.75 | 0.04 |
| 630 | 0.73 | 0.05 |

**Extended Data Table S11:** Results from Mann-Whitney t-test assessing the significance of variation between LptD* vs newly inserted LPS Pearson correlation coefficient (PCC) values for polar, mid-cell and entire OM regions of *E. coli* BW25113 cells imaged via two-colour dSTORM after simultaneous dual fluorescent labelling of LptD* and newly inserted LPS.

| Condition 1 | Condition 2 | $n_1$ | $n_2$ | Median $PCC_1$ | Median $PCC_2$ | *p* | U | Z |
|---|---|---|---|---|---|---|---|---|
| Left pole | Mid-cell | 30 | 30 | 0.49 | 0.34 | 0.32 | 518 | 0.99 |
| Left pole | Right pole | 30 | 30 | 0.49 | 0.52 | 0.46 | 399 | -0.75 |
| Left pole | Entire OM | 30 | 30 | 0.49 | 0.37 | 0.24 | 530 | 1.18 |
| Right pole | Mid-cell | 30 | 30 | 0.52 | 0.34 | 0.10 | 561 | 1.63 |
| Right pole | Entire OM | 30 | 30 | 0.52 | 0.37 | 0.07 | 574 | 1.83 |
| Mid-cell | Entire OM | 30 | 30 | 0.34 | 0.37 | 0.97 | 447 | -0.04 |

**Extended Data Table S12.** Background, Kdo-alkyne-LPS and newly inserted, Kdo-$N_3$-LPS dual labelling combinations used to ensure fluorescent dye type and labelling order did not influence two-colour dSTORM results

| First labelled species / fluorescent dye | Second labelled species / fluorescent dye |
|---|---|
| Newly inserted LPS / AF488-alkyne | Background LPS / AF647-azide |
| Background LPS / AF647-azide | Newly inserted LPS / AF488-alkyne |
| Newly inserted LPS / AF647-alkyne | Background LPS / AF488-azide |
| Background LPS / AF488-azide | Newly inserted LPS / AF647-alkyne |

**Extended Data Table S13.** Average background and newly inserted LPS cluster overlap and normalised overlap new / background LPS cluster surface area

| Time post Kdo-analogue exchange (min) | Average background LPS % | S.E. | Average new LPS % | S.E. | Overlap cluster area / (new + old cluster area) | S.E. |
|---|---|---|---|---|---|---|
| 0 | 54.48 | 6.86 | 0.26 | 0.07 | 0.0027 | 0.0008 |
| 15 | 45.31 | 4.32 | 7.24 | 1.79 | 0.0635 | 0.0154 |
| 30 | 37.29 | 4.12 | 13.88 | 2.14 | 0.0977 | 0.0153 |
| 45 | 32.11 | 1.90 | 19.64 | 1.59 | 0.1280 | 0.0201 |
| 60 | 22.28 | 3.09 | 21.34 | 3.04 | 0.1440 | 0.0212 |
| 75 | 18.10 | 3.12 | 32.52 | 3.99 | 0.1281 | 0.0205 |
| 90 | 15.63 | 2.51 | 37.04 | 3.46 | 0.1164 | 0.0185 |
| 105 | 11.09 | 1.89 | 41.76 | 4.04 | 0.1051 | 0.0159 |
| 120 | 5.95 | 0.89 | 47.73 | 6.73 | 0.0869 | 0.0125 |
| **Ref:** Figure 2E | | | | | **Ref:** Figure 2F | |

**Extended Data Table S14.** Composition of 4x TSDS-PAGE sample loading buffer

| Component | Final concentration |
|---|---|
| Tris-Cl pH 6.8 | 0.25 M |
| Sodium dodecyl sulphate (SDS) | 227 mM |
| Bromophenol blue | 0.02 % (w/v) |
| Glycerol | 4.3 M |

**Note:** β-ME added to a final concentration of 2% (v/v) immediately before use.

**Extended Data Table S15.** AF488 emission dSTORM image filtering settings

| Filter | AF488 value range |
|---|---|
| Precision / nm | 1 – 40 |
| Number of photons | 1 – 1500 |
| Point spread function half width / nm | 60 – 220 |
| Background variance | 1 – 300 |
| Chi squared | 0.4 – 1.2 |

**Extended Data Table S16:** Mean background LPS turnover rates in the outer membrane of *E. coli* BW25113 cells derived from dSTORM analyses during Kdo-$N_3$ / native Kdo pulse-chase experiments. Data show the mean AF488–LPS emission density ($\mu m^{-2}$) and standard error (S.E.) at 30-minute time intervals following removal of Kdo–$N_3$ from the culture medium. Each experimental replicate (N = 3) includes ≥10 individual cells per time point. In all replicates, background LPS levels decay exponentially over time. The rate of turnover does not appear to be altered by the transition from late exponential to stationary phase.

| **Experimental replicate** | **1.** | | **2.** | | **3.** | |
|---|---|---|---|---|---|---|
| **Time post Kdo-$N_3$ removal / h** | **Mean AF488-LPS emissions $\mu m^{-2}$** | **S.E.** | **Mean AF488-LPS emissions $\mu m^{-2}$** | **S.E.** | **Mean AF488-LPS emissions $\mu m^{-2}$** | **S.E.** |
| 0.0 | 1232.39 | 33.21 | 1425.17 | 50.28 | 1436.30 | 42.03 |
| 0.5 | 1032.55 | 36.27 | 1146.42 | 12.07 | 1022.91 | 17.80 |
| 1.0 | 693.62 | 21.23 | 817.34 | 22.27 | 835.98 | 14.37 |
| 1.5 | 337.77 | 11.71 | 472.33 | 23.17 | 380.13 | 10.87 |
| 2.0 | 257.49 | 12.80 | 503.20 | 6.63 | 312.24 | 8.67 |
| 2.5 | 241.37 | 8.25 | 418.62 | 21.09 | 254.52 | 6.48 |
| 3.0 | 201.97 | 9.49 | 265.33 | 9.85 | 234.98 | 6.32 |
| 3.5 | 164.07 | 5.15 | 153.80 | 9.56 | 178.45 | 5.15 |

**Extended Data Table S17:** Mean LPS insertion rates in the OM of *E. coli* BW25113 cells derived from dSTORM analyses during native Kdo / Kdo-$N_3$ pulse-chase experiments. Data show the mean AF488–LPS emission density ($\mu m^{-2}$) and standard error (S.E.) at 30-minute time intervals following addition of Kdo–$N_3$ to the culture medium. Each experimental replicate (N = 3) includes ≥10 individual cells per time point. In all replicates, new LPS levels increase in a linear manner over time. The rate of new LPS insertion does not appear to be altered by the transition from late exponential to stationary phase.

| **Experimental replicate** | **1.** | | **2.** | | **3.** | |
|---|---|---|---|---|---|---|
| **Time post Kdo-$N_3$ addition / h** | **Mean AF488-LPS emissions $\mu m^{-2}$** | **S.E.** | **Mean AF488-LPS emissions $\mu m^{-2}$** | **S.E.** | **Mean AF488-LPS emissions $\mu m^{-2}$** | **S.E.** |
| 0.0 | 209.85 | 8.13 | 244.28 | 7.08 | 206.01 | 13.22 |
| 0.5 | 306.97 | 20.91 | 382.12 | 24.63 | 273.46 | 28.30 |
| 1.0 | 555.61 | 27.56 | 535.04 | 21.62 | 451.67 | 25.19 |
| 1.5 | 732.44 | 61.67 | 714.16 | 16.45 | 741.65 | 44.98 |
| 2.0 | 965.05 | 36.97 | 887.01 | 27.06 | 833.49 | 30.79 |
| 2.5 | 1021.37 | 25.68 | 1096.33 | 29.17 | 1089.09 | 49.11 |
| 3.0 | 1196.48 | 24.91 | 1228.20 | 29.30 | 1140.78 | 34.65 |
| 3.5 | 1258.06 | 37.57 | 1385.65 | 67.81 | 1386.24 | 63.32 |

**Extended Data Table S18.** Summary of DBSCAN statistics for background LPS

| Time (mins) | Mean cluster number per µm² of OM | Mean cluster surface area (nm²) | Cluster density (localisations µm⁻²) |
|---|---|---|---|
| 0 | 24 | 17578 | 9419.57 |
| 30 | 19 | 20374 | 20373.65 |
| 60 | 19 | 8749 | 8748.68 |
| 90 | 14 | 7119 | 7119.06 |
| 120 | 13 | 6537 | 6537.44 |
| 150 | 13 | 6755 | 6754.92 |
| 180 | 12 | 6968 | 6967.52 |
| 210 | 12 | 5122 | 5121.69 |

**Extended Data Table S19.** Summary of DBSCAN statistics for newly inserted LPS

| Time (mins) | Mean cluster number per µm² of OM | Mean cluster surface area (nm²) | Cluster density (localisations µm⁻²) |
|---|---|---|---|
| 30 | 14 | 5962 | 15182.84 |
| 60 | 14 | 6619 | 15876.97 |
| 90 | 16 | 10558 | 12502.23 |
| 120 | 20 | 10295 | 14765.60 |
| 150 | 24 | 9820 | 21430.04 |
| 180 | 30 | 10095 | 21730.29 |
| 210 | 33 | 9724 | 23264.76 |

**Extended Data Table S20:** Additional settings for lattice SIM image acquisition with Zeiss Elyra 7 super-resolution imaging system

| Laser beam splitter | Laser | Emission filter (nm) | laser power (%) | Exposure (ms) | *xy* scaling (µm) | *z* scaling (µm) | Grating period (nm) | Phases |
|---|---|---|---|---|---|---|---|---|
| 405/488/561 /642 | 488 nm DPSS | BP420 – 480 + BP490 - 550 | 3 | 50 | 0.031 | 0.110 | 718.33 | 15 |

**BP:** Band pass. **LP:** Long Pass. **DPSS:** Diode pumped solid state. Images collected using a Plan-Apochromat 63x /1.4 (DIC M27) oil immersion objective lens. **Image size:** 10 µm x 10 µm x 2 µm, 16-bit

**Extended Data Table S21:** OMV lysis buffer composition. Prepared with HPLC-grade water. β-Mercaptoethanol added to a final concentration of 2% (v/v) immediately before use.

| Component | Concentration |
|---|---|
| Sodium dodecyl sulfate (SDS) | 2% (w/v) |
| Tris-Cl pH 6.5 | 60 mM |
| EDTA | 1 mM |

**Extended Data Table S22:** $OD_{600nm}$ measurements for *E. coli* BW25113 cells cultured in M9 CDM during Kdo-$N_3$ / native Kdo pulse-chase experiments. Each replicate adheres to a typical growth curve with cultures transitioning from late exponential to stationary phase growth between time points at 120 and 150 min.

| Time (min) | Rep. 1 $OD_{600}$ | Rep. 2 $OD_{600}$ | Rep. 3 $OD_{600}$ | Rep. 4 $OD_{600}$ | Rep. 5 $OD_{600}$ | Rep. 6 $OD_{600}$ | Rep. 7 $OD_{600}$ |
|---|---|---|---|---|---|---|---|
| 0 | 0.50 | 0.48 | 0.55 | 0.49 | 0.52 | 0.47 | 0.49 |
| 30 | 0.65 | 0.605 | 0.7 | 0.67 | 0.62 | 0.61 | 0.59 |
| 60 | 0.85 | 0.9 | 0.95 | 1.00 | 0.82 | 0.80 | 0.79 |
| 90 | 1.31 | 1.35 | 1.4 | 1.45 | 1.25 | 1.23 | 1.21 |
| 120 | 1.75 | 1.80 | 1.90 | 2.00 | 1.70 | 1.82 | 1.69 |
| 150 | 2.10 | 2.15 | 2.30 | 2.40 | 2.05 | 2.20 | 2.00 |
| 180 | 2.30 | 2.35 | 2.55 | 2.70 | 2.32 | 2.50 | 2.25 |
| 210 | 2.44 | 2.50 | 2.70 | 2.90 | 2.60 | 2.70 | 2.44 |

**Extended Data Table S23:** Concentration of viable bacterial cells in culture as a function of time. The colony forming units (CFU) were calculated for each time point using serial dilutions of the bacterial culture and the plate count method. Bacterial cells were cultured in supplemented M9 CDM at 37°C. The two separate bacterial cultures (ID numbers 1 and 2) used to determine CFU were grown under identical conditions.

| Time (min) | Growth phase | Culture ID number | $OD_{600}$ | CFU / ml in original culture (x $10^9$) | (CFU/ml) / $OD_{600}$ (x $10^8$) |
|---|---|---|---|---|---|
| 0 | --- | 1 | 0.150 | 0.0424 | 2.83 |
| 60 | exponential | 1 | 0.309 | 0.032 | 1.04 |
| 90 | exponential | 1 | 0.441 | 0.096 | 2.18 |
| 150 | exponential | 1 | 0.696 | 0.142 | 2.05 |
| 210 | exponential | 1 | 1.34 | 0.296 | 2.21 |
| 270 | late exponential | 1 | 2.47 | 0.592 | 2.40 |
| 330 | late exponential | 1 | 3.07 | 0.752 | 2.45 |
| 390 | stationary | 1 | 3.05 | 1.104 | 3.62 |
| 450 | stationary | 1 | 3.23 | 1.072 | 3.32 |
| 510 | stationary | 2 | 2.94 | 0.928 | 3.16 |
| 570 | stationary | 2 | 3.05 | 0.944 | 3.10 |
| 630 | stationary | 2 | 2.99 | 0.912 | 3.05 |
| 690 | stationary | 2 | 2.95 | 1.048 | 3.55 |
| 1500 | stationary | 1 | 3.21 | 1.176 | 3.66 |
