## Extended Methods - referenced where relevant in manuscript for "Persistent LPS insertion, spatial segregation and vesicle biogenesis drive growth-independent adaptation of the *Escherichia coli* outer membrane"

**Bacterial strains and plasmids**

Details of strains used in this study are provided in ***Extended Data Table S1***. The Keio collection mutant of *Escherichia coli* K12 subsp. BW25113 used in this work (Δ*ompA*) was validated previously via PCR, confirming the presence of kanamycin resistance cassette within the open-reading frame of interest [1-3].

pBAD-lptD* and pBAD-ompA* plasmids were produced via Gibson isothermal assembly [4]. PCR was used to amplify the *lptD* and ompA gene inserts ***(see Extended Data Table S2 for primers)*** from purified *E. coli* K12 subsp. MG1655 genomic DNA. The amplified genes were cloned into pBADcLIC vectors using a commercial Gibson isothermal assembly kit (NEBuilder® HiFi DNA Assembly). The pBADcLIC vector system was selected because its *araBAD* promoter provides tight regulation yet permits low-level basal (‘leaky’) expression in the absence of inducer (L-arabinose), allowing sufficient outer membrane protein production while minimising toxicity and misfolding that can occur under the strong induction conditions typical of IPTG-inducible recombinant expression systems. PCR site-directed mutagenesis ***(Extended Data Table S2)*** was then used to incorporate the amber stop codon TAG in place of the codon for the D600 residue in *lptD* ***(Extended Data Fig. S1B)*** and E89 residue in *ompA*. Successful cloning and mutagenesis were verified via DNA sequencing (Eurofins Genomics LLC). The pEVOL-pylRS plasmid, encoding the Pyrrolysyl-tRNA synthetase/pyrrolysyl-tRNA pair from *Methanosarcina mazei* was obtained from Dr. Edward Lemke (EMBL Heidelberg) under MTA [5,6].

pBAD-lptD* and pEVOL-pylRS were co-transformed into *E. coli* BW25113 via electroporation (MicroPulser, Bio-Rad Laboratories) (1.8 kV, 25 µF, time constant: 3.5 ms). *E. coli* BW25113 cells co-transformed with pBAD-lptD* and pEVOL-pylRS were selected by spreading some of the electroporated culture on an Lysogeny Broth (Miller formulation, LB-Miller) agar plate containing 100 µg $mL^{-1}$ ampicillin and 35 µg $mL^{-1}$ chloramphenicol and incubating the plate overnight at 37 °C. Unless otherwise specified, the incubation of bacterial cultures was done at 37 °C with shaking (220 rpm) in a 50 mL screw-cap polypropylene tube. Liquid pre-cultures were prepared by inoculating 5 mL of supplemented M9 chemically defined medium (M9 CDM, ***Extended Data Table S3)*** with appropriate antibiotics ***(Extended Data Table S1)*** using a well-isolated single colony of the desired strain picked from a freshly streaked LB-Miller agar plate. Post-inoculation cultures were incubated for 6 – 8 h or until cell densities reached a minimum optical density at 600 nm ($OD_{600}$) of 1.5.

Following plasmid purification (Qiagen© Midi plasmid purification kit) and DNA sequencing (Eurofins Genomics LLC), the TAG-containing pBAD-ompA vector (pBAD-ompA*) and pEVOL-pylRS were co-transformed into *E. coli* Δ*ompA* cells via electroporation as above. Co-transformed cells were selected using LB-Miller agar plates with 30 µg $mL^{-1}$ kanamycin, 100 µg $mL^{-1}$ ampicillin and 35 µg $mL^{-1}$ chloramphenicol. A single colony from this plate was used to inoculate a volume of supplemented M9 CDM with kanamycin, ampicillin and chloramphenicol antibiotics. Following prolonged pre-culture incubation, the appropriate volume of pre-culture was used to inoculate a fresh batch of supplemented M9 CDM with ampicillin and chloramphenicol antibiotics to an $OD_{600}$ of 0.05. The cultures were then incubated for 2 – 3 hours.

**Production of 8-Azido-3,8-dideoxy-D-manno-octulosonic acid (Kdo-$N_3$)**

Kdo-$N_3$ was synthesised and purified following established protocols[7] .The synthesised Kdo-$N_3$ was utilised for all super-resolution fluorescence microscopy experiments (dSTORM and 3D SIM[2]) and yielded the same results as commercially available Kdo-$N_3$ (Click Chemistry Tools, Vector Laboratories, Inc.) used in more recent work. Freeze-dried stocks of the synthesised compound and the purchased compound were resuspended to a concentration of 400 mM in sterile, deionised water and stored at -20 °C until required.

**Production of 8-N-(pent-4-ynamido)-3,8-dideoxy-N-D-manno-octulosonic acid (Kdo-alkyne)**

Kdo-alkyne was synthesised and purified following established protocols from commercially available Kdo-$N_3$ and synthesised Kdo-$N_3$ [7,8]. The synthesised Kdo-$N_3$ was utilised for all super-resolution fluorescence microscopy experiments (dSTORM and 3D SIM[2]). Freeze-dried stocks of the synthesised compound were resuspended to a concentration of 400 mM in sterile, deionised water and stored at -20 °C until required.

**LPS metabolic labelling with Kdo-$N_3$ or Kdo-alkyne (Kdo-analogues)**

Pre-cultures were used to inoculate fresh supplemented M9 CDM charged with 4 mM of the Kdo-analogue (Kdo-$N_3$ or Kdo-alkyne) to a starting $OD_{600}$ of 0.05. Post-inoculation these metabolic labelling cultures were incubated for 12 – 14 h. Cultures were subsequently harvested upon reaching late stationary phase. Cells were isolated via centrifugation (8,000 x *g*, 3 min, 4 °C) and washed three times with fresh volumes of supplemented M9 CDM to remove non-specifically bound Kdo-analogue from cell surfaces prior to fluorescent labelling of the LPS.

***In situ* fluorescent labelling of Kdo-analogue-containing LPS inner core domains via Cu(I) catalysed azide-alkyne cycloaddition (CuAAC)**

Cells with Kdo-analogue-containing LPS in the OM were pelleted via centrifugation (10,000 x *g*, 4°C, 3 min) and resuspended to an $OD_{600}$ of 1.0 in 'Click-iT' reaction mix prepared according to the 'Click-iT' kit protocol ('Click-iT' Cell Reaction Buffer Kit, Molecular Probes®, Invitrogen) supplemented with 4 mM *N*-acetylneuraminic acid (Neu5Ac) and charged with the alkyne- (*for Kdo-$N_3$-containing LPS*) or azide- (*for Kdo-alkyne-containing LPS*) functionalised fluorescent dye ***(concentrations are specified in Extended Data Table S4)***. The cell suspensions were transferred to sterile 2 mL microcentrifuge tubes and incubated on a rotary wheel (12 rpm, 80° incline relative to bench top) for 30 min at room temperature protected from light. Post-labelling suspensions was transferred to fresh sterile 2 mL microcentrifuge tubes and cells bearing fluorescent LPS pelleted via centrifugation (10,000 x *g*, 4 °C, 3 min). The labelled cell pellets were washed a further three times with supplemented M9 CDM to ensure removal of residual fluorophore and 'Click-iT' mix components.

**Live cell Kdo-$N_3$-containing LPS *in situ* fluorescent labelling via Cu(I)-free strain promoted azide-alkyne cycloaddition (SPAAC)**

Washed cell samples with Kdo-$N_3$-labelled LPS were diluted in supplemented M9 CDM to an $OD_{600}$ of 1.0 and transferred to a sterile 2 mL microcentrifuge tube. The appropriate volume of a dibenzocyclooctyne-amine (DBCO)-functionalised AF488 dye was then added directly to the cell suspensions ***(Extended Data Table S5)***. The SPAAC labelling reactions were then incubated for 45 min at 30 °C on a rotary wheel (12 rpm, 80° incline relative to benchtop) protected from light. The cell suspensions were then each transferred to a fresh, sterile 2 mL microcentrifuge tube and pelleted via centrifugation (8,000 x *g*, 2 min, 4 °C). The labelled cell pellets were then gently washed three times in supplemented M9 CDM. Extra care was taken during resuspension of cell pellets to ensure cells were exposed to minimal levels of shear stress thereby ensuring minimal loss of cell viability.

**LptD fluorescent labelling via ncAA incorporation followed by CuAAC**

Sites for ncAA incorporation into LptD in *E. coli* BW25113 were selected based on *in silico* modelling of *E. coli* LptD crystal structure deposited in the PDB database [PDB: 4RHB] ***(Extended Data Fig. S1B)***. Four sites for propargyl-L-Lysine ncAA incorporation were selected based on criteria designed to minimise the impact of ncAA incorporation on LptD function and structure, whilst maximising the probability of ncAA incorporation at an accessible site to enable efficient downstream bio-orthogonal fluorophore coupling and minimise dynamic quenching of the coupled extrinsic probe by certain amino acids (*e.g*. tryptophan, histidine, methionine and tyrosine) [9]. We selected two lysine (K473 and K602) and two aspartic acid (D592 and D600) acid residues located on two unstructured, exposed extracellular loops of LptD. These selections were informed by results from published structural, computational and biochemical studies of LptD [10-12]. By avoiding the extracellular loops, residues and regions implicated in LptD function by these previous studies, we were able to minimise the probability that ncAA incorporation would have a deleterious impact on LptD structure and function.

Introduction of propargyl-L-Lysine ncAA to the culture medium resulted in leaky expression of the recombinant, amber-stop codon-containing *lptD* gene (*lptD**), *pylRS* and $tRNA^{Pyl}$ and subsequent production of recombinant, full length LptD with the ncAA integrated at one of four positions in its extracellular surface loops (LptD*). The accessible alkyne groups in the surface loops of LptD* enabled conjugation of an azide-functionalised, extrinsic fluorescent organic dye molecule via CuAAC [13,14]. LptD* fluorescent labelling enabled *in vitro* characterisation of the recombinant protein by fluorescence imaging of LptD-containing protein bands after SDS-PAGE ***(Extended Data Fig. S1C)***.

Direct LptD* fluorophore conjugation via CuAAC allowed us to carry out experiments without the use of immunoblotting for LptD and LptDE visualisation. Validation of ncAA-LptD production and subsequent fluorescent labelling was done for all four ncAA-containing *lptD* variants along with SDS-PAGE screens of recombinant protein production levels and ability to form a complex with LptE. Based on this screening process, the $LptD^{D600}$ mutant was selected and used in all dual-labelling experiments for LptD* and newly inserted LPS followed by visualisation via two-colour dSTORM super resolution microscopy ***(Fig. 1 and Extended Data Fig. S1)***.

**OmpA fluorescent labelling via ncAA incorporation followed by CuAAC**

To investigate the spatial distribution and organisation of LPS and OMPs relative to one another in the outer leaflet of the OM, OmpA was selected for labelling via adaptation of the strategy used for LptD having demonstrated in a previous study that OmpA and LPS are organised in discrete clusters in the OM of *E. coli* BW25113 cells[15]. OmpA is one of the most abundant OMPs in *E. coli* with approximately 100,000 molecules per cell [16,17], and lateral confinement of OmpA in the OM does not depend on its peptidoglycan-binding domain [18]. A three-dimensional model of the *E. coli* OmpA structure was produced using AlphaFold2 [19]. We then identified regions of the OmpA primary sequence predicted to form unstructured extracellular loops. We selected an accessible glutamic acid (E89) residue in extracellular loop 2 for propargyl-L-Lysine incorporation via mutation of the respective codon in the *ompA* gene sequence to the amber stop codon (TAG) [20-22].

After isolation and amplification of the *E. coli* BW25113 *ompA* gene sequence from the purified bacterial chromosomal DNA via PCR ***(Extended Data Table S2)***, we incorporated the wild-type *ompA* gene into a pBADcLIC expression vector. The target codon in the *ompA* gene was mutated to TAG via PCR mutagenesis ***(Extended Data Table S2)***. Under standard laboratory culture conditions, OmpA is a non-essential OMP. Therefore, we were able to co-transform the TAG-containing *ompA* expression vector into an *E. coli* Δ*ompA* strain ***(Extended Data Table S1)*** along with the aaRS / suppressor tRNA vector pEVOL-pylRS [2]. Full length ncAA-containing OmpA (OmpA*) production via leaky expression was initiated by charging cell cultures with 1 mM *N*-propargyl-L-Lysine-OH. OmpA* fluorescent labelling with a functionalised small extrinsic organic dye molecule was carried out *in situ* via CuAAC.

**Dual differential fluorescent labelling of LptD* and newly inserted Kdo-$N_3$-containing LPS**

*E. coli* BW25113 cells were co-transformed with pBADcLIC-lptD* and pEVOL-pylRS via electroporation as detailed above. A 5 mL volume of supplemented M9 CDM with 100 µg $mL^{-1}$ ampicillin and 35 µg $mL^{-1}$ chloramphenicol was inoculated with a single colony of freshly co-transformed cells and incubated at 37 °C with shaking (220 rpm) for approximately 7 h (to an $OD_{600}$ of 1.5 – 2.0). This pre-culture was used to inoculate a fresh volume of supplemented M9 CDM plus selection antibiotics to a starting $OD_{600}$ of 0.05. The culture was incubated as before for approximately 3 h, until it reached an $OD_{600}$ of 0.5. Uninduced recombinant LptD* production and LPS Kdo-$N_3$ metabolic labelling were initiated simultaneously via the addition of propargyl-L-Lysine and Kdo-$N_3$ to final concentrations of 1.0 mM and 4.0 mM, respectively. The recombinant LptD* production / LPS metabolic labelling culture was then re-incubated under standard conditions for a further 30 min. The culture was then removed and placed immediately on ice to halt further cell growth, division, new LPS insertion and LptD* production. The culture was diluted with pre-chilled, supplemented M9 CDM plus selection antibiotics to an $OD_{600}$ of 1.0 and transferred to a sterile pre-chilled 2.0 mL microcentrifuge tube on ice.

Cells were then harvested via centrifugation (10,000 x *g*, 3 min, 4°C) and the supernatant was removed. The cell pellet was then resuspended to an $OD_{600}$ of 1.0 in fresh, pre-chilled supplemented M9 CDM plus selection antibiotics via repeated, gentle pipetting. The resuspended culture was transferred to a sterile pre-chilled 2.0 mL microcentrifuge tube on ice, and the cells were re-pelleted via centrifugation. This washing step was repeated three times in total to ensure removal of residual propargyl-L-Lysine and Kdo-$N_3$. The washed cell pellet was then resuspended to an $OD_{600}$ of 1.0 in 'Click-iT' mix charged with either AF488- or AF647-alkyne, transferred to a 2 mL microcentrifuge tube, and incubated at room temperature for 30 min on a rotary wheel (12 rpm, 80° incline relative to benchtop) protected from light. In replicate experiments, the newly inserted LPS and LptD$^*$ labelling order was alternated, as were the dye combinations ***(Extended Data Table S6)***. This labelling strategy was implemented to confirm that the order in which species were labelled and the dye combinations used did not influence the apparent spatial distributions and relative concentrations of LptD* and newly inserted LPS ***(Fig. 1B and Extended Data Figure S1A)***. Cells were pelleted via centrifugation (10,000 x *g*, 3 min, 4°C), and the spent 'Click-iT' mix supernatant was removed. The cell pellet was then resuspended to an $OD_{600}$ of 1.0 in 0.2 µm-filtered, pre-chilled phosphate buffered saline (PBS, 137 mM NaCl, 2.7 mM KCl, 4.3 mM $Na_2HPO_4$, Sigma-Aldrich) pH 7.4 via gentle repeated pipetting.

**Preparation of labelled cell samples for direct stochastic optical reconstruction microscopy (dSTORM)**

Immediately after the 'Click-iT' labelling step, the cell suspension was transferred to a fresh pre-chilled 2 mL microcentrifuge tube and pelleted via centrifugation (10,000 x *g*, 2 min, 4 °C). Labelled cell pellets were re-suspended to an $OD_{600}$ of 1.0 in 0.2 µm-filtered PBS pH 7.4 supplemented with 2 mM $MgSO_4$ and 0.5 mM $CaCl_2$, and transferred to a sterile, pre-chilled 2 mL microcentrifuge tube. Cells were re-pelleted via centrifugation. This washing step was carried out a total of three times. The washed and labelled cell pellet was fixed via resuspension to an $OD_{600}$ of 1.5 in 4% (v/v) ultrapure, methanol-free paraformaldehyde (PFA, Polysciences Inc.) in PBS pH 7.4. This solution was incubated for 30 min on a rotary wheel (12 rpm, 80° incline relative to benchtop) at room temperature protected from light. The fixed cell suspension was then transferred to a sterile1.5 mL microcentrifuge tube and pelleted via centrifugation. The pellet of fixed and labelled cells was resuspended to an $OD_{600}$ of 1.0 in PBS pH 7.4 with 0.5 mM $MgSO_4$ and 0.15 mM $CaCl_2$, transferred to a fresh microcentrifuge tube and pelleted via centrifugation. This washing step was carried out three times in total.

**Mounting of fixed fluorescently labelled cell samples using poly-D-Lysine coated coverslips for imaging by dSTORM**

The washed, fixed, and labelled cell pellet in a microcentrifuge tube was placed on ice to cool to 4 °C and resuspended in degassed, pre-chilled glucose oxidase (GluOx) (Sigma Aldrich) / catalase (Cat) (Sigma Aldrich) $O_2$ scavenging buffer to an $OD_{600}$ of 1.5 via repeated gentle pipetting ***(Extended Data Table S7)*** [23,24]. 45.5 µL of the suspension was transferred to a pre-chilled, sterile 500 µL microcentrifuge tube on ice. 2.5 µL of 1 M β-mercaptoethylamine hydrochloride (β-MEA) (50 mM final concentration), 2 µL of 0.5% (w/v) 5 µm diameter silica bead slurry (final bead concentration = 0.02% (w/v)) and 0.5 µL TetraSpeck™ 0.2 µm microsphere standard solution (stock concentration ~ 1.5 x $10^9$ particles $mL^{-1}$, final concentration ~ 1.5 x $10^7$ particles $mL^{-1}$,Invitrogen Ltd.) were added. TetraSpeck™ beads were added to enable accurate AF488 and AZ647 channel alignment during two-colour dSTORM image processing. 10 µL of the suspension was then pipetted onto the centre of a pre-chilled (to minimise $O_2$ scavenging buffer enzyme activity), clean 1.0 – 1.2 mm thick glass slide. The slide was covered with a poly-D-lysine-coated, 18 mm x 18 mm high precision (Zeiss), no. 1.5 glass coverslip ensuring no bubbles were present in the slide chamber. The chamber was sealed with clear nail varnish. The slide was then incubated at 4 °C for 20 min while protected from light to allow the nail varnish to dry and the cells to adhere to the poly-D-lysine-coated coverslip surface.

**Two-colour dSTORM data acquisition with AF488- and AF647-labelled samples**

dSTORM imaging was done on a Zeiss Elyra 7 super-resolution imaging platform in laser widefield beam path mode using ZEN Black 3.0 SR FP2 software (Biosciences Technology Facility, University of York). Experiments were carried out using a Plan-Apochromat 63 x / 1.46 NA Korr oil immersion objective Var 2 lens together with TIRF uHP (ultra-high power) laser power density setting. Image areas were set to 128 pixels x 128 pixels for two cells or 64 pixels x 64 pixels for a single cell and saved in 16-bit format. Initial excitation and intersystem crossing of all fluorophores in the target cell(s) from their ground singlet state to an excited triplet 'dark' state was achieved via application of a short (200 – 300 frames, 50 ms exposure time), high intensity laser pulse (488 nm: 20-25% laser power, 642 nm: 10-15% laser power) with the microscope in epifluorescence (EPI) mode. dSTORM time series image sequences were then collected using highly inclined and laminated optical sheet (HILO) illumination for 10,000 frames (50 ms exposure time) using reduced laser powers (**488 nm:** Starting at 1% increasing to 10% over the time series, **642 nm:** Starting at 0.7% and increasing to 5% over the time series). To propagate fluorophore 'blinking' in the latter 5000 frames, 405 nm light was introduced during the 'transfer' phase of individual image collection to reduce fluorophore photobleaching resulting from simultaneous exposure to both 488 nm / 642 nm and 405 nm light [24]. HILO illumination mode was used to enable excitation of fluorophores in the outer membrane furthest from the coverslip thereby maximising the signal to noise (S:N) ratio. Additional dSTORM image acquisition settings are summarised in ***Extended Data Table S8***.

Two-colour dSTORM imaging of AF488- and AF647-labelled species in the OM of in-frame, in-focus cells were done sequentially using the Zeiss Elyra 7 super-resolution imaging platform with 488 nm and 642 nm excitation lasers. The BP420-480 + BP490-550 emission filter ***(Extended Data Table S8)*** was used to prevent cross channel 'bleeding' of fluorescence emission from the two dyes. dSTORM data for AF647-labelled species was collected first using the second sCMOS camera due to the lower photostability of the dye compared to AF488. dSTORM data for AF488-labelled species was then collected using the first sCMOS camera. After the initial high intensity burst to trigger fluorophore intersystem crossing from ground singlet states to a 'dark' triplet state, the fluorescent emission 'blinking' events were collected over 10,000 frames using a 50 ms exposure for each channel.

### Processing of dSTORM data

dSTORM raw data were processed using the SMLM (single-molecule localisation microscopy) processing facility found within the ZEN Black 3.0 SR software. Time series image sequences were converted to a crude single dSTORM image discarding overlapping molecules using a $x,y$ Gauss fit model, with a peak mask size of 9 pixels and a peak intensity to noise ratio of 7.0. Typical filtering settings are detailed in ***Extended Data Table S9***. SMLM-grouping settings were adjusted to minimise double counting of individual dye molecules (5 frames for the maximum "ON" time, 50 frames for maximum 'OFF' gap, 1.7 pixel for capture radius). Pixel resolution was set at 10 nm pixel$^{-1}$ with localised peaks displayed in gauss mode reflecting the degree of emission localisation precision. dSTORM image data was converted into images in .CZI format using the ZEN 3.0 SR software. Processed dSTORM data for individual channels was exported in .csv table formats for further processing and analysis. Images were exported to FIJI / ImageJ for final image production. Representative final images, processed in FIJI / ImageJ were saved in both .TIFF and .PNG formats.

### Spatial assessment of LptD* / newly inserted LPS via cross-correlation analysis of two-colour dSTORM data outputs

To quantify the spatial relationship between newly inserted LPS and the essential outer membrane protein LptD, two-dimensional spatial cross-correlation analysis was performed on two-colour dSTORM localisation data (exported from ZEN Black 3.0 SR FP2 software in .csv table format) from the OM of individual *E. coli* cells using MATLAB R2024b (64-bit) software. Localisation coordinates for labelled LptD* and corresponding newly inserted LPS were first realigned using principal component analysis (PCA) to correct for rotational variance and align the long axis of the cell with the *x*-dimension, enabling geometrically consistent estimation of cell length and width. Cell area was approximated by multiplying these dimensions.

To extract biologically meaningful radial correlation profiles, symmetric square grid of pairwise spatial displacements between all LptD* *(reference)* and newly inserted LPS *(target)* localisations were constructed. Each pairwise distance vector was binned radially in increments defined by the localisation precision. The bin width was set to 1.5× the median localisation precision across both channels (rounded to the nearest 10 nm), to ensure biologically resolvable separation of localisations into discrete shells without over-fragmentation of signal. The maximum radial displacement range was defined as 50% of the larger cell dimension (typically the long axis), capturing pairwise interactions across the full cell body while avoiding spurious correlations driven by distant points beyond the OM envelope. A null reference distribution was computed by randomising the LPS coordinates within the same cell envelope boundary, maintaining point density and cell geometry while eliminating true co-localisation. The final g($r$) values represent the ratio of observed LptD*– newly inserted LPS pairwise counts to those expected under this null model, at each radial distance.

Radial cross-correlation values were exported as g(*r*) profiles ***(Fig. 1C, Extended Data Fig. S1D and S1E)*** and further summarised using biologically interpretable parameters, including r_half-max (the radial distance at which the correlation drops to half-maximal) and half-max (the corresponding g(*r*) value), as measures of the strength and spatial extent of association between LptD and newly inserted LPS. All pairwise data and g(*r*) outputs were exported per cell for population-level summary construction and figure generation using OriginPro 2024b (64-bit) software. Example two-colour dSTORM input files (LPS_647.csv and LptD_488.csv; LPS_488.csv and LptD_647.csv), MATLAB script (LptD_new_LPS_crosscorrelation.m), associated Read.me and output files (Individual_cross_correlation_data_sets.csv, ***Extended Data Table S10)*** file are provided as Supplementary Software and Supplementary Data, respectively.

**LptD* and newly inserted LPS demograph generator from two-colour dSTORM coordinates**

To visualise the spatial distribution of LptD and newly inserted LPS across individual *E. coli* cells (n = 30), demographs were generated from two-colour dSTORM localisation data using MATLAB R2024b (64-bit) software ***(Fig. 1D)***. Localisation co-ordinates for LptD* and newly inserted LPS channels exported from processed two-colour dSTORM data in .csv table formats were grouped by 'Cell_ID' (1-30) and rotated using PCA such that the long axis of each cell was aligned with the *x*-axis. This geometric standardisation ensured consistent longitudinal orientation. Coordinates were then binned along the x-axis using a fine bin width of 25 nm, chosen to balance spatial resolution with noise reduction based on typical bacterial cell length scales and localisation precision (10–40 nm). Each binned intensity profile was normalised to its own maximum, enabling relative comparison of spatial enrichment patterns between cells of varying brightness or size. Demographs were centre-aligned by aligning the mid-point of the localisation distribution for each cell to a common reference. Missing values (NaNs) from uneven cell lengths or edge effects were rendered black to distinguish absent data from low signal. This approach was used to highlight population-wide trends while preserving per-cell spatial heterogeneity. The input data (LptD-newly-inserted_LPS_demograph_coordinates.csv), MATLAB script (LptD_LPS_Demograph_Generator.m) and associated Read.me file are provided as Supplementary Software and Supplementary Data, respectively.

#### Pearson correlation coefficient calculation from processed two-colour LptD* / newly inserted LPD dSTORM data

To quantify spatial correlation between LptD* and newly inserted LPS in the OM of individual *E. coli* cells, photon-weighted Pearson correlation coefficients (PCCs) were calculated. Two-colour dSTORM localisations were segmented by cell and assigned to either the LptD* or newly inserted LPS channel, according to the channel in which they were observed. Coordinates were mean-centred and aligned along the principal cell axis using PCA. Localisations were binned along the longitudinal axis in 25 nm intervals, and photon counts per bin were used to construct channel-specific intensity profiles. These were smoothed using a Gaussian kernel (σ = 1.5 bins) to reduce noise. Each cell was divided into three equal-length regions (two poles and mid-cell), and PCCs were calculated for each region and across the full cell length using MATLAB 2024b (64-bit). This approach enabled region-specific quantification of LptD–LPS co-localisation dynamics ***(Fig. 1E)***. The input data (demograph_combined_spreadsheet_photons.csv), MATLAB script (calculate_photon_weighted_PCCs.m), associated Read.me file and output data ***(Extended Data Table S11,*** PhotonWeighted_PCC_Output.csv) are provided as Supplementary Software and Supplementary Data, respectively.

#### Monitoring changes in background and newly inserted LPS distributions over time in the OM of *E. coli* BW25113 cells via Kdo-alkyne / Kdo-N3 pulse-chase experiments followed by two-colour dSTORM characterisation

A 5 mL volume of supplemented M9 CDM was inoculated with a single colony of *E. coli* BW25113 cells from a freshly streaked LB / Agar plate. Post-inoculation this pre-culture was incubated at 37 °C with shaking (220 rpm) for approximately 7 – 8 hours. This pre-culture was used to inoculate a separate volume of supplemented M9 CDM charged with 4.0 mM Kdo-alkyne to a starting $OD_{600}$ of 0.05. This background LPS metabolic labelling culture was then incubated for 14 h at 37 °C with shaking (220 rpm).

The following day the culture containing cells with background, Kdo-alkyne-containing LPS in the OM was transferred to ice to halt growth and further insertion of LPS. The culture was diluted to an $OD_{600}$ of 1.0 via addition of pre-chilled supplemented M9 CDM and transferred to a sterile, pre-chilled 2.0 mL microcentrifuge tube. Extra care was taken when handling and manipulating the culture to ensure minimal loss of viability resulting from processing steps. Cells were pelleted via centrifugation (8,000 x *g*, 3 min, 4 °C). After careful supernatant aspiration the pellet was resuspended to an $OD_{600}$ of 1.0 in pre-chilled supplemented M9 CDM via gentle pipetting and intermittent tube swirling (thereby minimising cell viability loss resulting from excessive shear stress). The suspension was transferred to a sterile, pre-chilled 2.0 mL microcentrifuge tube and cells re-pelleted via centrifugation. This washing step was repeated three times in total to ensure complete removal of residual Kdo-alkyne.

Cells were then resuspended in pre-warmed (to 37 °C) supplemented M9 CDM to an $OD_{600}$ of 0.5 and transferred to a sterile 50 mL screw-cap polypropylene tube. A ‘0 min’ sample was removed immediately upon resuspension and transferred to a pre-chilled, sterile 1.5 mL microcentrifuge tube on ice. Initiation of newly inserted LPS Kdo-$N_3$ metabolic labelling was then initiated via Kdo-$N_3$ addition to the remaining culture to a final concentration of 4.0 mM. Following this addition the culture was incubated as before and samples removed at 15 min intervals over the 2 h monitoring window. Cells were pelleted via centrifugation (10,000 x *g*, 3 min, 4 °C). Pellets were washed three times with supplemented M9 CDM. Washed pellets from each time point sample containing cells with Kdo-alkyne-labelled, background LPS and Kdo-$N_3$ -labelled, newly inserted LPS in the OM were resuspended to an $OD_{600}$ of 1.0 in ‘Click-IT’ mix charged with AF488- or AF647-alkyne / AF488- or AF647-azide, transferred to a 2 mL microcentrifuge tube, and incubated at room temperature for 30 min on a rotary wheel (12 rpm, 80° incline relative to benchtop) protected from light. In replicate experiments, the newly inserted LPS and pre-existing, background LPS labelling order was alternated, as were the dye combinations ***(Extended Data Table S12)***. This alternative labelling strategy was implemented in the experimental replicates to ensure that the order in which species were labelled and dye combination used did not influence the spatial distributions and relative concentrations of background and newly inserted LPS ***(Figs. 2B and 2C)***.

Cells were pelleted via centrifugation (10,000 x g, 3 min, 4°C) and the spent ‘Click-iT’ mix supernatant was carefully removed. The cell pellet was then resuspended to an $OD_{600}$ of 1.0 in 0.2 µm-filtered, pre-chilled PBS pH 7.4 via gentle repeated pipetting. The suspension was transferred to a fresh 2.0 mL microcentrifuge tube and cells were re-pelleted via centrifugation (10,000 x g, 3 min, 4 °C). The PBS supernatant was carefully removed, and the pellet resuspended to an $OD_{600}$ of 1.0 in ‘Click-iT’ mix charged with AF647- or AF488-azide / AF647- or AF488-alkyne. The suspension was transferred to a 2.0 mL microcentrifuge tube and incubated on a rotary wheel (12 rpm, 80° incline relative to bench top) at room temperature for 30 min protected from light. Cells were then pelleted via centrifugation (10,000 x g, 3 min, 4 °C). The spent ‘Click-iT’ mix was carefully removed, and the pellet resuspended in pre-chilled, filtered PBS pH 7.4 to an $OD_{600}$ of 1.0. The suspension was transferred to a sterile, pre-chilled 1.5 mL microcentrifuge tube and cells re-pelleted via centrifugation (10,000 x g, 3 min, 4 °C). This washing step was repeated three times in total. Cells were then fixed by re-suspension to an $OD_{600}$ of 1.0 in 4% (v/v) paraformaldehyde in PBS pH 7.4 followed by incubation at room temperature for 30 min on a rotary wheel (12 rpm, 80° incline relative to benchtop) protected from light. The suspension was then transferred to a sterile, pre-chilled 1.5 mL microcentrifuge tube and cells were pelleted via centrifugation (10,000 x g, 3 min, 4 °C). The pellet consisting of dual-labelled, fixed cells was washed three more times in pre-chilled, 0.2 µm-filtered PBS pH 7.4 as before. After the final wash step, the cell pellets were resuspended to an $OD_{600}$ of 1.0 in PBS pH 7.4 and stored at 4 °C or on ice, protected from light until imaging was completed.

### Two-colour dSTORM background and newly inserted LPS cluster overlap analysis in DBSCAN

Two-colour dSTORM localisation tables (background LPS and newly inserted LPS) were exported from ZEN Black v3.3 (Carl Zeiss Microscopy) as .csv files and analysed in Python 3.10.12 (NumPy 1.26.0, pandas 2.2.1, shapely 2.0.3, scikit-learn 1.5.0). Both newly inserted and background LPS localisations for each individual *E. coli* cell were merged, and a convex hull was expanded by 10 nm to define the OM outline. The polygon was simplified (2 nm tolerance) to remove sub-pixel kinks. PCA rotated each cell, so its long axis lay horizontally, ensuring a common reference frame throughout the 120-min pulse–chase.

Spatial domains of background, pre-existing- and newly inserted LPS were identified independently using a tailored Density-Based Spatial Clustering of Applications with Noise (DBSCAN) algorithm ($\varepsilon = 30$ nm, min_samples = 7) [25], widely adopted for co-localisation analysis of multi-channel SMLM data [26]. The $\varepsilon$ value equals $1.5 \times$ the median nearest-neighbour distance between LPS localisations under our imaging conditions (median = 22 nm) and reflects the average precision associated with individual localisations (~30 nm), balancing sensitivity to genuine nanoscale domains with robustness to photon shot noise. A min_samples threshold of seven exceeds the upper-quartile localisation count for single fluorophore bursts, limiting false-positive clusters. Cluster polygons were clipped to the OM boundary and combined to obtain total old- and new-LPS cluster areas; their geometric intersection (background LPS $\cap$ newly inserted LPS) defined the overlap region. The script produces several output files. LPS_output_summary.csv contains numerical values for the OM area, individual and combined LPS areas, localisation counts inside the overlap, and normalised coverage metrics; these underpin the quantitative time-course displayed in ***Figs. 2E and 2F*** and are reproduced in ***Extended Data Table S13***. combined_overlap_map.png is a high-resolution composite image showing all localisation points, their DBSCAN-assigned clusters and the spatial overlap; the map provides morphological context for the representative cells provided in ***Extended Data Fig. S2E***. Cluster-size distributions are exported as old_lps_cluster_histogram.csv and new_lps_cluster_histogram.csv, enabling independent verification that old and new clusters are sampled from comparable size ranges. Examples of individual channel input dSTORM data for each time point (DBSCAN_new_old_LPS_dSTORM_input_folder), Python script (Python_code_New_Old_LPS_DBSCAN.txt) and associated READ.ME_Python_code_New_Old_LPS_DBSCAN file, example individual cell OM output data files (DBSCAN_new_old_LPS_outputs_folder) along with summary data output tables (DBSCAN_new_old_LPS_individual_cell_summary.xlsx and ***Extended Data Table S13***) are provided as Supplementary Software and Supplementary Data, respectively.

**LPS extraction**

LPS was extracted via adaption of the hot phenol – aqueous method developed by Westphal (1965) [27]. The cell pellet was re-suspended to an $OD_{600}$ of 0.5 in PBS pH 7.4 plus 0.5 mM $MgCl_2$ and 0.15 mM $CaCl_2$ and pelleted via centrifugation (10,000 x *g*, 10 min, 4 °C). The washed cell pellets were then resuspended to the same $OD_{600}$ in ultrapure deionised water (resistivity = 18.2 MΩ cm) and transferred to a glass dram with Teflon-coated lid. An equal volume of 90% (w/v) phenol, pre-warmed to 65 °C was added and the emulsion stirred vigorously at 65 °C for 15 min. Drams were then transferred to ice for 15 min to cool. Emulsions were then transferred to microcentrifuge tubes and centrifuged (8,500 x *g*, 10 min, 15 °C). The LPS-containing aqueous fraction was transferred to a 50 mL screw-cap polypropylene tube. A second aqueous extraction was then carried out on the remaining phenol layer and the aqueous fractions were pooled in the 50 mL screw-cap polypropylene tube. Sodium acetate was added to the pooled aqueous fractions to a final concentration of 0.5 M to ensure LPS precipitation. 10 volumes of 95% (v/v) ethanol, pre-chilled to -20 °C was then added and after mixing via repeated inversions the suspension was incubated overnight to maximise LPS precipitation. The following day LPS was pelleted via centrifugation (2,000 x *g*, 10 min., 4 °C). After supernatant aspiration, the LPS pellet was dried under a steady stream of $N_{2(g)}$. The dried LPS pellet was resuspended in 100 µL ultrapure deionised water and transferred to a 1.5 mL microcentrifuge tube. Sodium acetate was added to a final concentration of 0.5 M and 1.1 mL pre-chilled 95% (v/v) ethanol was added. After mixing the tube contents via repeated inversion, the LPS was re-precipitated via overnight incubation at -20 °C. Precipitated LPS was pelleted via centrifugation. The pellet was dried again under $N_{2(g)}$ and finally resuspended in 50 µL PBS pH 7.4.

**LPS Tricine-Sodium Dodecyl Sulfate-Polyacrylamide Gel Electrophoresis (TSDS-PAGE)**

A volume (15 µL) of the purified LPS was mixed with a volume (5 µL) of 4x loading buffer ***(Extended Data Table S14)*** plus 2% (v/v) β-mercaptoethanol (β-ME). After vortexing and a brief centrifugation, the samples were incubated for 1 h at 40 °C. The samples were then re-vortexed and centrifuged briefly before loading the entire sample volume into the well of a 10 cm x 10 cm x 1 mm 15% (w/v) acrylamide resolving / 6% (w/v) acrylamide stacking The Tricine gel was prepared according to the protocol published by Schägger (2006) [28].

Commercial smooth LPS standards (Thermo Scientific®) were analysed alongside extracted LPS samples to facilitate classification of sample LPS bands. O55:B5-AF488 LPS conjugate samples were prepared for TSDS-PAGE by adding 1 µL of 1 mg $mL^{-1}$ LPS conjugate stock solution to 9 µL ultrapure deionised water followed by 3.3 µL of 4 x loading buffer plus 2% (v/v) β-ME ***(Extended Data Table S14).*** Samples were vortexed and briefly centrifuged before being incubated at 40 °C for 1 h, then re-vortexed and briefly centrifuged again before loading the entire sample volume into the well of a Tricine gel. Electrophoresis was carried out at 4 °C using pre-chilled buffers with the gel tank protected from light to prevent fluorophore (AF488) photobleaching. Electrophoresis was done in constant current mode for 45 min at 30 mA until a dye front formed as a fine horizontal line at the stacking / resolving gel interface. The current was then increased to 60 mA for approximately 3 h until the dye front was approximately 1 cm from the base of the gel.

**Processing and visualisation of LPS TSDS-PAGE gels**

Upon completion of electrophoresis, the gels were fixed via immersion in 150 mL 50% (v/v) methanol / 3% (v/v) glacial acetic acid and incubated overnight at room temperature on a slow-moving rocker, protected from light. The following day gels were washed via immersion in 150 mL 3% (v/v) glacial acetic acid and incubated for 20 min at room temperature on a slow-moving rocker. This step was repeated three times in total. AF488-LPS bands were then visualised using a Typhoon 5 bioimaging system (Amersham) equipped with a 488 nm argon laser and 525 BP filter. To visualise total LPS bands, gels were stained using a Pro-Q™ Emerald 300 LPS gel stain kit according to the manufacturer's instructions (Thermofisher Scientific®). Briefly, after fixation and washing steps, LPS carbohydrates were oxidised via immersion in 30 mL 1% (w/v) $H_5IO_6$ followed by incubation for 45 min on a slow-moving rocker at room temperature. Gels were washed three times in 150 mL 3% (v/v) glacial acetic acid for 20 min. Gels were then stained via immersion in 25 mL dilute (1:50) Pro-Q Emerald 300 stain and incubated for 90 min at room temperature on a slow-moving rocker. After post-staining washing steps, Pro-Q stained LPS bands were visualised under UV illumination using a GeneGenius gel imaging system.

**Quantifying relative rates of background LPS turnover on a cell-by-cell basis via Kdo-$N_3$ / native Kdo pulse-chase experiments**

A 5 mL volume of supplemented M9 CDM was inoculated with a single colony of the *E. coli* BW25113 strain picked from a freshly streaked LB / agar plate. Post-inoculation the culture was incubated at 37 °C with shaking (220 rpm) for approximately 7 hours. This pre-culture was used to inoculate a fresh volume of supplemented M9 CDM containing 4.0 mM Kdo-$N_3$ to an $OD_{600}$ of 0.05. This background LPS Kdo-$N_3$ metabolic labelling culture was then incubated for 12 hours at 37 °C with shaking (220 rpm). The culture was then transferred to ice and diluted with pre-chilled M9 CDM without glucose or casamino acids ***(Extended Data Table S3)*** to an $OD_{600}$ of 1.0 and cells were pelleted via centrifugation (8,000 x *g*, 3 min, 4 °C). After careful supernatant aspiration, the pellet was resuspended to an $OD_{600}$ of 1.0 in pre-chilled supplemented M9 CDM via gentle pipetting and intermittent tube swirling (thereby minimising cell viability loss resulting from excessive shear stress). The suspension was transferred to a sterile, pre-chilled 2.0 mL microcentrifuge tube and cells re-pelleted via centrifugation. This washing step was repeated three times in total to ensure complete removal of all residual Kdo-$N_3$.

The washed pellet of cells with Kdo-$N_3$-labelled, background LPS in the OM were resuspended to an $OD_{600}$ of 0.5 in pre-warmed supplemented M9 CDM charged with 4 mM native Kdo and incubated under standard conditions. Samples were taken at 30 min intervals over a 3.5 h period and $OD_{600}$ measurements were done at time of sampling in triplicate to ensure accuracy and reliability. Samples were transferred to pre-chilled 2.0 mL sterile microcentrifuge tubes on ice and diluted with pre-chilled M9 CDM without glucose and casamino acids to ensure cell growth, division and LPS insertion were arrested. Cells were then pelleted via centrifugation (10,000 x *g*, 3 min, 4 °C).

After careful supernatant aspiration, the pellets were resuspended to an $OD_{600}$ of 1.0 in pre-chilled M9 CDM without glucose and casamino acids on ice. Suspensions were transferred to fresh sterile 2.0 mL microcentrifuge tubes and cells re-pelleted via centrifugation. This washing step was repeated three times in total. Washed pellets of *E. coli* BW25113 cells containing background, pre-existing Kdo-azide-LPS in the OM were resuspended to an $OD_{600}$ of 1.0 in 'Click-IT' mix charged with AF488-alkyne. Suspensions were transferred to sterile 2 mL microcentrifuge tubes and incubated at room temperature on a slow-moving rotary wheel for 30 min (12 rpm, 80 ° incline relative to benchtop) protected from light. Following this labelling step, the suspensions were transferred to sterile 1.5 mL microcentrifuge tubes and cells pelleted via centrifugation (10,000 x *g*, 3 min, 4 °C). After aspiration of the spent 'Click-iT' mix supernatants labelled cell pellets were resuspended to an $OD_{600}$ of 1.0 in pre-chilled, 0.2 µm-filtered PBS pH 7.4. Suspensions were transferred to fresh, sterile 1.5 mL microcentrifuge tubes and the cells were re-pelleted via centrifugation. This washing step was repeated three times in total. Cells were then fixed via re-suspension to an $OD_{600}$ of 1.0 in 4% (v/v) paraformaldehyde in PBS pH 7.4 followed by incubation at room temperature for 30 min on a rotary wheel (12 rpm, 80° incline relative to benchtop) protected from light. The suspension was then transferred to a fresh, pre-chilled 1.5 mL microcentrifuge tube and cells were pelleted via centrifugation (10,000 x g, 3 min, 4 °C). The pellet consisting of dual-labelled, fixed cells was washed three more times in pre-chilled, 0.2 µm-filtered PBS pH 7.4 as before. After the final wash step, the cell pellets were resuspended to an $OD_{600}$ of 1.0 in PBS and stored at 4 °C or on ice, protected from light until imaging was completed.

**Quantifying relative rates of newly inserted LPS insertion on a cell-by-cell basis via native Kdo / Kdo-$N_3$ pulse-chase experiments**

A 5 mL volume of supplemented M9 CDM was inoculated with a single colony of *E. coli* BW25113 cells picked from a freshly streaked LB / agar plate. Post-inoculation the culture was incubated at 37 °C with shaking (220 rpm) for approximately 7 hours. This pre-culture was used to inoculate a fresh volume of supplemented M9 CDM containing 4.0 mM native Kdo to an $OD_{600}$ of 0.05. This culture was then incubated with shaking (220 rpm) for 12 hours. The culture was then transferred to ice and dilute with pre-chilled supplemented M9 CDM without glucose or casamino acids to an $OD_{600}$ of 1.0 and cells were pelleted via centrifugation (8,000 x *g*, 3 min, 4 °C). After careful supernatant aspiration, the pellet was resuspended to an $OD_{600}$ of 1.0 in pre-chilled supplemented M9 CDM via gentle pipetting and intermittent tube swirling (thereby minimising cell viability loss resulting from excessive shear stress). The suspension was transferred to a sterile, pre-chilled 2.0 mL microcentrifuge tube and cells re-pelleted via centrifugation. This washing step was repeated three times in total to ensure complete removal of all residual native Kdo. The washed pellet of cells was resuspended to an $OD_{600}$ of 0.5 in pre-warmed supplemented M9 CDM charged with 4 mM Kdo-$N_3$ and incubated under standard conditions.

Samples were taken at 30 min intervals over a 3.5 h period and $OD_{600}$ measurements were done at the time of sampling in triplicate to ensure accuracy and reliability. After removal, the samples were transferred to sterile pre-chilled 2.0 mL microcentrifuge tubes on ice and diluted with pre-chilled M9 CDM without glucose and casamino acids to ensure cell growth, division and LPS insertion were arrested. Cells were then pelleted via centrifugation (10,000 x *g*, 3 min, 4 °C). After careful supernatant aspiration, the pellets were resuspended to an $OD_{600}$ of 1.0 in pre-chilled M9 CDM without glucose and casamino acids. Suspensions were transferred to fresh sterile pre-chilled 2.0 mL microcentrifuge tubes and cells re-pelleted via centrifugation. This washing step was repeated three times in total.

Washed pellets of *E. coli* BW25113 cells containing newly inserted Kdo-$N_3$-labelled LPS in the OM were resuspended to an $OD_{600}$ of 1.0 in 'Click-IT' mix charged with AF488-alkyne. Suspensions were transferred to sterile 2 mL microcentrifuge tubes and incubated at room temperature on a slow-moving rotary wheel for 30 min (12 rpm, 80 ° incline relative to benchtop) protected from light. Following this labelling step, the suspensions were transferred to sterile 1.5 mL microcentrifuge tubes and cells pelleted via centrifugation (10,000 x *g*, 3 min, 4 °C). After aspiration of the spent 'Click-iT' mix supernatants, the labelled cell pellets were resuspended to an $OD_{600}$ of 1.0 in pre-chilled, 0.2 µm-filtered PBS pH 7.4. Suspensions were transferred to fresh, sterile 1.5 mL microcentrifuge tubes and the cells were re-pelleted via centrifugation. This washing step was repeated three times in total. Cells were then fixed via re-suspension to an $OD_{600}$ of 1.0 in 4% (v/v) paraformaldehyde in PBS pH 7.4 followed by incubation at room temperature for 30 min on a rotary wheel (12 rpm, 80° incline relative to benchtop) protected from light. The suspension was then transferred to a fresh, pre-chilled 1.5 mL microcentrifuge tube and cells were pelleted via centrifugation (10,000 x g, 3 min, 4 °C). The pellet consisting of dual-labelled, fixed cells was washed three more times in pre-chilled, 0.2 µm-filtered 1 x PBS pH 7.4 as before. After the final wash step, the cell pellets were resuspended to an $OD_{600}$ of 1.0 in PBS and stored at 4 °C or on ice, protected from light until imaging was completed.

**Single-colour dSTORM imaging**

dSTORM imaging was done on a Zeiss Elyra 7 microscope in the laser widefield beam path mode using ZEN Black 3.0 SR FP2 software (Biosciences Technology Facility, University of York). Experiments were carried out using a Plan-Apochromat 63 x / 1.46 NA Korr oil immersion objective Var 2 lens together with TIRF uHP (ultra-high power) laser power density setting. Image areas were set to 128 pixels x 128 pixels for two cells or 64 pixels x 64 pixels for a single cell and saved in 16-bit format. Initial excitation and intersystem crossing of all fluorophores in the target cell(s) from their ground singlet state to an excited triplet 'dark' state was achieved via application of a short (200 – 300 frames, 50 ms exposure time), high intensity 488 nm laser pulse (15% laser power) with the microscope in epifluorescence (EPI) mode. dSTORM time series image sequences were then collected using highly inclined and laminated optical sheet (HILO) illumination for 5,000 frames (50 ms exposure time) in total. For the first 1000 frames a laser power of 1.5% was applied, this was increased to 2.5% between 1000 – 2000 frames, 3.5% for 2000 – 3000 frames, 4.5% for 3000 – 4000 frames and finally to 5.5% laser power between 4000 and 5000 frames.

To propagate fluorophore ‘blinking’ in the latter 3000 frames, 405 nm light was introduced during the ‘transfer’ phase of individual image collection to reduce fluorophore photobleaching resulting from simultaneous exposure to both 488 nm and 405 nm light [24]. HILO illumination mode was used to enable excitation of fluorophores in the outer membrane furthest from the coverslip thereby maximising the signal to noise (S:N) ratio. Additional dSTORM image acquisition settings are summarised in ***Extended Data Table S8***.

**Single-colour dSTORM data processing and measurement of discrete AF488 emission abundance in the outer membrane of *E. coli* BW25113.**

To ensure the compatibility of data derived from different cells, all dSTORM images were processed using a standard protocol with identical filter settings. dSTORM raw data were processed using the SMLM processing facility found within the ZEN Black 3.0 SR software. Time series image sequences were converted to a crude single dSTORM image discarding overlapping molecules using a $x,y$ Gauss Fit model, with a peak mask size of 9 pixels and a peak intensity to noise ratio of 6.5. Standard filtering settings used for all images are detailed in ***Extended Data Table S15***. SMLM grouping settings were adjusted to minimise double counting of individual dye molecules (5 frames for the maximum “ON” time, 50 frames for maximum ‘OFF’ gap, 1.7 pixel for capture radius). Pixel resolution was set at 9 nm pixel$^{-1}$ with localised peaks displayed in gauss mode reflecting the degree of localisation precision. The number of filtered spots was recorded and used to calculate AF488 emission abundance per µm$^2$ on the OM surface after measurement of the OM surface area in ImageJ.

**Fluorescence recovery after photobleaching (FRAP) and confocal fluorescence microscopy**

Fluorescence confocal microscopy analyses and FRAP experiments were conducted using either an upright Zeiss LSM710 confocal fluorescence microscope or an inverted Zeiss LSM 780 multiphoton confocal fluorescence microscope (Biosciences Technology Facility, University of York). Confocal imaging and FRAP quantitative and statistical analyses of FRAP data were done according to protocols used in a previous study [15].

**DBSCAN analysis of single-colour dSTORM data using half-rod surface projection in LPS pulse–chase experiments**

To quantify spatial features of newly inserted and background LPS-rich regions in the OM of individual *E. coli*, we developed a MATLAB pipeline that performed DBSCAN-based clustering on single-colour dSTORM datasets from pulse-chase experiments projected onto a three-dimensional (3D) half-rod geometry. Raw localisations were first pre-processed to remove missing values and aligned along the long axis of each cell using PCA. Data was then filtered to match a canonical *E. coli* rod geometry comprising a central cylindrical body and hemispherical polar caps. To account for projection distortion due to curvature of the bacterial surface, localisations were radially mapped onto a 3D half-rod surface defined by the visible hemisphere and top half of the cylindrical outer membrane. This strategy improved the geometric accuracy in density calculations, particularly at the poles. Unlike commonly used 2D clustering approaches, this method embeds localisation data into a biologically realistic 3D surface model, enabling more accurate quantification of cluster density and position across the full cell envelope.

Clustering was performed using DBSCAN ($\varepsilon$ = 30 nm, minPts = 7), parameters chosen to match those used previously for two-colour overlap analyses, and to reflect the spatial resolution and labelling density typical of our dSTORM datasets. For each identified cluster, 2D convex hulls were computed in the aligned rod frame to calculate area and per-cluster localisation density (points per $nm^2$). A Z-height threshold was used to exclude clusters located near the edge of the projected rod surface ($Z < 20\%$ of cell radius), which are more susceptible to projection artefacts. Global cluster density was calculated as the total number of clusters divided by the estimated surface area of the half-rod model, accounting for both cylindrical and hemispherical components. The pipeline was designed to be both modular and robust, with adjustable geometric and clustering parameters. It was designed for reproducible batch processing of individual cell datasets. The outputs of the pipeline were used to quantify temporal trends in LPS insertion and loss at both whole-cell and subcellular levels over the experimental time course. MATLAB scripts (MATLAB_DBSCAN_Background_or_New_LPS_dSTORM_individual_cell_code, new_lps_or_old_LPS_cluster_summary_graphs_code.txt), associated Read.me files, summary output data files (DBSCAN_background_LPS_ _all_data.xlsx, DBSCAN_new_LPS_all_data.xlsx) along with summary data output tables ***(Extended Data Tables S18 and S19)*** are provided as Supplementary Software and Supplementary Data, respectively.

**Region-resolved visualisation of LPS clusters across OM of single cells**

To investigate whether cluster properties for newly inserted or background LPS varied between distinct spatial domains in the OM of *E. coli* cells, we implemented a MATLAB script to generate globally scaled, region-coloured cluster maps for individual E. coli cells. DBSCAN-derived cluster centroids identified in the previous analyses were classified as *mid-cell* or *polar* based on their *x*-coordinate relative to the cell midpoint (with poles defined as ±1.5× the cell radius). Cluster marker sizes were scaled globally by the square root of cluster area, enabling direct visual comparison across cells and timepoints. This visualisation allowed interrogation of spatial patterning in both background LPS and newly inserted LPS clusters following Kdo-$N_3$ / native Kdo pulse–chase labelling. By applying a consistent classification scheme across all timepoints, the script enabled robust per-cell assessment of whether cluster distribution or size varied systematically between mid-cell and polar regions, and whether this organisation changed over time. The resulting maps provided an essential complement to global metrics, clarifying that distinct regional distributions of LPS clusters were maintained over time despite OM turnover. The MATLAB script (MATLAB_DBSCAN_Background_New_LPS_clusters_by_region.txt) and associated READ.ME_MATLAB_DBSCAN_Background_New_LPS_clusters_by_region.txt file, are provided as Supplementary Software and Supplementary Data, respectively.

**Preparation of unsupplemented M9 pH 7.2 / ultrapure low melting point agarose pads**

The optimised protocol was based on a method published previously by Skinner et al, 2013 [29]. Slides were cleaned without exposure to any type of detergent. Briefly, slides were washed twice in acetone, washed twice in propan-2-ol and rinsed for 10 minutes under slow flowing deionised water. They were then placed in a drying oven, protected from dust overnight. The following day a 125 µL gene frame (Thermo Fisher Scientific) was placed flat at the centre of the slide and left overnight to set.

The following day the slide plus adhered gene frame was placed in a drying oven and heated to approximately 40 °C. 1 mL sterile 2% (w/v) ultrapure low melting point (LMP) agarose (Thermo Fisher Scientific) in unsupplemented M9 CDM was prepared in a 50 mL screw-cap polypropylene tube via repeated, short duration, low power heating in a microwave (5 x 10 – 15 s at 5% power). Once the agarose had dissolved the tube was transferred to an ultrasonic water bath pre-heated to 50 °C and incubated for ~ 5 minutes to degas. 150 µL of the degassed, molten 2% (w/v) agarose / M9 solution was pipetted into the gene frame cavity in a laminar flow hood and covered with a clean glass slide. A weight was then placed on the top slide and the agarose / M9 was left to set for 60 min. After 60 min, the weight was removed and the agarose / M9 pad sealed in Parafilm® and stored at 4 °C until required.

**Mounting of live *E. coli* BW25113 cells with fluorescently labelled background LPS for time-lapse 3D SIM$^2$ imaging**

Post Cu(I)-free SPAAC labelling, the cells were resuspended to an $OD_{600}$ of 2.0 in pre-warmed (to 37°C) supplemented M9 CDM and incubated for 5 ‑ 10 min at 37 ° C. The agarose / M9 pad was incubated at 37°C for 30 min after which a 1.5 cm x 1.5 cm slice was cut using an ethanol cleaned scalpel and placed flat at the centre of a cleaned, pre-warmed glass slide. 10 µL of the labelled cell suspension was pipetted onto the centre of the pad and left for one minute in a laminar flow hood for the cells to adsorb. An additional 5 µL pre-warmed, supplemented M9 was deposited around the edges of the pad.

A pre-warmed, cleaned (without detergent) glass, high-precision 18 mm x 18 mm no. 1.5 glass coverslip (Zeiss) was then gently placed over the pad. The slide was sealed with a molten VALAP (1: 1: 1 mixture of Vaseline, lanolin, and paraffin) bead to prevent agarose pad desiccation during time course imaging [30].

**Time-lapse 3D SIM$^2$ imaging of live *E. coli* BW25113 cells with fluorescently labelled background LPS**

Slides were mounted onto a Zeiss Elyra 7 microscope in lattice SIM mode with a fitted incubator that had been prewarmed to 37°C (for at least 12 h prior to the experiment to allow all components to equilibrate fully to the elevated temperature). Individual image acquisition settings are summarised in ***Extended Data Table S20***. *Z*-stack images were collected at 2- to 5-minute intervals using low laser powers (0.5 – 1.0%) to minimise AF488 dye photobleaching thus extending the duration of time-lapse experiments whilst care was taken not to compress agarose pads by over-extending *z*-stack depth. *z*-stacks were collected over the largest possible cross-sectional areas (2560 pixels x 2560 pixels / 80.14 µm x 80.14 µm) to maximise data collection since increasing the field of view did not affect *xy* resolution (0.03 µm pixel$^{-1}$). Individual experiments were terminated once the signal intensity : noise ratio for the AF488 fluorescence signal was deemed inadequate for reliable analysis, *i.e.* at the point where fluorescence signal in the OM region could not be reliably assigned to fluorescently labelled LPS rather than resulting from background ‘noise’. The duration of each experiment was usually 25 – 45 minutes depending on initial labelling efficiency and AF488-LPS signal strength).

**3D SIM² z-stack image processing and re-construction**

3D SIM z-stack images were processed using the SIM² program within ZEN Black 3.0 SR software using 'weak' default settings to minimise loss of detail during image processing. Final 2D depth coded images were produced in ZEN 3.1 (blue edition) in Carl Zeiss Image (.czi) (for further processing) and .PNG (for presentation) formats. 2D image cross-sections (lower cell surface, mid-cell and upper cell surface) were produced in FIJI / ImageJ (version 1.54h, with BioFormats plug-in installed) by summing the two slices from the 3D SIM² z-stacks that corresponded to each of these image cross-sections. 3D depth coded images were produced in Zen Blue 3.1 software, and were saved in both .TIFF and .PNG formats. As a control, to demonstrate standard rates of cell elongation and division, the lengths of randomly selected cells were measured at each time point and their normalised cell length (to the maximum observed length for each cell) plotted against time ***(Extended Data Fig. S5D)***.

**Biochemical assessment of background LPS turnover in intact *E. coli* BW25113 cells and its corresponding accumulation in culture supernatants due to OMV release using TSDS-PAGE**

Labelled intact cell samples were prepared via the same Kdo-$N_3$ / native pulse-chase protocol used to monitor background LPS turnover on a cell-by-cell basis using dSTORM analysis, but without the final paraformaldehyde fixation step. LPS was extracted from cell pellets using the protocol described earlier. The volume of LPS sample loaded was normalised based on $OD_{600}$ measurements ***(Extended Data Figs. S3C and S3D)*** done at the point of sampling by diluting the sample with the appropriate volume of HPLC water to ensure equal amounts of total LPS were loaded in each lane. 6.67 µL aliquot of 4 x loading buffer ***(Extended Data Table S14)*** with 3% (v/v) β-Mercaptoethanol was added to each 20 µL cell pellet-derived LPS sample in a 500 µL microcentrifuge tube. Samples were then mixed via repeated pipetting and gentle vortexing, and then briefly centrifuged. Suspensions were then incubated at 40 °C for 1 h in a heated water bath. After incubation, the samples were re-vortexed and briefly centrifuged before loading 20 µL of each sample per well in a Tricine gel.

**OMV isolation, background LPS fluorescent labelling and OMV-derived LPS extraction.**

After intact cell pellet and culture supernatant isolation via centrifugation (10,000 x g, 20 min, 4 °C) from Kdo-$N_3$ / native Kdo pulse-chase samples, the supernatants were transferred to sterile pre-chilled 2.0 mL microcentrifuge tubes and immediately re-centrifuged to remove any remaining residual debris (10,000 x g, 20 min, 4 °C). Double centrifuged supernatants were then carefully aspirated into new, sterile pre-chilled 2.0 mL microcentrifuge tubes and then passed slowly through 0.45 μm and 0.20 μm pore size hydrophilic (PES) syringe filters into pre-chilled, polycarbonate tubes. OMVs in the culture supernatant were then pelleted via ultracentrifugation (Beckman Coulter TLX Optima Benchtop Ultracentrifuge, TLA 100.4 fixed angle rotor) (110,000 x g, 12 hr, 4 °C). Supernatants were aspirated and OMV pellets resuspended in 200 μL pre-chilled 0.2 μm-filtered PBS pH 7.4 (with 2 mM $MgSO_4$ and 0.5 mM $CaCl_2$ added) and transferred to sterile 2 mL microcentrifuge tubes. Background Kdo-$N_3$-containing LPS in OMVs was then fluorescently labelled via CuAAC using the identical protocol used to label background LPS in the OM of intact cells, thereby ensuring no variations in labelling efficiency arose from differences in labelling protocols. OMV 'Click-iT' labelling suspensions in 2.0 mL microcentrifuge tubes were incubated on a slow-moving rotary wheel at room temperature for 30 min (12 rpm, 80 ° incline relative to benchtop) protected from light. Suspensions were then transferred to 3.5 kDa D-Tube Midi dialysers (Merck) and suspensions dialysed twice for 6 to 8 hours into 3 litres of unsupplemented M9 CDM (with 2 mM $MgSO_4$ and 0.5 mM $CaCl_2$ added) at 4 °C, protected from light to remove spent 'Click-iT' mix components. Dialysed suspensions were transferred to 250 μL pre-chilled polycarbonate ultracentrifugation tubes and OMVs bearing AF488-labelled background LPS were pelleted via ultracentrifugation (110,000 x *g*, 12 h, 4 °C). Supernatants were aspirated and OMV pellets dried for 5 – 10 min under a steady stream of nitrogen gas. OMV pellets were then resuspended in 30 μL LPS lysis buffer with 2% (v/v) β-ME ***(Extended Data Table S21)*** via repeated pipetting and transferred to 500 μL microcentrifuge tubes. Tubes were incubated in an ultrasonic water bath for 5 minutes at 40 °C to lyse OMVs. Suspensions were stored at 4 °C, protected from light prior to TSDS-PAGE analysis. OMV-derived LPS samples were prepared for TSDS-PAGE using the same method as for LPS extracted from bacterial cell pellets.

**Quantifying background AF488-labelled LPS in bacterial OM and OMV-derived fractions from TSDS-PAGE gel images by densitometry**

To verify AF488-LPS signal intensities were not influenced by unexpected differences in total LPS loading, control experiments were done using non-Kdo-azide labelled samples (since Kdo-azide incorporation into LPS affects Pro-Q LPS staining efficiency). The samples were loaded into the wells of a Tricine gel based on $OD_{600}$ measurements at the point of sampling and total LPS visualised via Pro-Q 300 staining (according to the protocol detailed earlier, ***Fig. 6D and Extended Data Fig. S6C).*** Densitometric analyses were done using ImageJ / FIJI software. The average raw AF488-LPS band intensities were measured and recorded. The average AF488-LPS band intensity (minus background signal) was normalised to enable the calculation of average relative changes in background AF488-LPS content across multiple experimental replicates and to facilitate comparison between cell pellet and OMV-derived LPS. Normalised values ranged from 1 at 0 h post-Kdo-$N_3$ removal to 0 at 3.5 h post-Kdo-$N_3$ removal. Normalised background AZ488-LPS values for individual replicates and mean AZ488-LPS values for cell pellets ***(Fig. 6D, Extended Data Fig. S6A)*** and OMV-derived samples ***(Fig. 6D, Extended Data Fig. S6B)*** were plotted against time post-Kdo-$N_3$ removal.

**Assessing Kdo-$N_3$ incorporation into the OM of exponential and stationary phase *E. coli* BW25113 cells**

A starter culture was prepared by inoculating 5 mL of supplemented M9 CDM with a single colony of *E. coli* BW25113 cells from a freshly streaked LB/agar plate. This culture was incubated post-inoculation for 10 h at 37°C with shaking (220 rpm). Aliquots of this culture were used to inoculate two fresh 5 ml volumes of supplemented M9 CDM to a starting $OD_{600}$ value of 0.05. One of the two cultures was immediately charged with Kdo-azide to a final concentration of 4 mM, and both cultures were incubated for 10 h at 37°C with shaking (220 rpm). The $OD_{600}$ of the two cultures was measured (2.8 and 2.75, respectively) to check for equivalent rates of growth and to ensure both cultures had reached stationary phase. The exponential phase Kdo-azide charged culture was harvested via centrifugation and LPS was labelled with AF488-alkyne via CuAAC. The second culture was charged with 4 mM Kdo-azide and re-incubated for a further 10 h at 37°C with shaking (220 rpm). Cells were then harvested via centrifugation and the LPS was fluorescently labelled with AF488-alkyne via CuAAC using the standard protocol detailed earlier. Labelling efficiency was assessed via confocal fluorescence microscopy ensuring standard 488 nm laser powers and detector gain settings were employed to enable a quantitative comparison of the two cultures.

**OMV isolation and purification**

OMVs were isolated from the *E. coli* BW25113 strain cultured in nanoparticle-free supplemented M9 CDM. A volume of nanoparticle-free supplemented M9 CDM was inoculated using *washed* bacterial cells obtained from a stationary phase culture grown in the same nanoparticle-free medium. The bacterial cells were first isolated from the stationary phase culture by centrifugation at 3400 xg for 20 min at 20°C using a temperature-controlled bench-top centrifuge. After centrifugation, the supernatant was carefully removed under sterile conditions. The cell pellet was resuspended in 10 ml of room temperature, nanoparticle-free supplemented M9 CDM and the cell suspension was centrifuged as above. After centrifugation, the supernatant was removed, and the resulting bacterial cell pellet was resuspended as above. This wash step was repeated once more, and the final bacterial cell pellet was resuspended in 3 ml of nanoparticle-free supplemented M9 CDM. This cell suspension was used to inoculate a 100 ml volume of nanoparticle-free supplemented M9 CDM to an $OD_{600}$ of ~0.1 and the inoculated culture was incubated at 37°C with shaking (165 rpm). An exact volume (5-6 ml) was removed from the bacterial culture for the purification of OMVs at designated time points. The volume removed was immediately placed on wet ice to suspend bacterial growth and then stored for no longer than 24 h at 4°C prior to OMV purification.

Bacterial cells were removed from the OMV-containing culture volume by centrifuging at 5000 xg for 20 min at 4°C. The supernatant containing the OMVs was carefully removed without disturbing the cell pellet after each centrifugation step. After centrifuging twice in this manner, the OMV-containing supernatant was immediately passed through 0.45 µm (Filtropur S, Sarstedt) and 0.22 µm (Appleton Woods) pore size hydrophilic (PES) syringe filters. Aliquots (10-20 µl) of the purified OMVs were spotted onto LB-Miller agar plates and the plates were incubated at 37°C for 24 h to confirm the absence of intact, viable bacteria. The purified OMVs were stored at 4°C prior to measuring their concentration by nanoparticle tracking analysis (NTA).

**Nanoparticle tracking analysis (NTA) of OMVs**

NTA was done using a ZetaView® Quatt instrument (Particle Metrix) within 48 h of purifying the OMVs from bacterial culture. All OMV samples were centrifuged at 21000 xg at 4°C in a 2 ml micro-centrifuge tube for 30 min just prior to NTA, then transferred to a fresh micro-centrifuge tube and allowed to reach room temperature (21°C) before the measurement. Ideal OMV concentrations for NTA (yielding 50-150 particles/frame) were determined by pre-testing each OMV sample after dilution with nanoparticle-free unsupplemented M9 CDM (with < 10 particles/frame) to a final volume of 1 ml. The NTA instrument was calibrated daily using certified 100 nm diameter polymer nanospheres (NanoStandards, Applied Microspheres). Triplicate technical replicates were measured for each OMV sample. For each sample measurement, the cell was scanned at 11 different positions while capturing 60 frames of video at each position (with 488 nm laser illumination, video frame rate = 30 frames per second, video setting = high, shutter = 100, camera sensitivity = 85).

After capture, the videos were analysed using the ZetaView® software (version 8.06.01 SP1) and standardised analysis parameters (*i.e.* minimum particle area = 10, maximum particle area = 1000, minimum particle brightness = 30, tracking radius = 100, minimum trace length = 15 frames). Only videos where 7 or more positions met the selection criteria were used to determine the OMV concentration. Particles up to 1000 nm in diameter could be tracked using this NTA procedure; however, we routinely observed that > 90% of the purified OMVs had a diameter ≤ 200 nm. Non-linear regression of the skewed OMV size distribution for each set of technical replicates (n = 3) was done for particles with a diameter ≤ 200 nm using GraphPad Prism (v10.3.1) assuming a log-normal distribution of particle sizes.

**Determining the total viable cell count of bacterial cultures**

The concentration of viable bacterial cells was determined at each time point using serial dilutions of the culture and the plate count method. Colony forming units (CFU) were calculated for each time point and plotted against the measured $OD_{600}$ value. This yielded CFU values of $2.00 \times 10^8$ cells/ml/$OD_{600}$ for exponential phase culture and $3.44 \times 10^8$ cells/ml/$OD_{600}$ for stationary phase culture ***(Extended Data Table S23)***. Critically, the CFU value remained stable for stationary phase cultures (*i.e.* 6.5 to 25 h post-inoculation of the culture).

**Derivation of outer membrane buckling model for OMV formation**

To derive a general expression for the critical size of an OMV formed by membrane blebbing from an LPS-rich region in the bacterial OM, we assume these regions or patches are comprised solely of LPS and are pinned at their periphery to the cell wall by relatively immobile OMP-rich regions (***Fig. 7A***). These patches will tend to bulge outwards due to the intrinsic curvature of an asymmetric membrane bilayer with inner and outer leaflets comprised of phospholipid and LPS, respectively. However, the patches remain stable — resisting blebbing — due to the intrinsic bending stiffness of the OM and anchoring by Braun's lipoproteins, provided the characteristic radius of the LPS patch is below a critical value. For the isotropic case, the critical radius is given by

$$R_c = \sqrt{\frac{3\kappa_b}{2\sigma}}$$

where $\kappa_b$ is the membrane bending stiffness and $\sigma$ is the compressive stress in the membrane. The load bearing properties of the OM, including its ability to sustain compressive stress, have been demonstrated previously [31].When the patch radius exceeds $R_c$, the buckled state becomes unstable, leading to membrane blebbing. The equation for $R_c$ is consistent with the published literature [32]. However, using this expression to predict critical OMV size is challenging because the value of $\sigma$ is generally unknown.

It has been shown that during exponential phase growth, the OM bears little or no load, and consequently the value of $\sigma$ is relatively small [31]. Based on the equation for $R_c$, we would expect membrane blebbing to be a relatively rare event during early to mid-exponential phase growth, and this conclusion is consistent with our time-dependent measurements of OMV concentration for wild-type *E. coli* BW25113 cells (***Fig. 6F***). To understand the dynamics of $\sigma$ during stationary phase, we note that while the membrane area ($A_s$) anchored to the peptidoglycan cell wall by OmpA and Braun's lipoprotein remains constant, LPS continues to be inserted at approximately a constant rate. Continued insertion of LPS leads to a time-dependent increase in $\sigma$:

$$\sigma(t) = \kappa_a \frac{Qt}{A_s}$$

where $\kappa_a$ is the membrane stretch modulus, *Q* is the inserted membrane area flux, and *t* is time.

The characteristic radius of the blebbing membrane ($R_b$) in an LPS patch is obtained by evaluating the equation for $R_c$ using a time-dependent value of $\sigma$ corresponding to a characteristic membrane blebbing time ($t_b$):

$$R_b = R_c\big(\sigma(t_b)\big).$$

After a membrane blebbing event, the LPS patch is removed from the OM which leads to relaxation of the compressive stress. Afterwards, the continued insertion of new LPS leads to a gradual build-up of compressive stress as described by the equation for $\sigma(t)$ until another membrane blebbing event in an LPS patch. In a steady state, the rate of LPS insertion must balance the rate of membrane area loss due to membrane blebbing and OMV release. This steady state condition yields the following:

$$Q = \frac{\pi R_b^2}{t_b}.$$

Solving the system of equations $R_c$, $R_b$ and *Q* leads to the following expression for the characteristic radius of the blebbing membrane:

$$R_b = \sqrt{w\, l},$$

where

$$w = \sqrt{\frac{3\kappa_b}{2\kappa_a}} \quad \text{and} \quad l = \sqrt{\frac{A_s}{\pi}}.$$

Remarkably, the resulting equation for $R_b$ is independent of Q, and this fortuitous circumstance makes this scaling law robust against the natural variability of LPS insertion rates observed *in vivo*. Since the diameter (*d*) of a vesicle formed from a single LPS patch is equal to $R_b$, we can predict the critical OMV size with the following expression:

$$d = \sqrt{w\, l}.$$

During late exponential phase and stationary phase growth, asymmetric expansion of the OM relative to the cell wall would arise and lead to a time-dependent increase in $\sigma$. In a steady state, the rate of LPS insertion must balance the rate of OM area loss due to blebbing. The radius of a blebbing LPS patch ($R_b$) at steady state was estimated using literature values of $\kappa_a$ (= 0.03-0.24 N/m [33]) and our experimentally determined value of $A_s$ (= 12.6 μm$^2$ for a typical cell size). Estimates of $\kappa_b$ and $\kappa_a$ were made by fitting atomic force microscopy measurements of membrane stiffness to a thin-shell membrane model [31,34] where the characteristic length scale ($w$) is related to the shell thickness ($h$) by $w = h/2^{3/2}$ [35]. The characteristic diameter ($d$) of the OMV formed from the membrane bleb would be equal to $R_b$. We obtain $d$ = 73-122 nm for an OMV released by this blebbing mechanism assuming $h$ is equivalent to the OM thickness and using a range of experimentally determined OM thicknesses (= 7.5-21 nm [36,37]) and $l$ = 2 μm as measured for a typical cell. This estimate of $d$ is in good agreement with our experimental results (***Extended Data Fig. S8B***, shaded region) considering the simplicity of the model and the absence of fitting parameters.

**Computational modelling of OM organisation, turnover, and OMV formation**

**Simulation domain, mask and coarse-graining**. We simulated a half-cell field (2.6 μm × 1.1 μm; total area, $A_{field}$ = 2.86 μm²) mapped to a 780 pixels x 330 pixels grid, pixel size = 3.33 nm. A static allowed-area mask that covered a fraction, $f_{allow}$ = 75% of the field (allowed area, $A_{allow} = 0.75\ x\ 2.86 = 2.145$ μm²) to approximate the LPS-accessible outer leaflet [38-40]. The remaining 25% represented OMP-occupied, LPS-inaccessible regions ("voids") consistent with OMP islands and the confined lateral mobility of OMPs and LPS in the *E. coli* OM [3,15]. The number of simulated LPS molecules in the allowed area was held approximately constant at $N_{sim} = 8 \times 10^4$ per frame. This N is a coarse-grained representation of the biological copy number expected in this field. In the insertion-trapping simulation (model 1), the allowed-area mask is static. In the phase-separation simulation (Model 2), the mask undergoes a gentle, area-preserving annealing at discrete 10 min intervals that increases its correlation length and produces the slow coalescence of voids, simulating the slow OMP-domain merging expected in a membrane system where the evolution of component organisation is driven principally by phase separation [41-43].

To relate simulated counts to an estimate of the numbers of LPS molecules within $A_{allow}$, we estimated the area density of LPS as:

$$\rho_{LPS} \approx \frac{N_{LPS}}{f_{LPS}\, A_{OM}}$$

$N_{LPS}$: Total number of LPS molecules per cell. $N_{LPS} \in [1.4, 2.0] \times 10^6$ molecules per cell [39,40,44,45]

$A_{OM}$: Outer membrane surface area per cell (units: μm$^2$) $A_{OM} = 4.31$ μm$^2$ [46]

$\rho_{LPS}$: Area density of LPS molecules (units: LPS μm$^{-2}$)

$f_{LPS}$: Fraction of the OM surface occupied by LPS. $f_{LPS} = 0.75$ [38-40]

Numerically, this yields

$$\rho_{LPS} \in [4.33, 6.19] \times 10^5 \text{ LPS } \mu\text{m}^{-2}$$

The simulation field is

$$A_{field} = 2.6 \ \mu m \times 1.1 \ \mu m = 2.86 \ \mu m^2$$

The allowed (LPS-accessible) area is

$$A_{allow} = f_{allow} A_{field} = 0.75 \ x \ 2.86 = 2.145 \ \mu m^2$$

The expected number of LPS molecules within $A_{allow}$ is therefore

$$\langle N_{field} \rangle = \rho_{LPS} A_{allow} \Rightarrow \langle N_{field} \rangle \in [9.29, \ 13.27] \times 10^5 \text{ LPS}$$

Given the number of simulated particles restricted to the allowed area $N_{sim} = 8 \times 10^4$, the corresponding coarse-graining factor, $\gamma$ relative to the biological count is

$$\gamma = \frac{\langle N_{field} \rangle}{N_{sim}} \in [11.6, 16.6]$$

thus one simulated LPS particle represents between 12 – 17 LPS molecules.

**LPS turnover kinetics (common parameters for both models).** Simulations advanced in discrete steps of Δ*t* = 5 min from *t* = 0 to 120 min (plots / .CSVs were exported every 10 min).

Background ("old") LPS decayed with per-step probability where:

$$p_{decay} = 1 - 2^{-\Delta t / t_{1/2}}$$

with $t_{1/2}$ = 90 min, the single-exponential half-life obtained by fitting the decline of old-LPS coverage in the pulse–chase dSTORM experiments ***(Fig. 2E–2F, Fig. 3)*** and consistent with the average rate of background LPS loss from the OM of cells over the first two hours ***(Fig. 6D-6E)***.

The fraction of newly inserted LPS increases linearly to ~0.65 at 120 min, reproducing the measured composition shift (reciprocal rise of newly inserted LPS vs decline of background LPS) observed experimentally ***(Fig. 2E–2F, Extended Data Tables S16 and S17)***. To minimise discretisation artefacts, we implement this as a smooth, late-weighted ramp specifying a target newly inserted LPS fraction at each step (not an absolute number).

Total LPS coverage in the field is held approximately constant at the target $N_{sim} = 8 \times 10^4$ simulated particles (background + newly inserted) across the allowed area. At each 5-min step, we applied updates in the following order:

1. Background LPS decay (using $p_{decay}$) and OMV-associated removals.
2. Computation of the target newly inserted LPS fraction.
3. New LPS insertion up to a per-step cap so that the running composition follows the experimental trajectory and the combined count respects the rule

$$(\textit{Background LPS} + \textit{Newly inserted LPS}) \leq N_{sim}$$

When removals exceeded insertions in a given step, the total number of particles was allowed to transiently fall slightly below $N_{sim}$ and was not renormalised upward.

**Burst-like new LPS insertion and restricted lateral diffusion of LPS (common settings for both models).** Within the allowed area, we pre-sampled 300 candidate insertion hotspots and, at each 5-min simulation step, randomly activated 110–150 sites. Newly synthesised LPS was inserted as compact two-dimensional Gaussian deposits (typical σ ≈ 20 - 40 nm, with light ellipticity jitter), consistent with discrete LptDE-linked delivery to the outer leaflet [12,47,48].

Between updates, both LPS particle types attempted highly restricted lateral displacements (σ ≈ 40 nm per 5 min) in-line with experimental observations that have shown LPS in *E. coli* BW25113 undergoes tightly restricted lateral diffusion [15,49]. Any LPS particle proposed movements that resulted in their entry into voids or crossing boundaries were rejected, reflecting the very low lateral mobility in the outer leaflet of the OM and confinement by OMP assemblies [3,15,49].

Diffusive step sizes were matched across both models (per-step standard deviation on the order of a few nm at 5 min cadence); only Model 2 adds an interaction-driven drift term.

Since turnover, insertion, lateral diffusion and OMV biogenesis were identical in both the insertion-trapping and phase separation models, the observed variations in newly inserted versus background LPS distributions arose from differences in the principles underpinning organisation, i.e. insertion-trapping versus phase separation.

**OMV biogenesis via OM buckling (release of OM compressive stress by vesiculation).** We modelled OMV biogenesis identically in both models, as a means of releasing compressive stress in LPS-rich patches whose rims are pinned by OM–peptidoglycan (PG) anchoring (for example, by Braun's lipoprotein), and where patch centers are comparatively weakly anchored ***(Fig. 7A)***. Continued new LPS insertion elevates in-plane compressive stress until a patch bulges and blebs to release an OMV. At each 5-min step, we drew a Poisson number of events corresponding to ~3 per 30 min on average and placed them at peaks of total LPS density (top 10%) at locations away from OMP 'voids'. Each event removed only a fraction of the local LPS within a radius of 80-90 nm footprint and re-packed remaining LPS particles with a small jitter (re-packing jitter ~20 nm), so no persistent hole remained thereby mimicking OMV 'pinching' ***(Fig. 7A)*** and membrane reshaping, rather than perforation. This implementation aligns with the load-bearing role of the OM and OM–PG tethering mechanics [31,50] along with the observed hyper-vesiculating phenotypes observed in *E. coli* mutant strains (Δ*tol-pal* and Δ*lpp)* with weakened envelope tethers [51]. The white 'preview' circles used in the figures are visual markers only and do not reflect OMV dimensions.

**Model 1 - Insertion-trapping (no phase separation).** Model 1 enforced persistent segregation of background and newly inserted LPS by application of two local spatial rules while maintaining identical global insertion and turnover kinetics to model 2.

1. **Newly inserted vs background (old) LPS avoidance**. At active hotspots, candidate new LPS particle insertions were accepted only where the smoothed OLD density is below a fixed threshold. This biases NEW to fill OLD-sparse patches, producing spatial anti-coincidence without altering the total number of new LPS particles inserted.
2. **Local bias of background LPS turnover**. Background LPS removal was locally increased in regions already enriched for new LPS insertion (probability scaled by the local unit-normalised new LPS particle density) and globally normalised at each step so that the population half-life of LPS particles remained at 90 min.

To emulate the compressive stress at highly active insertion sites without creating artefacts, recent newly inserted LPS 'bursts' nudge background LPS slightly outward within a 50-60 nm neighbourhood in small, isotropic displacements, preserving local LPS packing density in background LPS-rich patches. The Python script (Python_code_insertion_trapping_1.txt) and associated READ.ME_Python_code_insertion_trapping_1.txt) are provided as Supplementary Software.

**Model 2 - Phase separation.** Model 2 incorporated the same LPS insertion and turnover schedule, confined lateral diffusion and OMV biogenesis module as model 1 but replaced the insertion-trapping rules with like-like interactions and slow OMP void coalescence.

1. **Like-like attraction (mixed LPS field).** At each step, we computed smooth density fields and added a weak drift up the gradient of each particle's own density (Metropolis/Cahn–Hilliard-style), increasing co-locating (co-clustering) and promoting domain coarsening whilst conserving particle numbers. No explicit newly inserted versus background LPS avoidance rules were applied. and
2. **Void coalescence.** The allowed-area mask is lightly annealed at each 5 min step (diffuse-threshold cycle) to increase the mask's correlation length whilst preserving $f_{allow}$, yielding the slow merging of OMP-rich island 'voids' expected in a membrane system undergoing demixing[52-55].

Since all global kinetics matched those applied in model 1, the distinctive signatures of this model (increasing DICE coefficient and $g_{NB}(0) > 0$) arise from its intrinsic phase-separation characteristics and not from differences in lateral diffusion, or LPS insertion and turnover rates. The Python script (Python_code_phase_seperation_model_2.txt) and associated READ.ME_Python_code_phase_seperation_m2.txt) are provided as Supplementary Software.

**Zero-lag spatial cross-covariance (excess overlap over random), $g_{NB}(0)$**

To quantify co-location of background and newly inserted LPS clusters independent of marginal coverage, we computed a zero-lag, coverage-normalised overlap statistic, $g_{NB}(0)$ on binary cluster masks using DBSCAN parameters equivalent to those used in the dSTORM analyses ***(Figs. 2E and 4, Extended Data Figs. S2E and S4)***.

**Mask construction.** For each time point in the simulation, background and newly inserted (x, y) particle co-ordinates (in nm) were clustered via the DBSCAN application (ε = 30 nm; minPts = 7; core points only). Only core points were used to define robust cluster interiors. These points were converted to the image grid (780 pixels x 330 pixels, 3.33 nm pixel$^{-1}$) on the allowed-area mask. This produced two Boolean masks $A$ (background) and $B$ (newly inserted) indicating pixels that belonged to the DBSCAN clusters. The DBSCAN parameters were fixed and held constant across models and simulation replicates (N = 5 for each model).

**Definition.** $|A|$ and $|B|$ are the number of true pixels in the allowed area of the OM, and $|A \wedge B|$ is the count of the co-occupied pixels. $N$ is the number of allowed pixels.

$$p_A = \frac{|A|}{N} \qquad p_B = \frac{|B|}{N} \qquad p_{AB} = \frac{|A \wedge B|}{N}$$

The relative-to-random co-location is given by

$$g_{NB}(0) = p_{AB} - p_A p_B$$

This subtracts the expected random overlap $p_A p_B$ from the observed overlap. Thus $g_{NB}(0)$ is dimensionless and interpretable with $g_{NB}(0) > 0$ indicating greater co-location compared to random, $g_{NB}(0) = 0$ indicating random localisation, and $g_{NB}(0) < 0$ indicating anti-association with less co-location compared to random. $g_{NB}(0)$ values were computed per time point for each run (N = 5) in both simulations, and mean run values with standard errors were calculated and reported ***(Fig. 7D)***. Edge cases (with no clusters in *A* or *B*) yielding undefined overlap were marked *NaN* and excluded from reported summaries.

**Co-clustering metric (DICE coefficient)**

As a complementary, bounded similarity index, the DICE coefficient between background and newly inserted LPS particle cluster masks was computed at each time point in every run for both simulations. Masks were constructed via DBSCAN (ε = 30 nm; minPts = 7; core points only), with identical $g_{NB}(0)$ settings.

**Definition**

$$DICE(A, B) = \frac{2|A \wedge B|}{|A| + |B|}$$

The resulting unitless DICE coefficient value range was [0,1] where 0 indicates disjointed masks, and 1 indicates identical masks (perfect colocalisation of background and newly inserted LPS clustered pixels). If $|A| + |B| = 0$ (no clusters detected in either channel) then the DICE coefficient was undefined and recorded as *NaN* and excluded from downstream mean calculations.

**Interpretation.** Dice is sensitive to the spatial overlap of DBSCAN-defined cluster cores (i.e., the densest cluster pixels)and penalises cases where clusters are present in one channel but absent in the other channel for the same region. Outputs from our insertion-trapping simulation produced lower DICE values ***(Fig. 7C)***, resulting from spatial segregation of newly inserted and background LPS ***(Fig. 7B)***, compared to the equivalent time point outputs from the phase-separation simulation. The phase-separation simulation yielded higher, persistent DICE values ***(Fig. 7C)*** indicative of background and newly inserted LPS domain coarsening ***(Fig. 7B)***. We reported DICE means ± SEM across n = 5 independent runs per model at each time point.
